# Efficient detection and characterization of targets of natural selection using transfer learning

**DOI:** 10.1101/2025.03.05.641710

**Authors:** Sandipan Paul Arnab, Andre Luiz Campelo dos Santos, Matteo Fumagalli, Michael DeGiorgio

## Abstract

Natural selection leaves detectable patterns of altered spatial diversity within genomes, and identifying affected regions is crucial for understanding species evolution. Recently, machine learning approaches applied to raw population genomic data have been developed to uncover these adaptive signatures. Convolutional neural networks (CNNs) are particularly effective for this task, as they handle large data arrays while maintaining element correlations. However, shallow CNNs may miss complex patterns due to their limited capacity, while deep CNNs can capture these patterns but require extensive data and computational power. Transfer learning addresses these challenges by utilizing a deep CNN pre-trained on a large dataset as a feature extraction tool for downstream classification and evolutionary parameter prediction. This approach reduces extensive training data generation requirements and computational needs while maintaining high performance. In this study, we developed *TrIdent*, a tool that uses transfer learning to enhance detection of adaptive genomic regions from image representations of multilocus variation. We evaluated *TrIdent* across various genetic, demographic, and adaptive settings, in addition to unphased data and other confounding factors. *TrIdent* demonstrated improved detection of adaptive regions compared to recent methods using similar data representations. We further explored model interpretability through class activation maps and adapted *TrIdent* to infer selection parameters for identified adaptive candidates. Using whole-genome haplotype data from European and African populations, *TrIdent* effectively recapitulated known sweep candidates and identified novel cancer, and other disease-associated genes as potential sweeps.

## Introduction

The study of biological diversity and adaptation relies on the identification and understanding of past evolutionary events. These events shed light on the processes that have shaped the genetics of organisms throughout history and those that are ongoing. Identifying genomic regions that are subject to the pressures of natural selection enables the discovery and isolation of genetic variants that may have contributed to adaptive traits in past environments. Understanding the genetic basis of vital characteristics like disease resistance, reproductive success, and physiological adaptations to changing environmental conditions illuminates the evolutionary history of a species. Predicting how species will respond to shifting weather patterns [Hoffmann and Sgrò, 2011], altered habitats [Grant and Grant, 2002], and the emergence of new diseases [Allison, 1954] is enhanced by these findings. A window into the complex dynamics of evolution and adaptation can be found in the discovery of evolutionary events in genomes, which has implications for both theoretical and applied research.

A number of approaches have been taken to uncover traces of natural selection from polymorphisms within a population in light of neutral evolutionary processes, such as mutation, recombination, migration, and genetic drift, which can both bolster and erode past adaptive signals [Kimura, 1979, Slatkin, 1987, Barton and Charlesworth, 1998, Lynch, 2010]. Initial efforts for detecting such non-neutral regions involved summary statistics based on alterations of the distribution of allele frequencies, such as Tajima’s *D* [Tajima, 1989] and Fay and Wu’s *H* [Fay and Wu, 2003]. These summary statistic approaches have been enhanced in contemporary studies to evaluate distortions in haplotype frequency distributions, such as H12 [Garud et al., 2015], and to assess the footprint of reduced genomic diversity through extended haplotype homozygosity methods, such as EHH [Sabeti et al., 2002], iHS [Voight et al., 2006], and *nS_L_* [Ferrer-Admetlla et al., 2014]. Further, more model-based approaches founded in population genetic theory or sensible probability distributions have revealed improved power, ability to isolate loci at which selection has likely acted, and effectiveness at estimating underlying parameters of the adaptive process [Nielsen et al., 2005, Stephens and Balding, 2009, Chen et al., 2010, Pasaniuc et al., 2014, Vy and Kim, 2015, Racimo, 2016, Lee and Coop, 2017, Lloyd-Jones et al., 2019, Harris and DeGiorgio, 2020, Setter et al., 2020, DeGiorgio and Szpiech, 2022]. However, more recently, there has been a focus on a promising branch of selection detection methods based on machine learning or artificial intelligence that consider evidence from multiple input statistics or that operate on raw haplotype alignments or genotype calls [Schrider and Kern, 2018, Korfmann et al., 2023].

Supervised machine learning represents an important branch of such methods, where models are trained on a set of observations often composed of a multidimensional input and a paired output value [Hastie et al., 2009]. Training algorithms learn patterns and the functional relationship between the input and output data from these example observations (termed the training set) to hopefully make accurate output predictions on unseen future input data. Classification and regression tasks both make use of supervised learning algorithms to achieve their respective goals of organizing inputs into predefined categories (termed classes) and making continuous outcome predictions. There exist numerous supervised learning algorithms for classification and regression tasks, each with its own assumptions about data distributions and functional relationships (*e.g.*, degree of nonlinearity) between the input and output values. Of course, no one approach is best, and the prediction abilities of the trained models highly depend on the correctness of the assumed data distribution and true underlying function relating the input and output. Artificial neural networks are universal approximators [Hornik, 1991], having the capacity to represent any underlying function with enough model parameters, and can thus learn complicated patterns in a manner reminiscent of how the human brain works. One recent type of neural network that has enjoyed extensive success in image classification and recognition tasks is the convolutional neural network (CNN) [Krizhevsky, 2012], which is optimized for handling structured grid-like data.

Deployment of advanced machine learning and artificial intelligence tools has brought about a revolution in the realm of natural selection pattern recognition. Some of the applications of machine learning include the extraction of features from one or many statistics computed in contiguous genomic windows, significantly increasing power to detect patterns compared to the classical usage of such statistics. Many of the state-of-the-art applications in this paradigm make use of powerful contemporary feature extraction tools on summary statistic arrays, ranging from simple linear to complex nonlinear models. Nevertheless, massive volumes of training data are typically necessary for nonlinear models, such as neural networks, to learn their parameters due to high model complexity. Such models are prone to overfitting and underperformance in the absence of large training sets, or through the use of automated frameworks to reduce the capacity or representation ability of the learned models [Hastie et al., 2009].

While data availability can be a limiting factor in some fields, population genetics benefits from the ability to generate effectively unlimited training data via simulation, particularly with simulation-on-the-fly techniques that provide new data with each iteration [Chan et al., 2018, Torada et al., 2019, Flagel et al., 2019, Battey et al., 2020], thus mitigating overfitting. However, minimizing the need for extensive training data remains beneficial to reduce computational demands and environmental impact, as highlighted by recent studies on the carbon footprint of bioinformatics tools [Grealey et al., 2022]. Efficient and adaptable methods are crucial for analyzing genomic patterns shaped by natural selection. A framework that can swiftly summarize and capture these patterns without relying on application-specific summary statistics offers significant advantages. Utilizing a robust feature extractor capable of handling generalized genomic summaries ensures both adaptability and effectiveness, particularly in contexts with limited computational resources. This approach enhances the ability to detect and analyze complex genomic patterns, balancing the need for efficiency with sophisticated analysis.

This challenge can be addressed by applying transfer learning, a method of pre-training a neural network on unrelated data [Bozinovski, 2020]. Pre-trained models are used in image classification to speed up convergence and improve outcomes [Hendrycks et al., 2019]. The field of computer vision has been significantly enhanced by models trained on the ImageNet dataset [Deng et al., 2009]. ImageNet is a massive collection of approximately 14 million annotated images organized into 1,000 categories that provide adequate diversity for developing powerful image categorization applications. Harnessing the diverse spectrum of images in this dataset and by leveraging high-performance computing resources, deep learning has made several prominent architectures the gold standard for image classification [Russakovsky et al., 2015, Beyer et al., 2020]. A transfer learning model trained on a massive set of labeled images can be used as a feature extraction tool, thus eliminating the need to train complex models to succinctly represent input genomic data in a transformed space that is usable for downstream prediction tasks. Various transfer learning architectures have been successfully deployed in non-natural image categorization tasks, such as medical imaging or industrial defect detection [Ming et al., 2021, Suganyadevi et al., 2022, Shafique et al., 2022], which contrasts with natural images that typically capture scenes from the natural world (*i.e.*, landscapes or wildlife). This distinction underscores the robustness of these architectures, alleviating concerns about substandard performance when classifying images that capture patterns left by evolutionary events.

Instead of constructing a shallow CNN that operates on input images [Zhao et al., 2023] from a dataset with limited numbers of samples, it may be better to use pre-trained deep CNN architectures. Unlike shallow CNNs, which are unable to capture the complex relationship between input images and the category they belong to, these pre-trained deep CNNs are able to significantly improve the ability to model such complex relationships. Another advantage of pre-trained CNNs is their capacity to generalize learning from one domain to another [Yosinski et al., 2014, Sharif Raza-vian et al., 2014]. This ability helps pre-trained CNNs perform better than shallow custom-made CNNs constructed and trained using smaller datasets [Azizpour et al., 2015, Tajbakhsh et al., 2016]. Feature extraction from small datasets is enhanced by pre-trained CNN architectures, leading to better generalization performance and resilience in tasks that have few data samples. Because training deep CNNs from scratch requires substantial computational resources, making use of pre-trained models becomes a more feasible and efficient option.

In this article, we aim to develop powerful and robust natural selection detection and characterization tools by exploiting the feature extraction efficacy afforded by pre-trained CNN models. Specifically, we apply five neural networks architectures pre-trained on the large ImageNet database, which we use to extract transformed feature sets for input to classifiers to detect adaptive events and to regressors to predict parameters of such events. By training models with a small number of simulated replicates and then testing their predictions with an array of confounding factors, we demonstrate that transfer learning can be a reliable tool for building predictive population genomic models that take images of haplotype variation as input without requiring extensive training data. Moreover, we introduce an efficient approach for constructing input images of haplotype variation that organizes diversity in a way that is easier for downstream image-based sweep classifiers to extract key patterns, ultimately improving their accuracies and powers. Here, we present *TrIdent* (*Tr* ansfer learning for *Ident* ification of adaptation), which is implemented as open-source software available at https://www.github.com/sandipanpaul06/TrIdent. As an empirical case study, we apply *TrIdent* to haplotype variation from a well-studied European human population (CEU) from the 1000 Genomes Project dataset, and are able to recapitulate established selective sweep candidates, such as *LCT* and *OCA2*, and discover candidate sweep signals in novel cancer-associated genes, such as *MTOR*, *LAMC2*, *RBMS3*, and *NKAIN2*, with high support. Similarly, when applied to a well-studied African human population (YRI) from the same dataset, *TrIdent* successfully identifies established sweep candidates, such as *HEMGN*, *SYT1*, *GRIK5*, *FOXP2* and *APOL1*, while also uncovering novel candidate sweep signals in *ROBO2*, with strong support.

## Results

### Modeling description

We used the coalescent simulator discoal [Kern and Schrider, 2016] under two nonequilibrium demographic histories estimated for European (CEU) and sub-Saharan African (YRI) humans [Tennessen et al., 2012], which respectively include a recent severe population bottleneck and a recent population expansion, to simulate neutral and sweep replicates to train and evaluate *TrIdent*. We simulated sweeps with per-generation selection strengths (*s*) ranging from 0.005 to 0.1, beneficial allele frequencies (*f*) from 1*/*(2*N_e_*) to 0.2, and fixation times of beneficial alleles (*τ*) from 0 to 2,000 generations prior to sampling. Moreover, sweep and neutral replicates were generated with per-site per-generation mutation rates (*µ*) from 2.21 × 10*^−^*^9^ to 2.21 × 10*^−^*^8^, and per-site per-generation recombination rates (*r*) from 0 to 3 × 10*^−^*^8^ with a mean rate of 10*^−^*^8^ (see *Simulation protocol* sub-section of the *Methods*). Together, these settings and parameters allowed for the examination of a broad range genetic and adaptive conditions, while focusing on two distinct demographic histories. Given the wide range of selection strengths and beneficial allele frequencies, along with varying mutation and recombination rates and fixation events that may have occurred far in the past, and further complicated by fluctuations in population size, substantial overlap in the distribution of genetic variation between neutral and selection settings is expected. This overlap creates significant challenges in distinguishing the sometimes subtle genetic patterns driven by positive selection from those that emerge under neutrality.

For the purpose of producing inputs to feed *TrIdent*, we formed the CEU and YRI training datasets by using 1,000 neutral and 1,000 sweep replicates that were simulated using their respective demographic histories. In addition, we generated two sets of test and validation datasets to complement the CEU and YRI training datasets, each consisting of 1,000 neutral and 1,000 sweep replicates. We detail the processing of input images from these simulated replicates in the *Image Generation* subsection of the *Methods*. The image generation process (Figure 1; top panel) transforms these replicates into grayscale images representing sorted minor allele counts across shifting small and overlapping windows, and then resized to match the input expectations of the pre-trained models.

**Figure 1:**
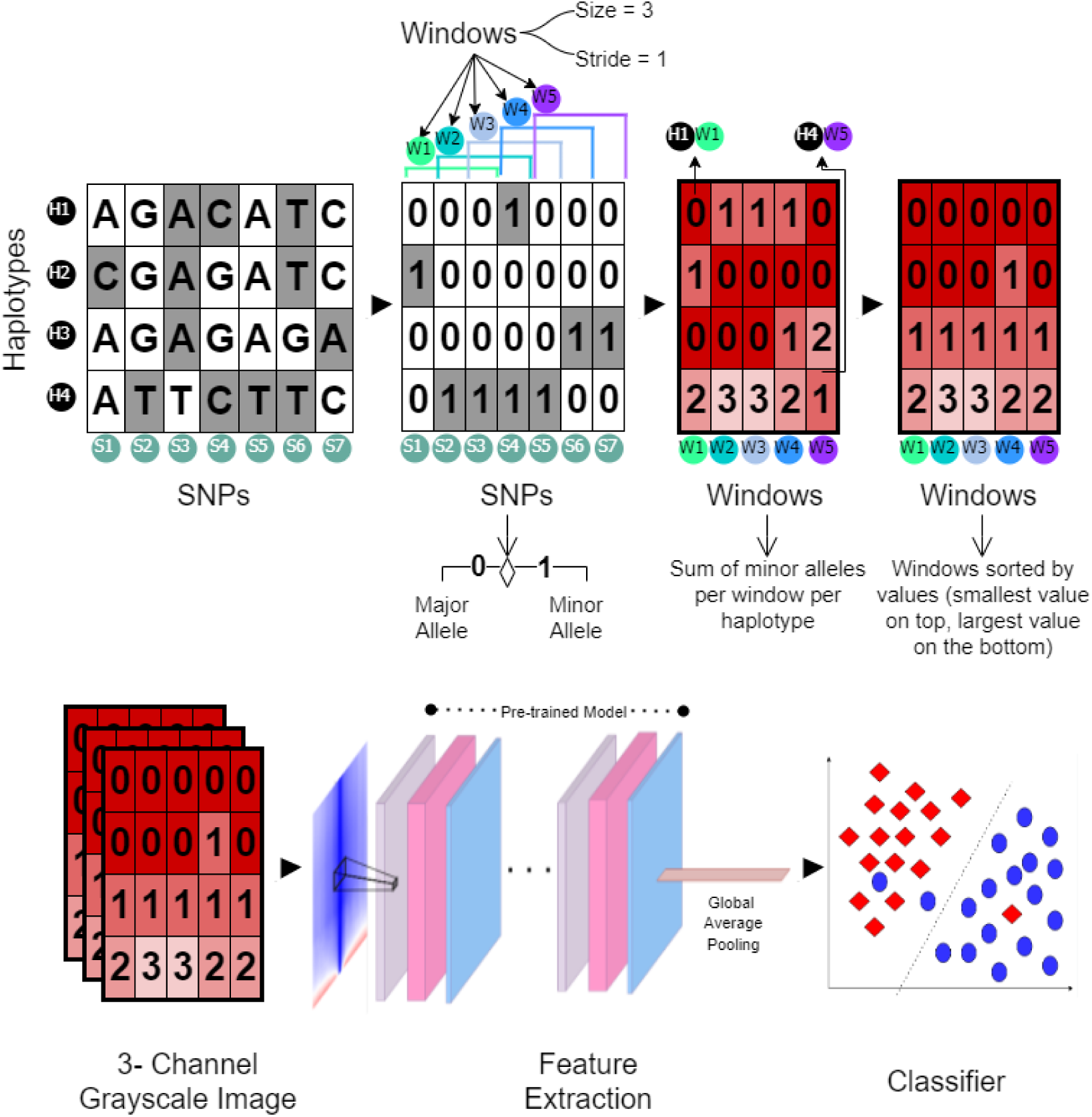
Depiction of the *TrIdent* [*IRV2*] model (bottom panel), including the native *TrIdent* image generation method (top panel). As described in *Image generation* subsection of the *Methods*, for a haplotype alignment with haplotypes on rows and SNPs on columns, the major allele is represented by zero and the minor allele by one at each SNP. From this processed alignment, the number of minor alleles for each haplotype is counted in a window and window locations are shifted by a specific stride, which is chosen as window size of three SNPs and a stride of one SNP in this schematic. The minor allele counts for each window are then sorted, such that the top row with have smallest value and the bottom row the largest value for a given window. A matrix based on a certain number of consecutive windows (here five windows) is created, this matrix is copied over two more channels to create a tensor, resulting in a three channel grayscale image. This image is fed as input to the “Feature Extraction” block consisting of a pre-trained deep convolutional neural network model that may incorporate various combinations (indicated by blocks of different colors) of a subset of the following layers: convolutional, maxpooling, dense, dropout, squeeze-and-excitation, depthwise separable convolutional, and residual connections. A global average pooling layer is attached to the pre-trained model to generate a feature vector, which is then used to train a classifier. The *TrIdent* [*IRV2*] model, which is focused on this article, combines the use of *InceptionResNetV2* as the pre-trained model and penalized logistic regression as the binary classifier.

Heatmaps depicting mean images for the sweep and neutral classes for the CEU and YRI datasets show that, in contrast to the neutral image, the sweep image features a dark vertical segment in the central columns representing a loss of diversity in the central columns due to high frequency haplotypes (Figure S1). That is, these regions contain a string of major alleles that are at high frequency near the center of the sweep replicates. Once these input images have been resized, each pixel is standardized according to its value at that particular position across all training images, so that each pixel has a mean of zero and a standard deviation of one. Heatmaps depicting mean standardized neutral and sweep images show that the neutral image has a red segment of positive values in the central columns to complement the blue segment of negative values in the sweep image (Figure 2). Thus, standardization reveals a clear distinction between the two classes, and serves as a proof of concept that the applied image generation technique presents a pattern that can be employed to discriminate between positive selection and neutrality.

**Figure 2:**
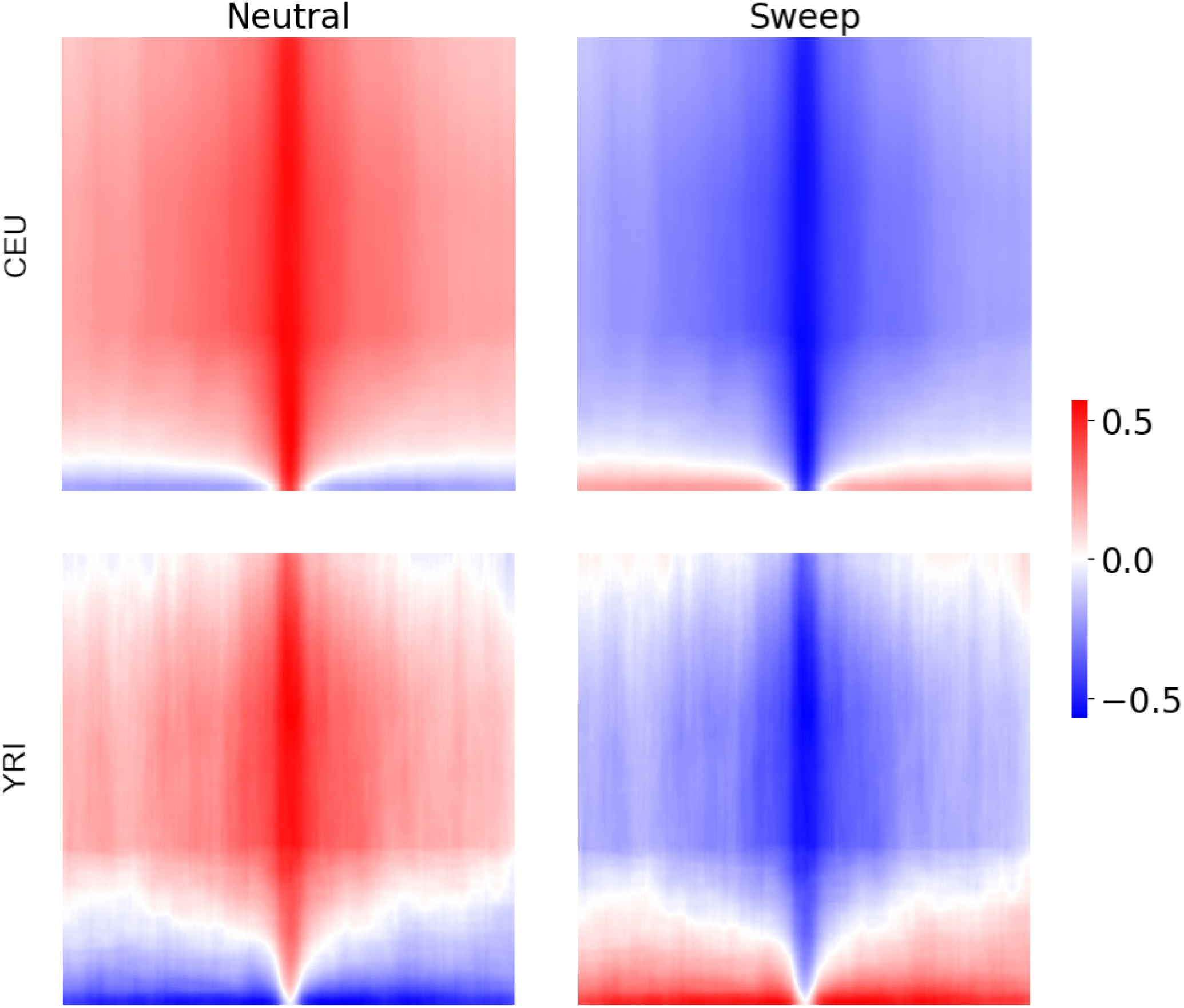
Heatmaps depicting standardized input images of size 224×224, averaged across the 1,000 training replicates for either the neutral or sweep class simulated under either the European (CEU) or Sub-Saharan African (YRI) human demographic history [Tennessen et al., 2012]. Standardized input images are processed as in the *Image generation* subsection of the *Methods*, with standardization occurring across the 2,000 neutral and sweep training replicates for each pixel. Rows of the images represent haplotypes, whereas columns represent genomic window of 25 contiguous SNPs within a haplotype, with an equal number of windows flanking the center of a simulated genomic region. The colorbar indicates a measure proportional to the number of minor alleles within the haplotype window relative to the mean number (scaled by the standard deviation of minor allele counts) for that window across the neutral and sweep training observations. Darker blue shading represents a higher number of major alleles than average and darker red shading represents a higher number of minor alleles than average.

These standardized images are fed to pre-trained models, after which we retain the output vector, representing transformed features or embeddings, from the global average pooling (GAP) layer that we attach to the pre-trained model (Figure 1; bottom panel). The GAP layer computes the mean value of each feature map, effectively reducing the spatial dimensions and summarizing the most salient features of the image. By averaging each feature map into a single node, GAP reduces the number of parameters in the model and makes the model more robust to spatial translations, offering advantages over traditional fully connected layers [Szegedy et al., 2015]. A penalized logistic regression classifier is then trained using the GAP layer outputs from 1,000 neutral and 1,000 sweep simulations, along with an elastic-net regularization penalty to simultaneously modulate model complexity and sparsity [Zou and Hastie, 2005, Hastie et al., 2009]. In particular, the magnitudes of model parameters associated with each transformed input feature are controlled by an *L*_2_-norm penalty, and feature selection is carried out to control sparsity by setting some model parameters to zero through an *L*_1_-norm penalty. The hyperparameter *α* ∈ {0.0, 0.1*, … ,* 1.0} specifies the proportion of model regularization that derives from the *L*_2_-norm penalty, whereas the hyperparameter *λ* ∈ {10*^−^*^6^, 10*^−^*^5^*, … ,* 10^5^} designates the amount of total regularization. We utilized the LogisticRegression module from the *Scikit-learn* [Pedregosa et al., 2011] package in Python to train the classifier. The best pair of hyperparameters to train the classifier was determined based on binary cross-entropy loss across the validation set.

### Choosing an appropriate pre-trained model

Numerous pre-trained models are available for transfer learning, including *VGGNet* s, *GoogleNet* s, *ResNet* s, and *EfficientNet* s that each have their own architectures and distinguishing features. A minimalist architecture and a modest number of convolutional filters distinguish the Visual Geometry Group (*VGG*)*Net* from other models pre-trained on the ImageNet dataset [Simonyan and Zisserman, 2014]. Szegedy et al. [2015] found that the Inception modules of *GoogleNet* improved performance and allowed it to capture features at multiple scales, whereas *ResNet* mitigated the vanishing gradient problem, a hurdle in training extremely deep networks consisting of multiple layers designed to learn complex hierarchical representations of the input data by focusing on residual learning, which helps information flow more smoothly through the layers via skipped connections between layers [He et al., 2016]. Recent developments like *EfficientNet* and *MobileNet* employ a compound scaling parameter, through which the numbers of layers and nodes in a layer are controlled. Capacity to modulate the depths of layers and width of nodes allows these models to offer a trade-off between efficiency and complexity [Tan and Le, 2019]. The *InceptionResNet* architecture is an augmentation of the *Inception* architecture by incorporation of residual learning [Szegedy et al., 2017], and represents a robust image recognition system with high accuracy in numerous tasks. By adding residual connections to its Inception blocks, *InceptionResNetV2* improves network information flow and gradient propagation during training [Wang et al., 2021]. We first study the efficacy of five pre-trained models (*InceptionResNetV2*, *VGG16*, *EfficientNetB0*, *EfficientNetB7*, and *MobileNetV2*) as complex feature extraction methods to develop adaptive event classification tools. We accessed these models with TensorFlow [Abadi et al., 2015], as their APIs make communication with these pre-trained models as well as their integration into workflows easier.

We evaluated the accuracy and power of the *TrIdent* logistic regression classifiers developed based on GAP layer outputs attached to the pre-trained *InceptionResNetV2*, *VGG16*, *Efficient-NetB0*, *EfficientNetB7*, and *MobileNetV2* on both the CEU and YRI datasets. For convenience of presentation, we respectively refer to these models as *TrIdent* [*IRV2*], *TrIdent* [*VGG16*], *TrIdent* [*ENB0*], *TrIdent* [*ENB7*] and *TrIdent* [*MNV2*]. On the CEU dataset, we found that *TrIdent* [*IRV2*], with its high number of parameters and computational demand, achieves the highest classification accuracy of 86.2% along with the highest sweep detection accuracy among all five classifiers (Figure S2). In contrast, the most lightweight and computationally efficient *TrIdent* [*VGG16*] as well as the moderately accurate and moderately efficient architecture of *MobileNetV2* achieve identical accuracies of 85.1% (Figure S2). The two *EfficientNet* classifiers, *TrIdent* [*ENB0*] and *TrIdent* [*ENB7*], are the worst performers out of the five *TrIdent* models. However, *TrIdent* [*ENB0*], with an accuracy of 84.95%, outperforms *ENB7*, which achieves an accuracy of 83.8%. This finding suggests that the shallower architecture of *ENB0* performs better in this case, contrary to the expectation that deeper architectures typically yield higher accuracy. These results are mirrored by the powers of these methods to detect sweeps based on receiver operating characteristic (ROC) curves, with *TrIdent* [*IRV2*], *TrIdent* [*MNV2*], and *TrIdent* [*VGG16*] exhibiting similar ROC curves and *TrIdent* [*IRV2*] slightly edging out *TrIdent* [*MNV2*] and *TrIdent* [*VGG16*] (Figure S2). On the other hand, *TrIdent* [*ENB7*] showcases the lowest power to detect sweeps among the five classifiers (Figure S2).

Our results also show that the classification behavior of *TrIdent* on the YRI dataset relative to CEU depends on network architecture (compare Figures S2 and S3). Specifically, *TrIdent* [*IRV2*] and *TrIdent* [*MNV2*] lead the set of classifiers with overall accuracies of 87.55% and 87.50%, respectively, with *TrIdent* [*IRV2*] marginally edging out *TrIdent* [*MNV2*] (Figure S3), similarly to the application on the CEU dataset (Figure S2), with slightly higher accuracies than on the CEU dataset, which may be expected as the YRI dataset has higher mean neutral haplotype diversity. The ROC curves of this pair of classifiers also indicate their proximity in performance metrics, while clearly distancing themselves from the other three classifiers. Moreover, *TrIdent* [*ENB0*] and *TrIdent* [*ENB7*] again prove to be the weakest of the set of classifiers with accuracies of 74.35% and 77.30%, respectively (Figure S3). However, unexpectedly, their accuracies are significantly lower than on the CEU dataset. *TrIdent* [*VGG16*], similar to the CEU test case, stations itself in the middle of these pairs of classifiers with an overall accuracy of 84.55% (Figure S3). Following the superior classification performance of *TrIdent* [*IRV2*] compared to the other *TrIdent* models in both the CEU and YRI test cases, we elected to pursue *TrIdent* [*IRV2*] for future analyses. Given our simulation setup and the two classes analyzed, *TrIdent* [*IRV2*] proved to be the most effective and consistent model. However, it may not be the best choice for all applications. Other pre-trained models, including but not limited to the ones we tested, could perform better depending on the specific context and requirements.

### Viability of alternate architectures and methods

Though *TrIdent* [*IRV2*] uses a linear model (logistic regression classifier) with features extracted from the *InceptionResNetV2* architecture, we also evaluated nonlinear models based on artificial neural networks (ANNs) for classification using the same extracted features. We refer to this approach as *TrIdent* [*IRV2*, *ANN*]. We optimized the *TrIdent* [*IRV2*, *ANN*] architecture by varying the number of hidden layers *L* ∈ {1, 2, 3, 4, 5}, the number of nodes *n_ℓ_* ∈ {100, 200*, … ,* 1000} in each hidden layer *ℓ* ∈ {1, 2*, … , L*}, and the activation function for nodes in each hidden layer *ϕ_ℓ_* ∈ {ReLU, sigmoid, tanh}. The node in the output layer has a sigmoid activation function, and the input layer has 1,536 nodes, which is identical to the number of features generated by the GAP layer attached to the *InceptionResNetV2* architecture. Like our linear *TrIdent* [*IRV2*] model, *TrIdent* [*IRV2*, *ANN*] also employs *L*_1_- and *L*_2_-norm regularization penalties for each hidden and output layer, with regularization hyperparameters chosen from the same ranges as those for the linear model (see *Modeling description* subsection). Each model is trained to minimize binary cross-entropy loss using the Adam optimizer [Kingma and Ba, 2015]. The optimal architecture and hyperparameters were selected based on the smallest binary cross-entropy validation loss on the CEU dataset, which resulted in a model of *L* = 3 hidden layers with *n*_1_ = 1000, *n*_2_ = 500, and *n*_3_ = 500 nodes in the first, second, and third hidden layers, respectively, sigmoid activations for nodes in the first and second hidden layers, and a tanh activation for nodes in the third hidden layer. We obtained a near identical optimal architecture for the YRI dataset, with the exception of an additional hidden layer (*L* = 4) with *n*_4_ = 500 nodes and a tanh activation. However, when applied to the same test data used for the linear model, *TrIdent* [*IRV2*, *ANN*] classification performance was virtually unchanged relative to *TrIdent* [*IRV2*] (Figure S4). This result was consistent even with various training set sizes (1,000, 3,000, and 5,000 observations per class), suggesting that the nonlinearity introduced by the ANN did not lend additional benefit beyond that of the *InceptionResNetV2* feature extractor. We therefore decided to continue with the linear *TrIdent* model for classification tasks, as the simpler logistic regression model has fewer parameters and is faster to train than the ANN.

We also tested the classification performance of *TrIdent* [*IRV2*] against *T-REx* [Amin et al., 2023], which is a high performing tensor decomposition based sweep classifier that operates on a windowed aggregation of genetic diversity as its input, similar to our image generation method, and that uses an *L*_1_- and *L*_2_-norm penalized logistic regression classifier for prediction, which is identical to *TrIdent*. To provide a fair comparison of performance, we tested *TrIdent* [*IRV2*] and *T-REx* with both image generation techniques. That is, *TrIdent* [*IRV2*] and *T-REx* were trained and evaluated based on their own native image generation approaches, as well as the image generation approaches of the competing model. For clarity, we denote the *TrIdent* [*IRV2*] model trained and tested on the alternate image generation style of *T-REx* as *TrIdent* [*IRV2*, *alt*], and the *T-REx* model trained and tested on the alternate image generation style of *TrIdent* as *T-REx* [*alt*]. We find that on the CEU dataset, the *TrIdent* [*IRV2*, *alt*] alternate implementation loses 2% accuracy compared to the stock *TrIdent* [*IRV2*] model (Figure 3). Moreover, *T-REx* trained on its native image generation style exhibits lower classification accuracy (79.6%) compared to both the *TrIdent* [*IRV2*] models (Figure 3). However, the *T-REx* [*alt*] alternate implementation reports a higher classification accuracy (81.75%) compared to the stock *T-REx* model, but falls short of *TrIdent* [*IRV2*] by 4.45% (Figure 3). The powers of each of these models to detect sweeps at a 5% false positive rate (FPR), lend further support that *TrIdent* models significantly outperform the *T-REx* models, regardless of the input image representation used by *T-REx* (Figure 3). Specifically, *TrIdent* [*IRV2*] marginally surpasses *TrIdent* [*IRV2*, *alt*] as the most powerful model of the four at a 5% FPR.

**Figure 3:**
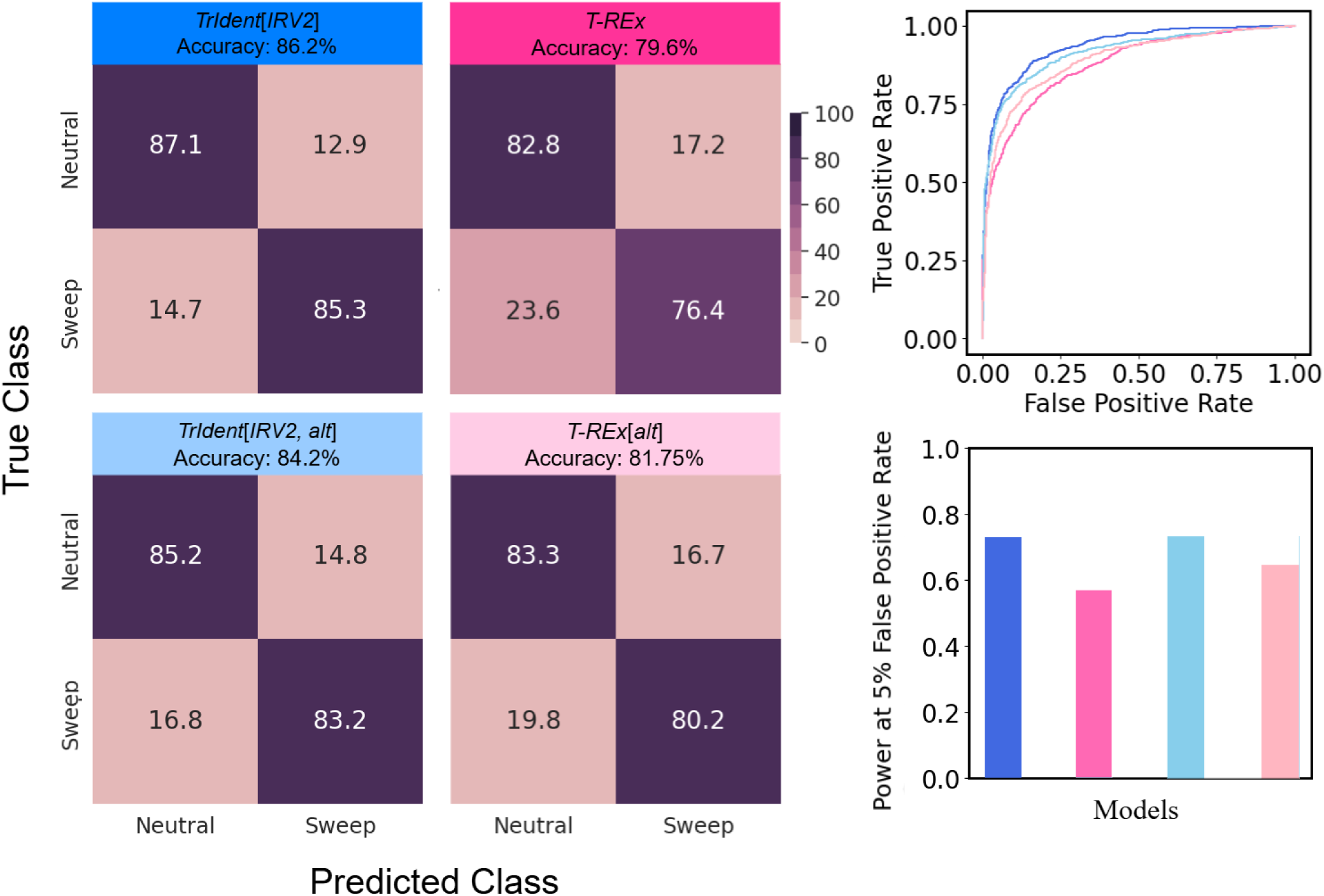
Classification rates and accuracies as depicted by confusion matrices, powers (true positive rates) to detect sweeps as depicted by receiver operating characteristic curves to differentiate sweeps from neutrality, and powers at a 5% false positive rate to detect sweeps on the CEU dataset for the best performing *TrIdent* model (*TrIdent* [*IRV2*]) compared to *T-REx*. The comparison also includes *TrIdent* [*IRV2*, *alt*] and *T-REx* [*alt*], which are trained and tested using their alternate image generation styles (see *Viability of alternate architectures and methods* subsection of the *Results* for details).

A parallel illustration is observed when the performances of the stock and alternate models of *TrIdent* [*IRV2*] and *T-REx* are evaluated on the YRI dataset (Figure 4). The stock *TrIdent* [*IRV2*] model leads the four models with an accuracy of 87.55%, followed by *TrIdent* [*IRV2*, *alt*] (85.30%), *T-REx* [*alt*] (83.85%), and *T-REx* (82.80%). The detection capability of *TrIdent* [*IRV2*] for identifying sweeps at a 5% FPR provides additional evidence that *TrIdent* [*IRV2*] is the superior model of the four in this test case (Figure 4). Though a deeper inspection of model performances shows that the performance difference seen among the models in the CEU test case (Figure 3) is noticeably reduced in the YRI test case (Figure 4), with the absence of a population bottleneck in the YRI dataset apparently bridging the performance gap among models.

**Figure 4:**
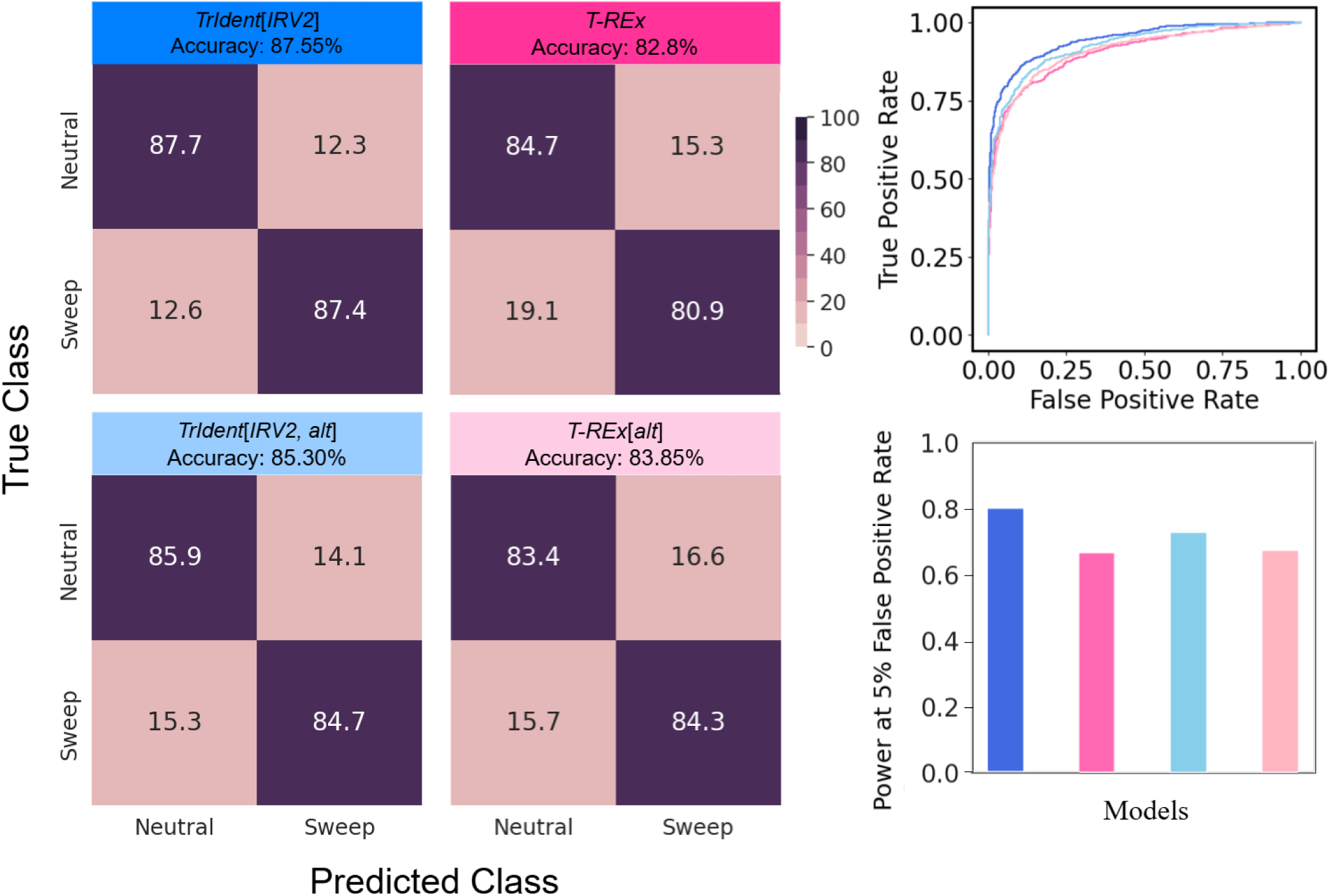
Classification rates and accuracies as depicted by confusion matrices, powers (true positive rates) to detect sweeps as depicted by receiver operating characteristic curves to differentiate sweeps from neutrality, and powers at a 5% false positive rate to detect sweeps on the YRI dataset for the best performing *TrIdent* model (*TrIdent* [*IRV2*]) compared to *T-REx*. The comparison also includes *TrIdent* [*IRV2*, *alt*] and *T-REx* [*alt*], which are trained and tested using their alternate image generation styles (see *Viability of alternate architectures and methods* subsection of the *Results* for details).

In addition to comparing *TrIdent* [*IRV2*] with *T-REx*, we conducted a comprehensive evaluation by also comparing it against diploS/HIC [Kern and Schrider, 2018], a state-of-the-art selective sweep detector that integrates summary statistics with comparatively shallow CNNs. We trained and tested diploS/HIC in its native setting, with one modification as we calculated the summary statistics in 101 windows instead of 11. While diploS/HIC typically uses 11 windows (resulting in 11 features for the downstream CNN model for each summary statistic), we chose to use 101 windows (resulting in 101 features for the downstream CNN model for each summary statistic), which retains the focal window but within an increased number of flanking windows. This adjustment resulted in a finer representation, and enhanced the prediction performance of diploS/HIC in our initial investigation. We also extended our analysis to evaluate a variant of the *TrIdent* [*IRV2*] architecture to better understand its performance when operating on summary statistic data, which we denote as *TrIdent* [*IRV2*, *SS*]. Specifically, we employed the 12 diploS/HIC summary statistics—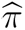 [Tajima, 1983], 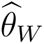[Watterson, 1975], Tajima’s *D* [Tajima, 1989], the variance, the skewness, and the kurtosis of multilocus genotype distances between pairs of sampled individuals, the multilocus genotype equivalents of expected haplotype homozygosity *H*_1_, *H*_12_ [Garud et al., 2015, Harris et al., 2018], and *H*_2_*/H*_1_ [Garud et al., 2015, Harris et al., 2018], unphased *Z_ns_* [Kelly, 1997, Rogers and Huff, 2009], and the maximum value of *ω* [Kim and Nielsen, 2004, Rogers and Huff, 2009]—which provide insights into multiple aspects of genetic diversity affected by selective sweeps, including the structure of haplotype variation, nucleotide diversity, and linkage disequilibrium. Together, these components offer a comprehensive view of the effects of selective sweeps on genomic regions [Schrider and Kern, 2016, Kern and Schrider, 2018]. We computed these summary statistics in 101 contiguous and non-overlapping windows (each of length 11 kb) across simulated sequences of length 1.1 Mb, resulting in input matrices of dimension 12 × 101, with summary statistics on the rows and computed values across windows along the columns. We then resized these matrices to dimension 299 × 299 to match the expected input image size of *InceptionResNetV2*. Moreover, though diploS/HIC was originally designed to differentiate among five classes, we retooled it to differentiate between two classes, as in other studies [Mughal et al., 2020, Arnab et al., 2023].

Next, we evaluated an alternate version of diploS/HIC by training a shallow CNN using the images produced by the native image generation method of *TrIdent*, which we refer to as *smbCNN* (shallow multi-branch CNN). We employed the identical CNN architecture used in diploS/HIC [Kern and Schrider, 2018] to extract features from input images. This shallow architecture is comprised of three branches, each containing two convolution layers with ReLU activation [Nair and Hinton, 2010] as well as max pooling [LeCun et al., 1998], dropout [Srivastava et al., 2014], and flatten layers, with the kernel sizes and dilation rates [Yu and Koltun, 2015] utilized in the convolution layers determining the differences among these three branches. These branches are then concatenated prior to adding two pairs of dropout and dense layers to form the final model.

By examining *TrIdent* [*IRV2*, *SS*] and *smbCNN*, we aim to understand the potential impact of incorporating summary statistics into the *TrIdent* [*IRV2*] architecture, as well as the effect of using shallow CNNs trained on *TrIdent* native images. Comparing *TrIdent* [*IRV2*], *TrIdent* [*IRV2*, *SS*], diploS/HIC, and *smbCNN* across the CEU (Figure 5) and YRI (Figure 6) datasets, we can assess the impact of different architectural choices on model performance. In both demographic test cases, *smbCNN* was the poorest performing model among the four, with accuracies of 80.1% and 83.7% on the CEU and YRI datasets, respectively (Figures 5 and 6). However, the two models employing summary statistics, *TrIdent* [*IRV2*, *SS*] (accuracy of 90.05% on CEU and 94.95% on YRI) and diploS/HIC (accuracy of 88.30% on CEU and 94.15% on YRI), consistently demonstrate higher classification accuracy compared to *TrIdent* [*IRV2*] (Figures 5 and 6). *TrIdent* [*IRV2*, *SS*] outperformed diploS/HIC in both cases. Prior to resizing, summary statistic-based images are significantly smaller than native *TrIdent* images (12×101 compared to 198×499 for CEU or 216×499 for YRI). Because different summary statistics computed within a given window are arranged along a column in consecutive rows, some correlation among values is anticipated. Moreover, values along rows, which are individual summary statistics computed across consecutive windows, are also expected to be correlated. For methods that utilize summary statistics as input, the arrangement of haplotypes within a window does not affect the computation, as the statistics are derived from the aggregated properties of the window rather than the specific ordering of haplotypes.

**Figure 5:**
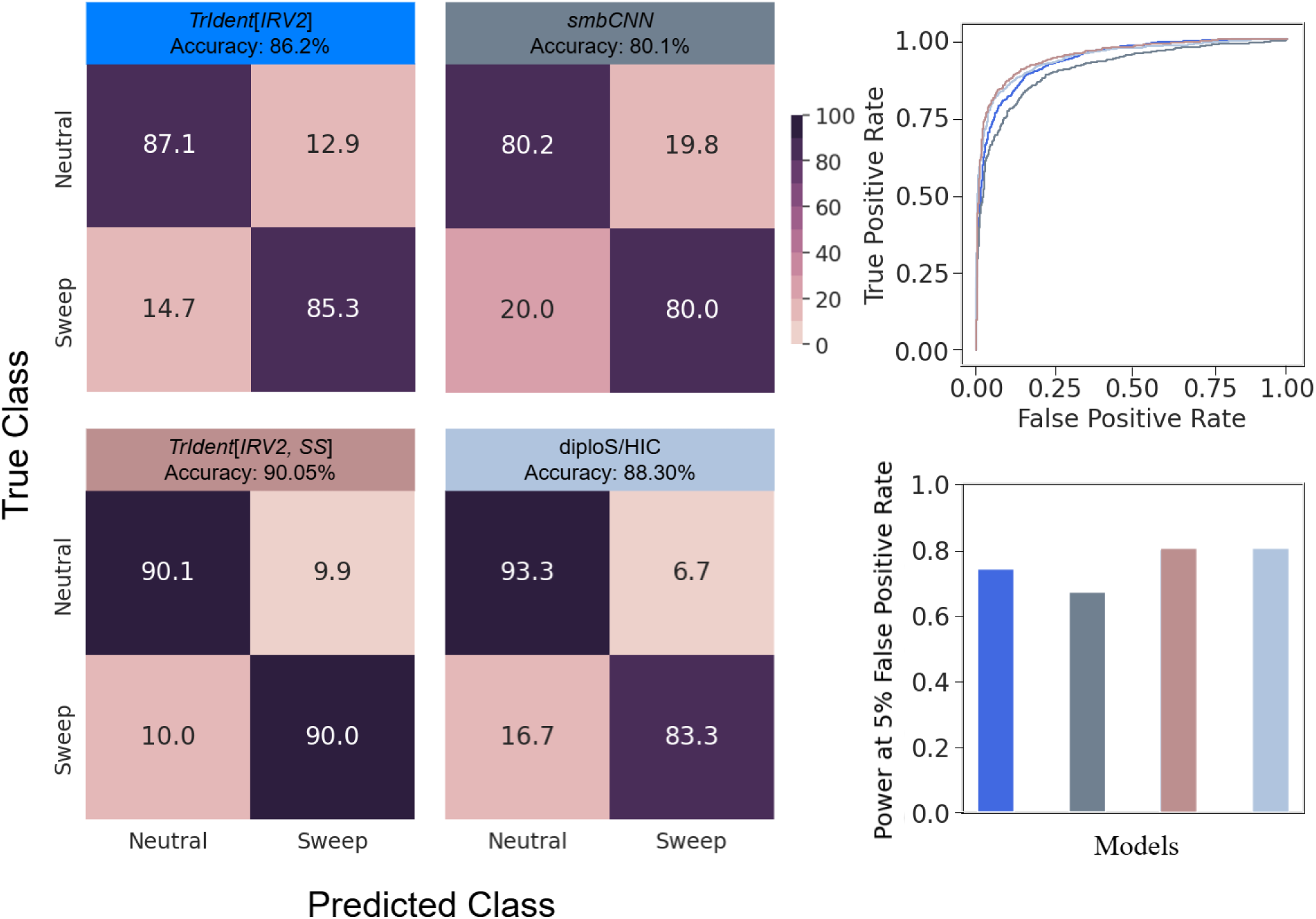
Classification rates and accuracies as depicted by confusion matrices, powers (true positive rates) to detect sweeps as depicted by receiver operating characteristic curves to differentiate sweeps from neutrality, and powers at a 5% false positive rate to detect sweeps on the CEU dataset for the best performing *TrIdent* model (*TrIdent* [*IRV2*]) compared to diploS/HIC and *smbCNN*. The *smbCNN* model represents a custom-built shallow CNN trained using *TrIdent* ’s native images, and *TrIdent* [*IRV2, SS*] represents an alternate *TrIdent* [*IRV2*] architecture trained using summary statistics based images (see *Viability of alternate architectures and methods* for details).

**Figure 6:**
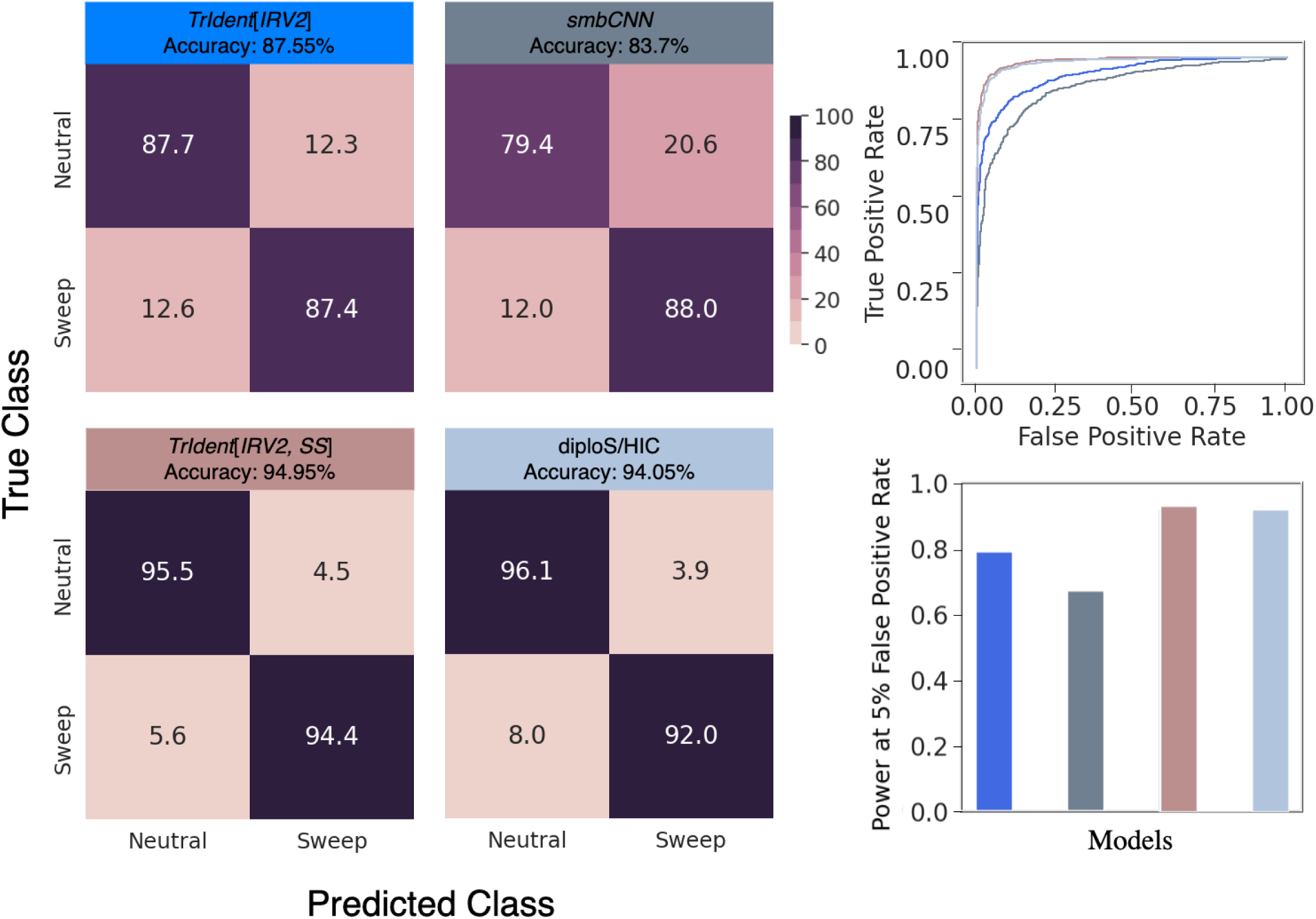
Classification rates and accuracies as depicted by confusion matrices, powers (true positive rates) to detect sweeps as depicted by receiver operating characteristic curves to differentiate sweeps from neutrality, and powers at a 5% false positive rate to detect sweeps on the YRI dataset for the best performing *TrIdent* model (*TrIdent* [*IRV2*]) compared to diploS/HIC and *smbCNN*. The *smbCNN* model represents a custom-built shallow CNN trained using *TrIdent* ’s native images, and *TrIdent* [*IRV2, SS*] represents an alternate *TrIdent* [*IRV2*] architecture trained using summary statistics based images (see *Viability of alternate architectures and methods* for details).

### Fine-tuning vs. training

Building upon our comparison of five pre-trained architectures in the previous subsection, we conducted an additional experiment to explore whether the strong performance of *TrIdent* is attributable to the model itself or to the pre-trained feature extraction capabilities of the *InceptionResNetV2* backbone. We developed a modified model, termed *IRV2* (*InceptionResNetV2* architecture trained on *TrIdent* images), which extends the *InceptionResNetV2* architecture by incorporating a GAP layer and an output layer consisting of a single node with a sigmoid activation function but does not include an elastic-net style penalty for parameter fitting. The model was trained directly on the *TrIdent* input images, with starting weights obtained from pre-trained weights of ImageNet.

The training dataset size for *IRV2* was the same as that utilized for the *TrIdent* [*IRV2*] model in one experiment (1,000 samples per class) and was increased to 10,000 samples per class in a second experiment. Following the training of *IRV2* on 1,000 samples per class, a slight enhancement in performance was noted in comparison to *TrIdent* [*IRV2*]. The accuracy improved by 1.7% on the CEU dataset (87.9% compared to 86.2%) and by 3.3% on the YRI dataset (90.85% compared to 87.55%) (Figure S5). When trained on 10,000 samples per class, *IRV2* achieved even higher accuracies: 90.85% on CEU and 92.45% on YRI. These results illustrate the capacity of *InceptionResNetV2* to achieve superior performance with increased number of training samples, consistent with observations in other domains where large-scale training datasets unlock additional classification accuracy and power [Yu et al., 2015, Alzubaidi et al., 2021].

The resource demands of training *IRV2* (see the *Computational resources and requirements* sub-section of the *Methods*) underscore the computational challenges inherent in training deep neural networks from scratch using simulated data, rather than utilizing pre-trained architectures. These findings highlight the balance between accuracy and computational efficiency. While *IRV2* out-performed *TrIdent* [*IRV2*], such gains come at the expense of significantly greater computational resources and training time. *TrIdent* [*IRV2*] demonstrates that by employing pre-trained feature extraction, strong performance can be achieved with substantially fewer training samples and lower computational overhead. This balance positions *TrIdent* [*IRV2*] as a practical and effective solution for natural selection detection, especially where computational resources are constrained.

### Performance in the presence of missing genomic regions

Due to undiscovered polymorphisms, haplotypic diversity can be reduced when segments of the genome are missing as a result of poor mappability, alignability, sequencing, or variant calling [Alkan et al., 2011, Talkowski et al., 2011]. As a consequence, this decline in local genomic variation can resemble a footprint of a selective sweep, misleading sweep detection approaches to mistakenly attribute such diversity as positive selection in neutrally-evolving genomic regions [Vitti et al., 2013, Mughal et al., 2020]. Hence, we aim to evaluate whether *TrIdent* [*IRV2*] and *T-REx*, both with native and alternate ways of generating images, mistakenly identify neutrally-evolving regions harboring missing genomic segments as selective sweeps as well as lose capacity to differentiate sweeps from neutrality under such settings. We removed polymorphisms from test replicates according to the human empirical distribution of CRG (Centre for Genomic Regulation) mappability and alignability scores (see *Filtration of empirical data* subsection of the *Methods*), which past studies have shown to negatively affect machine learning sweep classifiers [Mughal and DeGiorgio, 2019, Mughal et al., 2020, Arnab et al., 2023, Amin et al., 2023].

We simulated additional sets of neutral and sweep test replicates under both CEU and YRI demographic histories using discoal, and introduced missing regions using the identical protocol as Arnab et al. [2023] based on the empirical human distribution. Specifically, we randomly chose one of the 22 human autosomes with probability proportion to length of each autosome. From this autosome, we selected a starting genomic location uniformly at random for a 1.1 Mb segment. We removed SNPs that intersected with human genomic regions of low mappability and alignability (see *Filtration of empirical data and empirical image generation* subsection of the *Methods*). Because *TrIdent* only considers the central 499 SNPs when generating images from simulated replicates, we ensured that missing data blocks removed at least one of the central 499 SNPs, repeating the procedure until this criterion was satisfied. We found that this procedure removed on average approximately 23.1% of the original set of central 499 SNPs across all test replicates.

On the CEU test case, when *TrIdent* [*IRV2*] trained with its stock image generation technique is applied to replicates with missing polymorphisms, the overall accuracy drops by 4.45% (Figure S6) compared to no missing segments (Figure 3), whereas missing data causes *TrIdent* [*IRV2*, *alt*] trained on the alternate image generation technique to experience a drop in overall accuracy by 6.1% (Figure S6) compared to no missing data (Figure 3). In particular, while neutral detection rate was increased by roughly 1% with either style of input image, sweep detection rate was drastically decreased with missing segments regardless of image generation style (Figures 3 and S6). While *TrIdent* [*IRV2*] saw a sweep detection rate drop of 10.5%, *TrIdent* [*IRV2*, *alt*] trained on the alternate image generation style saw an even higher drop of 13%. Conversely, when *T-REx* trained with its stock image generation technique is applied to test replicates with missing segments, its overall accuracy dropped by 3.65% (Figure S6) compared to no missing polymorphisms (Figure 3). On the other hand, *T-REx* [*alt*] trained on the alternate image generation style undergoes an 11.3% decrease accuracy (Figure S6) compared to no missing segments (Figure 3). The neutral detection rate is preserved for both image generation styles, though the sweep detection rate for*T-REx* [*alt*] showcases a significantly greater drop of 22.7% compared to the 7.1% decrease in classification accuracy for *T-REx* when both are applied to missing polymorphisms (Figures 3 and S6).

On the YRI test case, however, the drop in test accuracies of all four models when presented with replicates containing missing genomic regions (Figure S7) appears to be less severe than in the CEU test case (Figure S6). Our results show a loss of 4.05% by *TrIdent* [*IRV2*], 2.65% by *TrIdent* [*IRV2*, *alt*], 2.80% by *T-REx*, and 7.50% by *T-REx* [*alt*] (Figure S7) compared to test set with no missing data (Figure 4). All four models present comparable neutral classification rates (from 84.3% to 88.5%). On the other hand, with a sweep detection rate of 81.7%, *TrIdent* [*IRV2*] again establishes itself as the most robust against missing genomic regions. The overall test accuracies lend further support to our observation that the *TrIdent* models, with accuracies of 83.50% and 82.65%, comfortably outperform the *T-REx* models, with accuracies of 80.0% and 76.35% (Figure S7). Our findings demonstrate that the drops in accuracies for the *TrIdent* and *T-REx* model variants under missing genomic segments is generally driven by misclassification of sweep replicates as neutral, rather than the false attribution of selection for neutral settings.

### Robustness against background selection

Background selection poses a potential challenge for *TrIdent* [*IRV2*], as it has the potential to mislead sweep classifiers into falsely detecting signatures of positive selection. The occurrence of this phenomenon is caused by the elimination of harmful genetic variations through negative selection, resulting in distortions in the distribution of allele frequencies that may resemble positive selection [Charlesworth et al., 1993, Hudson and Kaplan, 1995, Charlesworth, 2012]. Specifically, background selection can produce patterns in allele frequency distributions that resemble those caused by selective sweeps [Charlesworth et al., 1993, 1995, 1997, Keinan and Reich, 2010, Seger et al., 2010, Nicolaisen and Desai, 2013], which can result in misclassification of evolutionary events [Huber et al., 2016]. However, recent studies suggest that selective sweeps and background selection leave distinct genetic footprints, particularly in haplotype distributions indicating that background selection is unlikely to pose a significant issue when analyzing haplotype data [Fagny et al., 2014, Schrider, 2020, Lauterbur et al., 2023]. Nevertheless, it is imperative to evaluate the resilience of *TrIdent* [*IRV2*] to the pervasive force of background selection that shapes patterns of genomic variation [McVicker et al., 2009, Comeron, 2014].

We generated 1,000 test replicates using the forward-time simulator SLiM [Haller and Messer, 2019] under the same genetic and demographic parameters of the CEU and YRI simulations used to train *TrIdent*, with the addition of background selection [Tennessen et al., 2012, Adrion et al., 2020]. Specifically, in a simulated 1.1 Mb region, a 55 kb protein-coding gene was subjected to negative selection according to the methodology described by Cheng et al. [2017]. This gene was composed of 50 exons each of length 100 bases, 49 introns each of length one kb, a 5*^′^* untranslated region (UTR) of length 200 bases, and a 3*^′^* UTR of length 800 bases, which are fairly close to mean lengths for these elements in human genomes [Mignone et al., 2002, Sakharkar et al., 2004]. Recessive (*h* = 0.1) deleterious mutations with selection coefficients (*s*) drawn from a gamma distribution with mean of -0.0294 and shape parameter of 0.184 [Boyko et al., 2008, Schrider and Kern, 2017] arose within this gene, where 75%, 10%, and 50% of mutations in exons, introns, and UTRs, respectively, were deleterious. Consequently, we used the procedure outlined in the *Image Generation* subsection of the *Methods* to create input images, which we subsequently fed into trained *TrIdent* [*IRV2*] models as test observations.

Our findings suggest that the *TrIdent* [*IRV2*] models trained on the CEU and YRI datasets are resilient to background selection (Figure S8), with the incidence of false sweep signals attributed to background selection closely paralleling the false positive rate obtained from neutral test observations (Figure S8), particularly at acceptable false positive rates (Figure S8; bottom panels). This robustness to background selection is not only observed for settings in which recombination rates are drawn from the same distribution as the training sets (Figure S8, left panels), but also for the challenging setting in which mean recombination rates are an order of magnitude smaller than what was used in the training sets (Figure S8, right panels). Furthermore, for a mean recombination rate of 10*^−^*^8^, *TrIdent* [*IRV2*] models identified 92.15% and 94% of background selection simulations as neutral for the CEU and YRI datasets, respectively. These rates are higher than the neutral detection rates of 87.1% and 87.7% for respective CEU and YRI neutral simulations (Figures 3 and 4), indicating that *TrIdent* [*IRV2*] is even less likely to falsely attribute background selection as a sweep than it is for neutrally-evolving regions.

### Performance with unphased multilocus genotypes

Sweep classifiers can face challenges when applied to unphased multilocus genotype data, which harbor less information than phased haplotypes, as selection acts to alter frequencies of nearby neutral haplotypic variation and only indirectly on multilocus genotype variation. Using unphased genotypes also decreases statistical power and introduces complexity in interpreting genetic signals, potentially omitting subtle yet significant signatures of selection within populations. However, *TrIdent* models are trained with phased haplotypic data, which is often difficult or impossible to reliably generate for many study systems, especially for most non-model organisms. Thus, to enhance the versatility of these models, it is crucial that they can accommodate unphased data. Given the demonstrated capability to detect sweeps using unphased multilocus genotypes in prior studies [Kern and Schrider, 2018, Harris and DeGiorgio, 2020, Mughal and DeGiorgio, 2019, Gower et al., 2021, Arnab et al., 2023], we expect that *TrIdent* will continue to achieve excellent classification accuracy and power when applied to unphased data.

To generate *TrIdent* input images from unphased data, we first merge pairs of rows (haplotypes) into a single multilocus genotype to create values of zero, one, and two representing the number of copies of the minor allele at a diploid genotype. We then followed the same procedure described in the *Image generation* subsection of the *Methods* to create images for the *TrIdent* [*IRV2*] model. We refer to the *TrIdent* [*IRV2*] model using images from unphased multilocus genotype data as *TrIdent* [*IRV2*, *MLG*]. We find that for both the CEU and YRI (Figure S9) test cases, *TrIdent* [*IRV2*, *MLG*] achieves comparable, though marginally lower, accuracies and powers to *TrIdent* [*IRV2*], with a 1.95% drop in classification accuracy on the CEU test case and a more narrow drop of 0.15% in accuracy on the YRI test case. Moreover, under both scenarios, *TrIdent* [*IRV2*, *MLG*] is more conservative than *TrIdent* [*IRV2*], with a slight bias toward predicting observations as neutral (Figure S9). Overall, these results indicate that while *TrIdent* [*IRV2*, *MLG*] is slightly less accurate than *TrIdent* [*IRV2*], it remains a robust method for analyzing unphased multilocus genotype data, demonstrating the flexibility of the *TrIdent* approach.

### Capacity to uncover incomplete sweeps

The less pronounced genomic footprints of incomplete or partial sweeps may increase false negative rates of sweep classifiers by misleadingly assigning such patterns as neutral [Schrider et al., 2015, Xue et al., 2021]. Specifically, these genomic signals are less pronounced than what is expected by recent complete sweeps, including localized and weaker linkage disequilibrium, shorter and less frequent haplotypes, a more mildly distorted site frequency spectrum, and a comparatively more marginal decrease in genetic variation [Vy and Kim, 2015]. As a result, it is important to evaluate the power of *TrIdent* to detect incomplete sweeps relative to other sweep classifiers. To evaluate power under this setting, we used discoal [Kern and Schrider, 2016] to generate sweep test replicates for both the CEU and YRI demographic histories for which an advantageous allele does not reach fixation. Specifically, we simulated 1,000 replicates for each incomplete sweep scenario, considered situations for which the beneficial allele stopped being advantageous at a frequency of *f*_end_ ∈ {0.5, 0.6, 0.7, 0.8, 0.9} while fixing all other genetic, demographic, and selection parameters as detailed within the *Simulation protocol* subsection of the *Methods*. We then applied *TrIdent* [*IRV2*] and *T-REx* originally trained on complete sweeps to these incomplete sweep test sets.

We evaluated both accuracy and power (true positive rate) at a 5% false positive rate to detect incomplete sweeps (Figure S10). Under the CEU demographic history, accuracy for both *TrIdent* [*IRV2*] and *T-REx* climbed with increasing final frequency of the beneficial allele *f*_end_, with values as low as 31.06% and 21.7% at *f*_end_ = 0.5 for *TrIdent* [*IRV2*] and *T-REx*, respectively, and achieving values of 83.15% and 70.1% at *f*_end_ = 0.9 for *TrIdent* [*IRV2*] and *T-REx*, respectively. Notably, *TrIdent* [*IRV2*] achieves an accuracy close to 80% for incomplete sweeps to a frequency of *f*_end_ = 0.8, whereas *T-REx* never reaches 80% sweep detection rate even for complete sweeps for which *f*_end_ = 1.0 (Figure S10). A similar increasing trend in power as a function of the degree of sweep completeness can be observed (Figure S10), with values as low as 0.201 and 0.11 at *f*_end_ = 0.5 for *TrIdent* [*IRV2*] and *T-REx*, respectively, and achieving values of 0.73 and 0.56 at *f*_end_ = 0.9 for *TrIdent* [*IRV2*] and *T-REx*, respectively. Moreover, *TrIdent* [*IRV2*] achieves a power slightly above 0.7 at *f*_end_ = 0.8 (which is virtually identical to the power for a complete sweep), whereas *T-REx* never even achieves a power of 0.6 for complete sweeps. Thus, *TrIdent* [*IRV2*] consistently and significantly outclassed *T-REx* in terms of accuracy and power to detect incomplete sweeps.

For the YRI test case, we observed an overall higher accuracy and power for detecting incomplete sweeps compared to the CEU test case. Specifically, the accuracy for both *TrIdent* [*IRV2*] and *T-REx* increased with *f*_end_. At *f*_end_ = 0.5, the accuracy values for *TrIdent* [*IRV2*] and *T-REx* were 44.11% and 38.6%, respectively, and climbed to 87.03% for *TrIdent* [*IRV2*] and 78.99% for *T-REx* at *f*_end_ = 0.9. A similar trend was seen in power, with values starting from 0.307 and 0.212 at *f*_end_ = 0.5 for *TrIdent* [*IRV2*] and *T-REx*, respectively, and respectively rising to 0.7404 and 0.607 at *f*_end_ = 0.9. These results show that *TrIdent* [*IRV2*] consistently and significantly outperformed *T-REx* in both accuracy and power for the YRI test case, expectedly surpassing the performance levels observed in the CEU test case that includes a recent, severe population bottleneck. In general, the results highlight the ability of *TrIdent* in recognizing even modest adaptive signals in genomes.

### Interpretability of the sweep classifier

In non-natural image classification tasks, class activation maps can reveal the decision-making process of a trained model [Zhai and Shah, 2006, Zhou et al., 2016]. In particular, these maps are able to highlight regions of pixels in non-natural images, like those generated here to discriminate sweeps from neutrality, where patterns may not be immediately discernible to humans that explain or validate what features the model places emphasis when classifying observations. In addition, class activation maps can aid in model selection based on performance and robustness in image classification tasks by acting as a diagnostic tool to identify errors in the learning process of a model by visualizing key regions of input images, allowing for targeted adjustments to improve accuracy and generalization and thereby creating more transparent and reliable classification systems [Zhou et al., 2016].

For pre-trained CNN models, gradient-weighted class activation mapping (GradCAM) provides a mechanism for creating class activation maps that are easy to understand and work with [Selvaraju et al., 2017]. To accomplish this task, GradCAM makes use of the gradient information that flows into the last convolutional layer of the CNN during its backward pass. It computes the importance of each feature map by taking the gradients of the target class score with respect to these feature maps and weighting them accordingly. By highlighting the relative importance of each pixel, this approach successfully pinpoints the regions within an image that help discriminate among classes [Selvaraju et al., 2017].

We employed GradCAM to generate class activation maps from each of the training set images based on their output values from the last convolution layer of *TrIdent* [*IRV2*]. The resulting maps for both the CEU and YRI test cases revealed a concentrated focus in the lower-middle region of the images (Figure 7), coinciding with regions of input images that might distinguish sweeps from neutrality on average (Figure 2). Moreover, this consistent regional focus across both CEU and YRI test cases suggests that *TrIdent* [*IRV2*] effectively captures the critical haplotype features driving the divergence between sweep and neutrality in these populations.

**Figure 7:**
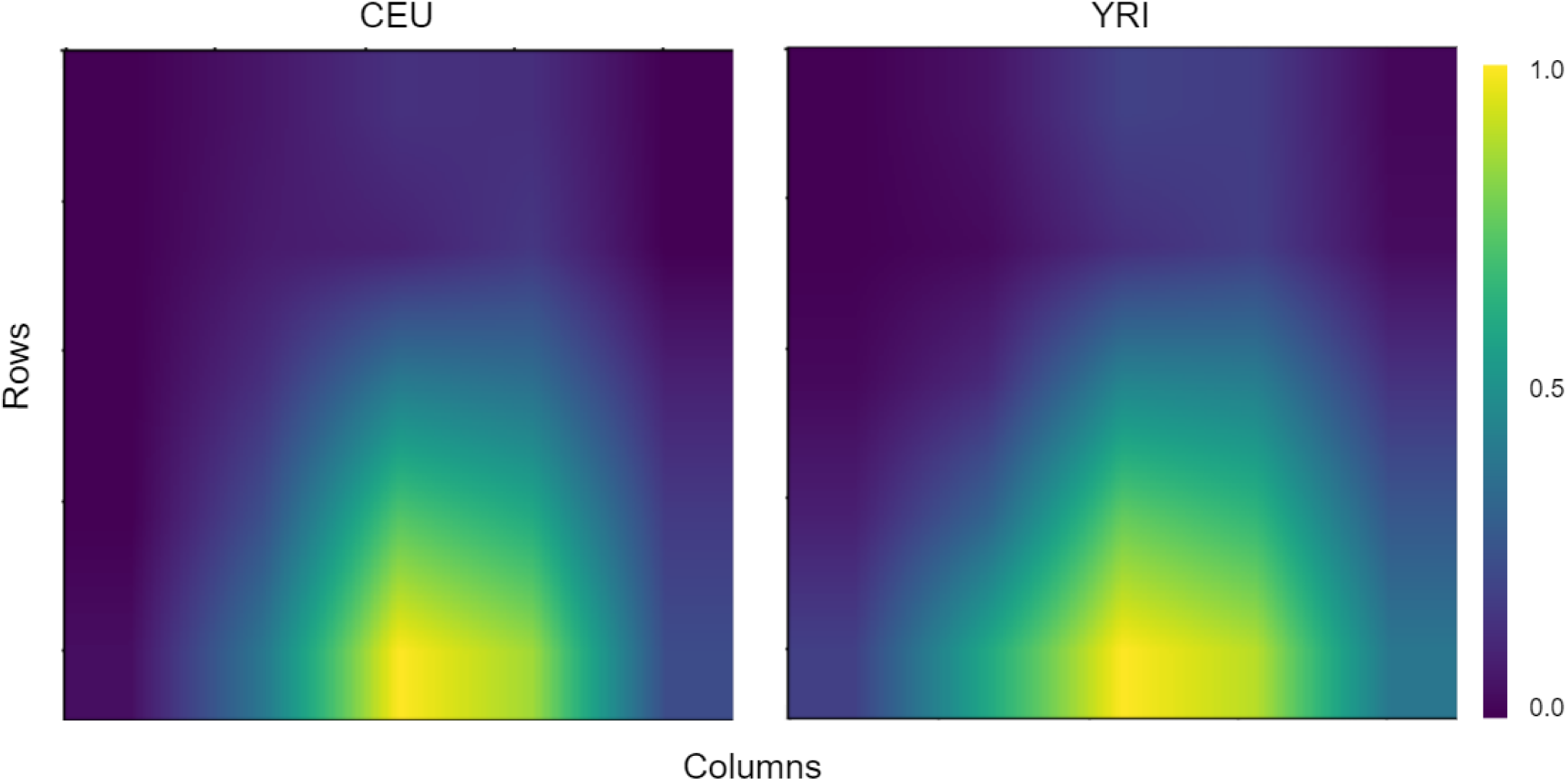
Heatmaps of mean gradient-weighted class activation maps (GradCAM) from *InceptionResNetV2* applied to CEU (left panel) and YRI (right panel) training data, with the mean taken across 2,000 training observations with 1,000 observations from each of the neutral and sweep classes.

### Ability to predict sweep parameters

So far, we have assessed the sweep detection accuracy and power of *TrIdent* [*IRV2*] in relation to the sweep classifiers *T-REx*, diploS/HIC and many architectural variations of the the *TrIdent* model. We next consider how features obtained from transfer learning architectures fare on regression tasks for predicting evolutionary parameters of sweeps. We trained three distinct models, each tasked with inferring one of the three selection parameters used to generate sweep replicates for training the *TrIdent* [*IRV2*] classifier. These parameters are: the number of generations in the past when the beneficial allele reached fixation (*τ*), the selection strength acting on the beneficial allele (*s*), and the frequency of the beneficial allele at the onset of selection (*f*). As with the *TrIdent* classifiers, we first extracted GAP layer outputs of *InceptionResNetV2* from the 1,000 sweep replicates to represent the set of features in the training dataset. As the output for each regression model, we performed a logarithmic transformation of the parameters so that they take both positive and negative values, rather than only non-negative values, as well as to better highlight parameters that were drawn across different orders of magnitude (see *Modeling description* subsection).

Our initial evaluation of a linear model to predict the selection parameters showed that such a model was not accurate enough based solely on the *InceptionResNetV2* extracted features. We therefore chose to employ nonlinear models based on ANNs for predicting sweep parameters, and we term this nonlinear regression model *TrIdent* [*IRV2*, *ANN*], similar to the nonlinear classification model of the *Viability of alternate architectures and methods* subsection. To determine the architecture of each sweep parameter predictor model (*f*, *s*, and *τ*), we use a model selection scheme identical to that of the nonlinear *TrIdent* [*IRV2*, *ANN*] classifier, with the exception of assessing model fit with mean squared error (MSE) loss instead of binary cross entropy along with the usage of linear activation in the output layer compared to sigmoid activation. Here, we choose the best number of hidden layers, number of nodes in each layer, and activation function used in each layer for each of the three regression models based on smallest MSE validation error.

To compare distributions between true and predicted parameter values, we summarize the distributions using violin plots to capture distribution shapes with embedded box plots to depict distribution locations and spreads (Figure 8). For *τ*, the medians for true and predicted distributions are comparable in both CEU and YRI populations. The MSE is 0.0198 for CEU and 0.0243 for YRI, and the distributions are left skewed, with the difference between the second and first quartiles of the predicted distribution smaller in the YRI than in CEU. For *s*, the MSE is 0.0383 for CEU and 0.0114 for YRI, with the predicted distributions preserving the characteristic “lip” shape observed in the true distribution, though with predicted distributions tighter than the true distributions. The medians of the true and predicted distributions are comparable in both datasets, but the inter-quartile range (IQR) is narrower in the predicted distributions. For *f*, the predicted distributions display skewed “lip” shapes—right-skewed for CEU and left-skewed for YRI—though the true distributions show a more symmetrical “lip” shape. Moreover, the predicted median is shifted slightly downward relative to the true median in the CEU, whereas it is shifted upward in the YRI. However, whereas the IQR is smaller for the predicted distribution in the CEU dataset than for the true distribution, the IQR for the predicted distribution in the YRI dataset is comparable to that of the true distribution, with MSE values of 0.2001 and 0.1665 for CEU and YRI, respectively. While the MSE values may appear high, it is important to note that the parameters were logarithmically scaled, which stretches their bounds, inherently inflating the apparent magnitude of the MSE compared to unscaled values. Overall, in terms of MSE, *TrIdent* [*IRV2*, *ANN*] seems to perform better on predicting *s* and *f* on the YRI dataset compared to on the CEU dataset, and *τ* on the CEU dataset compared to on the YRI dataset.

**Figure 8:**
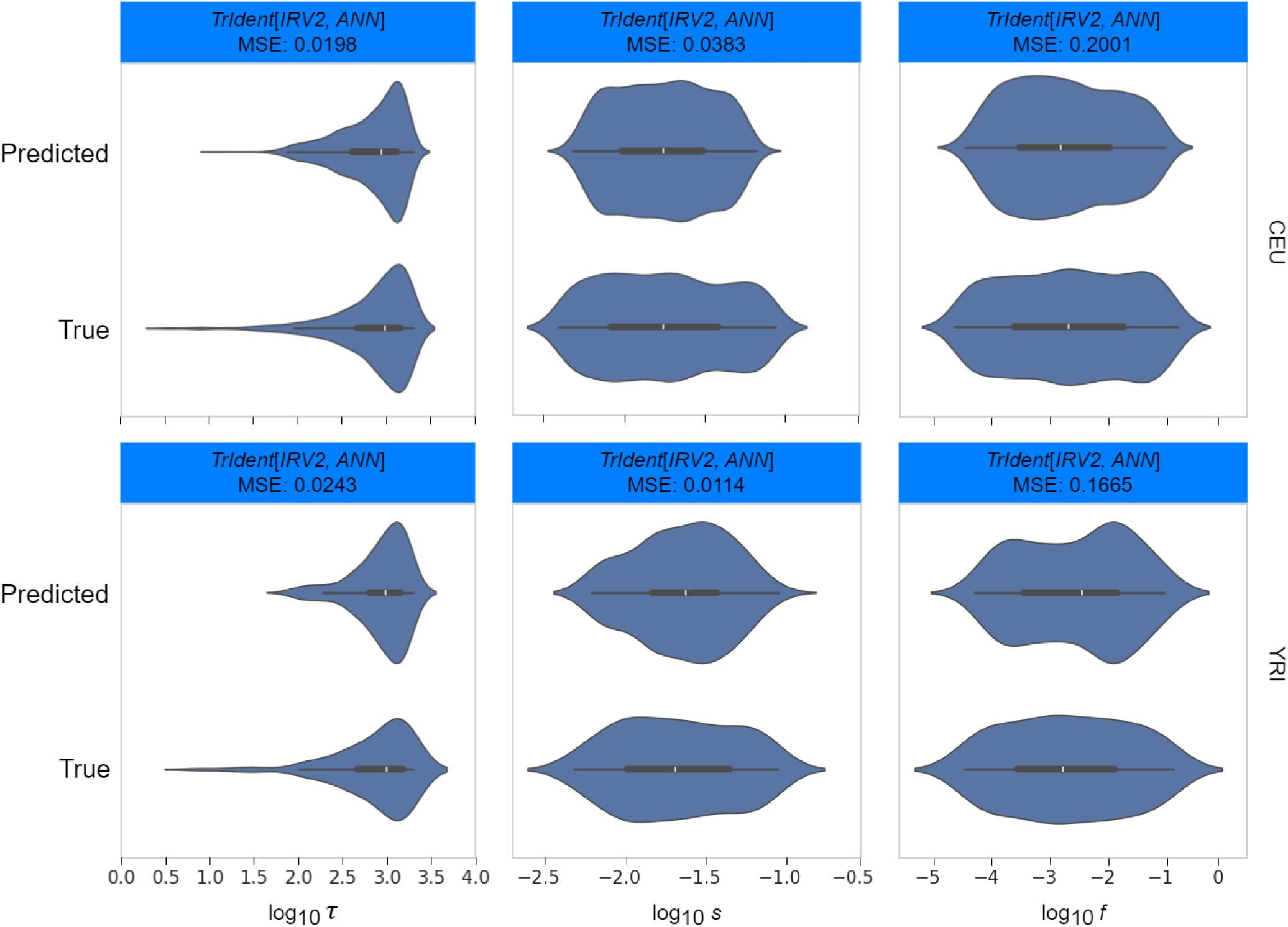
Summaries of distributions for true and predicted values of selection parameters (*f*, *s*, and *τ*) using the nonlinear *TrIdent* [*IRV2*, *ANN*] regression model. Distributions are summarized using violin plots with embedded box plots for the CEU (top) and YRI (bottom) datasets.

To benchmark against a comparative baseline for selection parameter estimates, we employed the selection coefficient inference tool CLUES2 [Vaughn and Nielsen, 2024]. While CLUES2 is limited to estimating the selection coefficient, one of the three parameter estimates by the *TrIdent* [*IRV2*, *ANN*] regression model, it serves as a useful reference point. Details of our CLUES2 pipeline are presented in the *Application of CLUES2* subsection of the *Methods*. We generated ancestral recombination graphs (ARGs) using the entire 1.1 Mb simulated replicates using SINGER [Deng et al., 2024], as it is one of the two software supported by CLUES2. We implemented CLUES2 as closely to the recommended settings as possible, given the constraints imposed by our simulated replicates. In both CEU and YRI test cases, CLUES2 lagged considerably behind the accuracy and precision of *TrIdent* [*IRV2*, *ANN*] (Figure S11). To maintain consistency with *TrIdent* [*IRV2*, *ANN*], we applied a logarithmic transformation to the selection coefficient estimates from CLUES2, which resulted in MSEs on this transformed scale of 0.454 for CEU and 0.402 for YRI.

### Application to human genomic data

Given the excellent performance of *TrIdent* [*IRV2*], we next applied it to variant calls and phased haplotypes of 99 individuals in the CEU population and 108 individuals in the YRI population from the 1000 Genomes Project [The 1000 Genomes Project Consortium, 2015]. This application served as a proof-of-concept, where we evaluate the ability of *TrIdent* to recapitulate sweep candidates established in the literature from European and African humans, as well as to potentially discover novel candidates.

Before scanning the CEU and YRI genomes, we eliminated SNPs in specific regions using the procedure detailed in the *Filtration of Empirical Data* subsection of the *Methods*. Following the elimination of these SNPs, we selected the initial 499 SNPs of a chromosome and created a 299×299 input image of haplotype variation (see *Modeling description* subsection of the *Results*). From this starting image, we created new empirical input images by advancing the 499-SNP window by a stride of 10 SNPs along the chromosome, repeating the process for all 22 autosomes. These images are then fed through the *TrIdent* [*IRV2*] pre-trained model, and the GAP layer outputs are used as input to the trained logistic regression classifier (see *Modeling description* subsection of the *Results*).

In the CEU scan, we found that the majority of genomic windows are classified as neutral with a probability threshold of 0.9 for calling sweeps (approximately 95.18%; Table 1). The threshold of 0.9 for individual windows is strategically chosen to reduce the risk of false positives, thus prioritizing the most significant candidate sweeps and improving the reliability of the detection process. We evaluated the false positive and true positive rates at this threshold of 0.9 using the CEU and YRI simulated test sets (Figure S12). This stringent threshold demonstrated its effectiveness in minimizing false positives, achieving high detection rates for neutral replicates (97.5% for CEU and 97.7% for YRI, compared to 87.1% for CEU and 87.7% for YRI with a threshold of 0.5). While the detection rates for sweep replicates were notably lower (69.9% for CEU and 72.2% for YRI, compared to 85.3% for CEU and 87.4% for YRI with a threshold of 0.5), this reflects a more conservative approach to identifying true adaptive signals.

**Table 1:** Percentage of windows classified as a sweep by *TrIdent* under sweep probability threshold of 0.9 for each of the autosomes within the CEU and YRI 1000 Genomes Project populations.

| Chromosome | CEU | YRI |
| --- | --- | --- |
| 1 | 5.48 | 4.84 |
| 2 | 4.68 | 3.78 |
| 3 | 4.02 | 2.16 |
| 4 | 4.65 | 6.11 |
| 5 | 4.22 | 4.64 |
| 6 | 4.47 | 5.06 |
| 7 | 4.38 | 4.40 |
| 8 | 5.92 | 1.92 |
| 9 | 5.15 | 4.51 |
| 10 | 3.96 | 3.77 |
| 11 | 3.75 | 4.40 |
| 12 | 4.76 | 3.86 |
| 13 | 3.89 | 3.66 |
| 14 | 5.32 | 3.78 |
| 15 | 5.10 | 3.52 |
| 16 | 5.53 | 5.14 |
| 17 | 6.67 | 3.32 |
| 18 | 4.34 | 2.01 |
| 19 | 7.08 | 6.69 |
| 20 | 6.17 | 3.06 |
| 21 | 4.97 | 4.07 |
| 22 | 7.17 | 2.65 |

Additionally, we require a minimum mean prediction probability of 0.9 across 10 consecutive prediction windows for sweep footprint detection. This stringent threshold is designed to filter out potentially spurious observations with high sweep probability, ensuring that only the most robust signals are identified [Arnab et al., 2023]. This approach resulted in 1,206 candidate sweep peaks, which intersected 575 genes. Out of these genes, *LCT* shows a clear peak with a 10-window mean peak probability score of 0.93 (Figure 9A). The detection of *LCT* serves as a positive control, because it has been identified as a recent sweep candidate in numerous studies [Tishkoff et al., 2007, Field et al., 2016, Ségurel and Bon, 2017] with strong estimates of selection pressure [Bersaglieri et al., 2004, Gerbault et al., 2009]. The *LCT* gene codes for the lactase enzyme that hydrolyzes lactose, a disaccharide in milk and dairy products. Early agriculture and dairy farming in Europe profoundly influenced lactase persistence. Domesticating animals for milk grew more common when hunter-gatherer societies became sedentary agricultural groups. We also applied the *TrIdent* regression model to predict selection parameters at *LCT* (Table 2), with estimates that a sweep on a standing genetic variant at frequency *f* ≈ 0.01 became beneficial with strength *s* ≈ 0.06 and completed *τ* ≈ 46 generations ago (approximately 1,300 years ago assuming a generation time of 29 years). For comparison, Bersaglieri et al. [2004] estimated that the persistence-associated haplotype began to increase rapidly in frequency between 2,188 and 20,650 years ago from selection with coefficients ranging from 0.09 to 0.19 across different European populations at this locus. Thus, the estimates produced by *TrIdent* seem to generally agree well with the findings of Bersaglieri et al. [2004].

**Figure 9:**
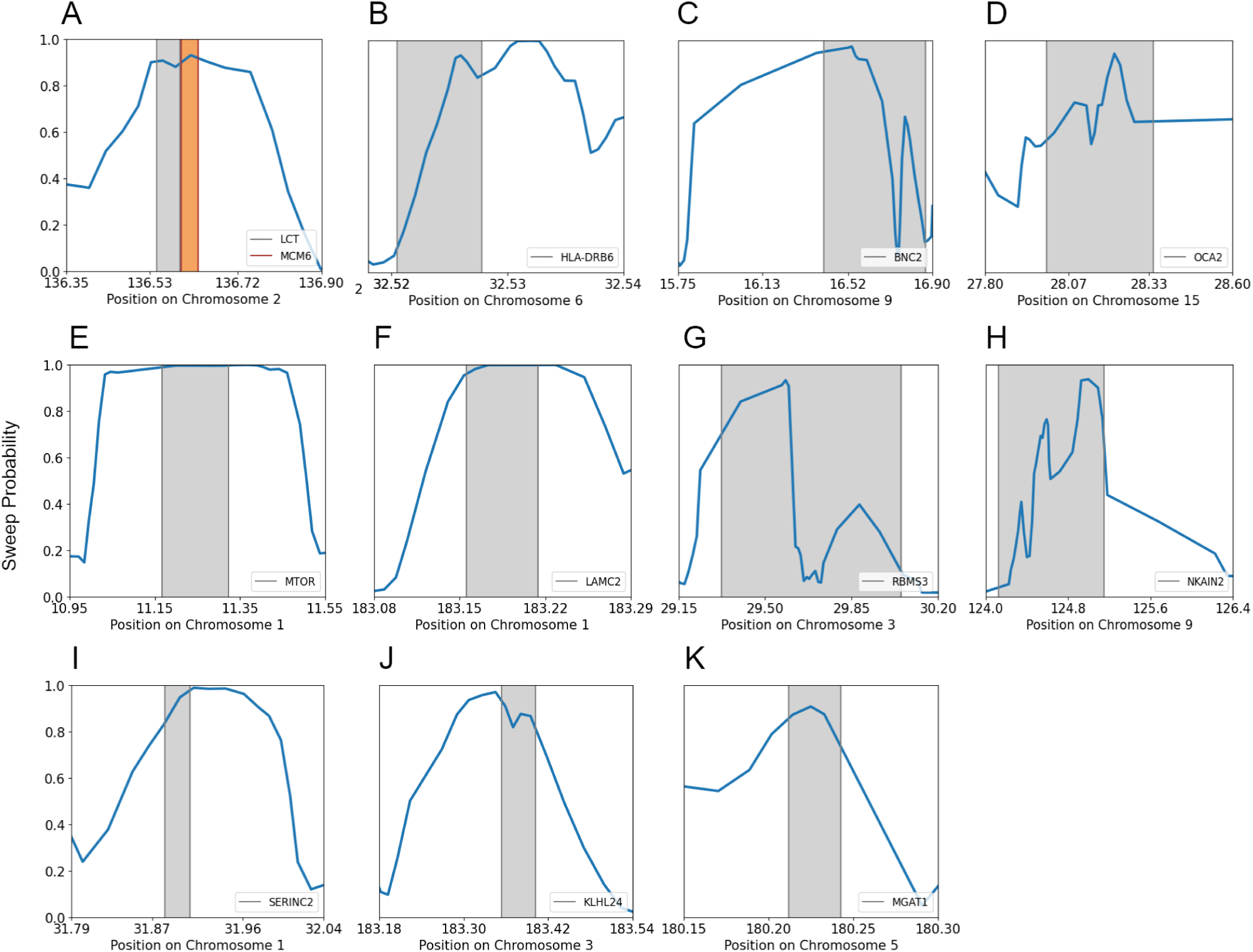
Identified candidate sweep regions from the genome-wide scan produced using the trained *TrIdent* [*IRV2*] model on the central European humans (CEU) population in the 1000 Genomes Project dataset. Regions were classified as being under positive selection if they had ten consecutive windows with a sweep probability higher than 0.9. A total of 575 genes across 22 autosomes exhibit qualifying signs of selective sweeps, of which a few of the most interesting candidates are reported here. Figure S21 provides a visual representation of the haplotype diversity surrounding the candidate genes in the plotted panels.

**Table 2:**
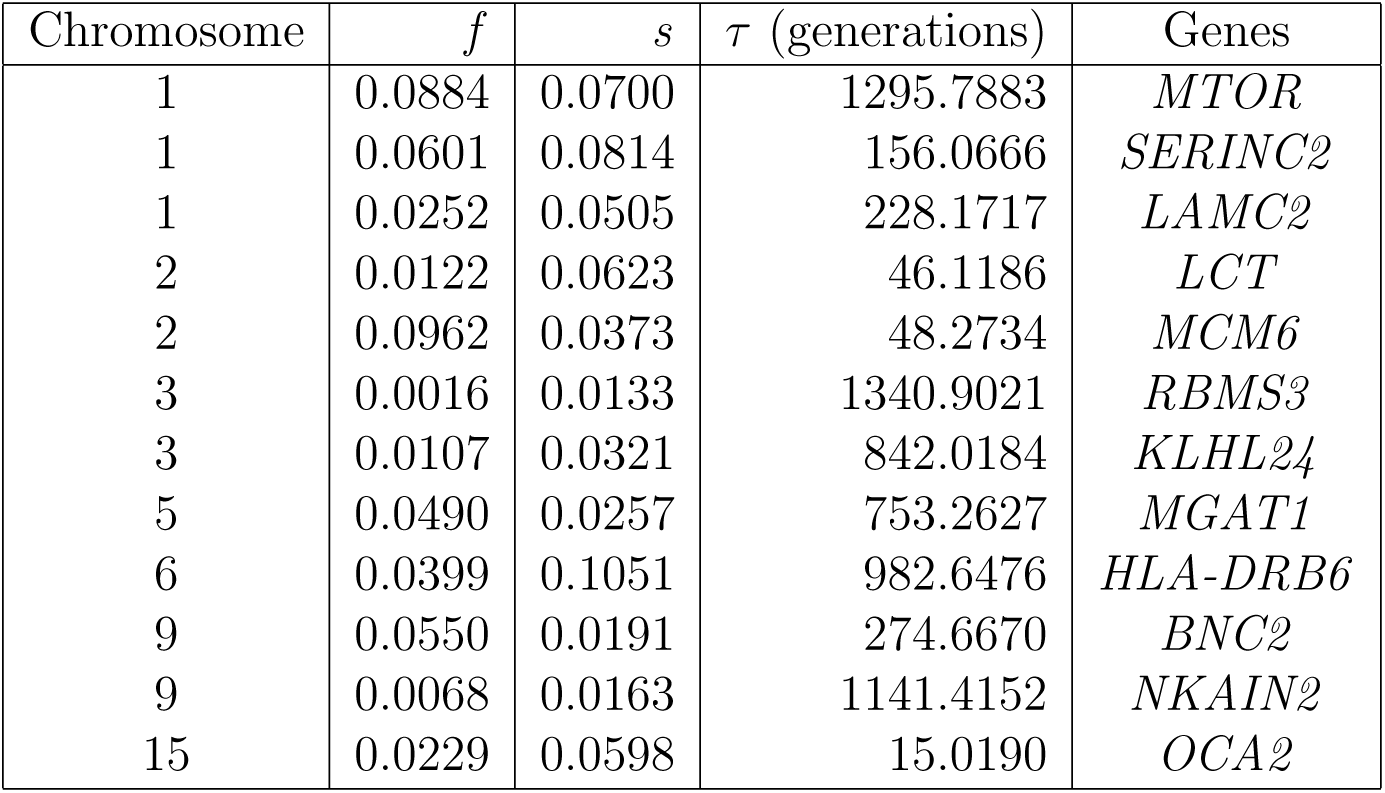
*TrIdent* regression model inferences for the frequency at which a selected allele became beneficial (*f*), selection strength (*s*), and time at which the beneficial allele reached fixation (*τ*) for genes reported in Figure 9 or the CEU scan.

Moreover, the *MCM6* gene located upstream of *LCT* not only exceeds the mean prediction probability threshold of 0.9 but also has a higher probability compared to *LCT* (Figure 9A). Selection parameter predictions at this gene (Table 2) support a sweep on a standing genetic variant at frequency *f* ≈ 0.1 became beneficial with strength *s* ≈ 0.04 and completed *τ* ≈ 48 generations or approximately 1,400 years ago. Previous studies also detected strong selection signals at *MCM6* within Europeans, though their selection coefficient estimates were generally lower (0.0161 in Stern et al. [2019] and 0.018 in Mathieson and Mathieson [2018]). The time frame that selection ended is similar to that of *LCT*, but with a softer sweep of lower strength. Due to its regulatory control of the expression of *LCT* [Labrie et al., 2016, Anguita-Ruiz et al., 2020], *MCM6* also serves a positive control. An enhancer within one of the introns of the *MCM6* gene has been discovered to impact the production of the lactase enzyme [Anguita-Ruiz et al., 2020], and has been found to show sweep signals by other studies [Oleksyk et al., 2010, Cheng et al., 2017, Amin et al., 2023].

One of the most pronounced sweep candidates identified by *TrIdent* is *OCA2*, with a 10-window mean sweep probability of 0.97 (Figure 9D), estimated to have completed *τ* ≈ 15 generations ago (Table 2), indicating that a putative sweep at *OCA2* completed more recently compared to that of *LCT* or *MCM6*. Previous studies have found *OCA2* as a target of positive selection in Europeans [Voight et al., 2006, Sulem et al., 2007, Wilde et al., 2014, Mughal et al., 2020], and therefore this gene also serves as an additional positive control. We estimate a selection coefficient of *s* = 0.059 for *OCA2*, indicating strong selection at this locus. For comparison, Stern et al. [2019] also identified evidence of selection at *OCA2*, estimating the strength at *s* = 0.04, whereas Mughal et al. [2020] reported *s* = 0.06 for the same locus. Hence, our selection strength estimates are consistent with prior estimates at this gene. Eye color is associated with variants within the *OCA2* gene, which plays a crucial role in the production and dispersion of melanin, the pigment that gives color to hair, skin, and eyes [Duffy et al., 2007]. The amount and type of iris melanin, which determines eye color, can vary due to structural variations in *OCA2* [Sturm and Larsson, 2009].

The human leukocyte antigen (HLA) system serves as a compelling example of how natural selection has operated to preserve genetic diversity. HLA genes, particularly those in class I and class II, play a crucial role in presenting antigens to T cells, which are responsible for initiating cell-mediated immune responses [Shankarkumar, 2004]. The extensive range of alleles and notable polymorphisms observed in HLA class I and class II genes [Hedrick and Thomson, 1983] serves as evidence of the ongoing selection that promotes genetic diversity at these immune-related locations. *TrIdent* follows the recent trend of sweep detectors [Goeury et al., 2018, Kern and Schrider, 2018, Harris and DeGiorgio, 2020, Amin et al., 2023, Arnab et al., 2023] in uncovering genes within the HLA region. Notably, we found that the class II HLA gene *HLA-DRB6* shows a clear signal consistent with positive selection, attaining a 10-window mean sweep probability of 0.92 (Figure 9B). Other studies have identified sweep signals at this gene in Europeans [DeGiorgio and Szpiech, 2022, Arnab et al., 2023], as well as other class I and class II genes within the region [Albrechtsen et al., 2010, Goeury et al., 2018, Harris and DeGiorgio, 2020, DeGiorgio and Szpiech, 2022]), lending credence to this finding. Furthermore, *TrIdent* predicts a notably high selection coefficient of *s* = 0.1051 for *HLA-DRB6*. Mughal et al. [2020] have also reported high selection coefficients for an HLA gene, finding *s* = 0.14 at *HLA-DRB1*, further supporting the notion that genes in the HLA region have been subject to intense selective pressures.

*BNC2* is another well-supported sweep candidate with a peak mean sweep probability of 0.99 (Figure 9C). This gene encodes a protein that plays a vital role in various essential cellular processes. These processes include controlling the expression of genes that code for proteins responsible for binding and regulating collagen, as well as for facilitating the growth of new tissues [Orang et al., 2023]. Its involvement in key developmental pathways is supported by its association with skin cell differentiation [Jacobs et al., 2013]. A structural variant in the first intron of *BNC2* causes lighter skin color by reducing *BNC2* expression in human melanocytes [Visser et al., 2014, Szpak et al., 2019]. In addition, some haplotypes at *BNC2* have been hypothesized to be introgressed from Neanderthals with high frequency that influences skin pigmentation levels [Visser et al., 2014, Szpak et al., 2019, McArthur et al., 2021]. Prior selection scans suggest that this variation at this gene represents a mode of positive selection termed adaptive introgression [Racimo et al., 2015, 2017, Mughal et al., 2020, Gower et al., 2021], by which selection acts on variants residing on haplotypes that were donated from another species through introgression. Furthermore, the likely importance of *BNC2* in tumor growth has made it a focal point in cancer research [Cesaratto et al., 2016, Wu et al., 2016, Orang et al., 2023], with some cancers linked to alterations in *BNC2* expression and function [Orang et al., 2023, Wu et al., 2016].

Along with the established candidate *BNC2*, *TrIdent* also detected several other cancer-related genes as sweep candidates. Specifically, with a 10-window mean sweep probability of 0.996 we identified the *MTOR* gene (Figure 9E), which produces the protein mTOR that is a critical regulator of cellular growth and survival and that is frequently dysregulated in breast cancer [Miricescu et al., 2020]. Cancer progression and uncontrolled growth of cells are caused by abnormal mTOR activation [Takei and Nawa, 2014, Costa et al., 2015]. *RBMS3* is another detected sweep candidate (10-window mean sweep probability of 0.9969) (Figure 9G) that is linked to tumor suppression in breast cancer [Yang et al., 2018]. Another candidate gene *LAMC2* (10-window mean sweep probability of 0.9999) (Figure 9F), which encodes the gamma-2 subunit of the protein laminin, is commonly upregulated in oral cancer [Nguyen et al., 2017]. Furthermore, the gene *NKAIN2* may inhibit tumors in some cancers [Zhao et al., 2015], which is another candidate identified by our scan with a 10-window mean sweep probability of 0.98 (Figure 9H). A few other intriguing sweep candidates revealed by our scan are *SERINC2* (10-window mean sweep probability of 0.997) (Figure 9I), *KLHL24* (10-window mean sweep probability of 0.995) (Figure 9J), and *MGAT1* (10-window mean sweep probability of 0.92) (Figure 9K), with association studies linking variants in *SERINC2* with alcohol dependence in European women [Zuo et al., 2013, 2014, 2015], mutations in *KLHL24* associated with loss of keratin 14, which provides structural support to epithelial cells [Lin et al., 2016], and a role of *MGAT1* in the development of type 1 diabetes in Europeans [Rudman et al., 2023]. There appears to be a trend in recent machine learning studies where genes associated with deleterious phenotypes, including cancer, are identified as candidate targets of positive selection [Schrider and Kern, 2017, Mughal and DeGiorgio, 2019, Arnab et al., 2023, Amin et al., 2023]. Schrider and Kern [2017] suggested that weakly deleterious alleles may have hitchhiked to high frequency alongside beneficial variants, resulting in the present-day manifestation of problematic traits. This phenomenon could also be explained by the historical advantages conferred by these genes in past environments, which outweighed their negative impacts under contemporary conditions [Di Rienzo and Hudson, 2005, Di Rienzo, 2006]. Such hitchhiking events illustrate the complexity of evolutionary processes and the trade-offs between short-term benefits and long-term consequences in gene selection, with further work needed to fully understand these dynamics and their implications for modern human health.

In the YRI scan, we found that most genomic windows are classified as neutral with a probability threshold of 0.9 for calling sweeps (approximately 95.41%; Table 1). Using a 10-window mean sweep probability of 0.9, we identified 2,145 candidate sweep peaks, which intersected 666 genes. This scan revealed several prominent candidate selection signals at genes supporting some findings from the CEU scan and exposing distinct selection patterns particular to the YRI population. Several genes highlighted as possible adaptive targets in previous studies of the YRI that were also recovered by our scan include *NNT*, *HEMGN*, *SYT1*, *GRIK5*, and *APOL1* (Figure 10) [Voight et al., 2006, Pickrell et al., 2009, Fagny et al., 2014, Pierron et al., 2014, Harris et al., 2018, Mughal et al., 2020, Harris and DeGiorgio, 2020]. We also applied the *TrIdent* regression model to predict selection parameters at reported sweep candidates found in the YRI scan (Table 3).

**Figure 10:**
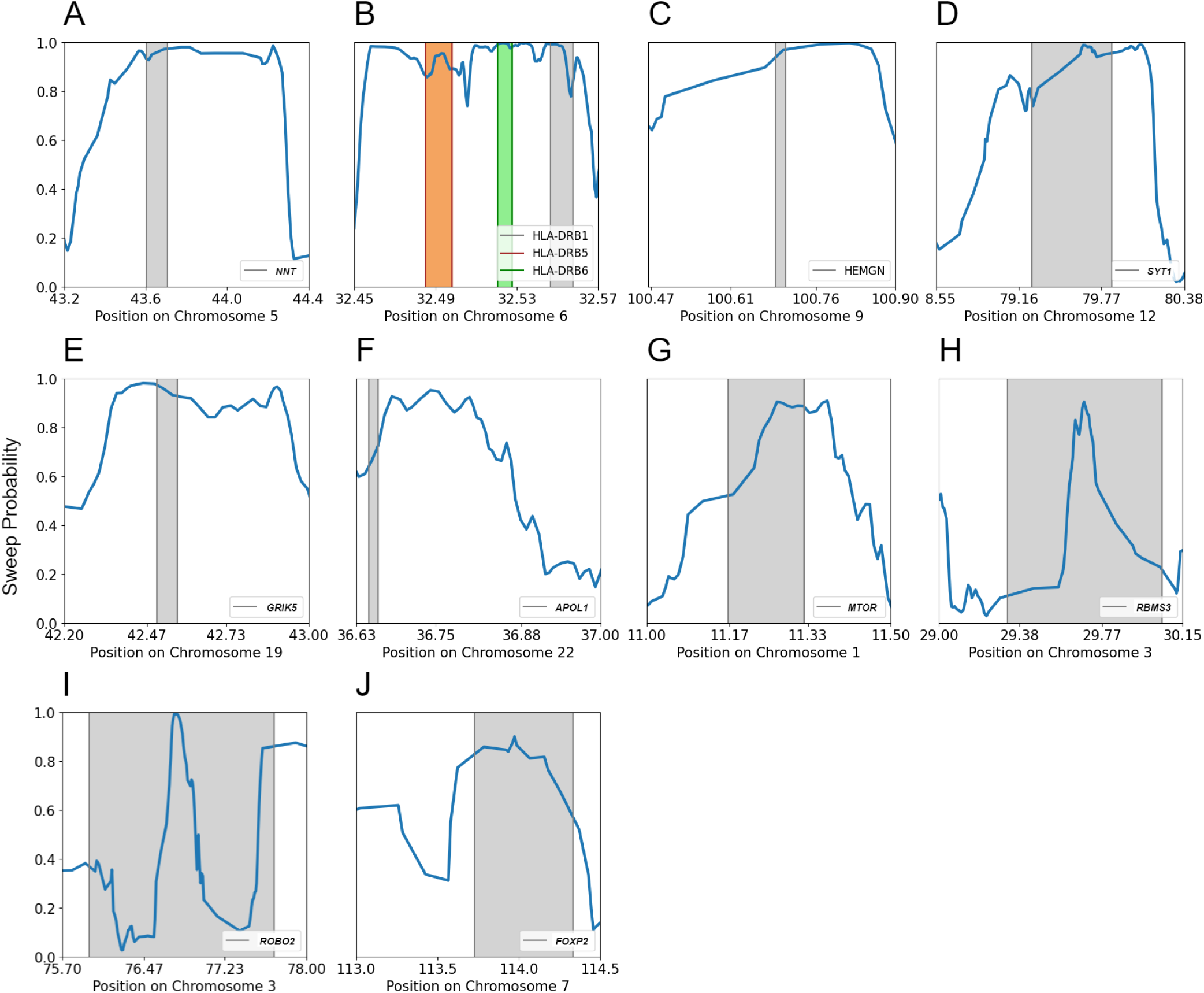
Identified candidate sweep regions from the genome-wide produced using the trained *TrIdent* [*IRV2*] model on the sub-Saharan African (YRI) population in the 1000 Genomes Project dataset. Regions were classified as being under positive selection if they had ten consecutive windows with a sweep probability higher than 0.9. A total of 666 genes across 22 autosomes exhibit qualifying signs of selective sweeps, of which a few of the most interesting candidates are reported here. Figure S22 provides a visual representation of the haplotype diversity surrounding the candidate genes in the plotted panels.

**Table 3:** *TrIdent* regression model inferences for the frequency at which a selected allele became beneficial (*f*), selection strength (*s*), and time at which the beneficial allele reached fixation (*τ*) for genes reported in Figure 9 or the YRI scan.

| Chromosome | $f$ | $s$ | $\tau$ (generations) | Genes |
| --- | --- | --- | --- | --- |
| 1 | 0.0418 | 0.0678 | 1049.3189 | *MTOR* |
| 3 | 0.0097 | 0.0879 | 89.8202 | *RBMS3* |
| 3 | 0.0387 | 0.1033 | 2710.1717 | *ROBO2* |
| 5 | 0.0290 | 0.0579 | 1546.6411 | *NNT* |
| 6 | 0.0832 | 0.0322 | 37.2414 | *HLA-DRB1* |
| 6 | 0.0133 | 0.0323 | 61.7333 | *HLA-DRB5* |
| 6 | 0.0063 | 0.0731 | 69.3420 | *HLA-DRB6* |
| 7 | 0.0011 | 0.0900 | 2901.9021 | *FOXP2* |
| 9 | 0.0200 | 0.0354 | 1883.3812 | *HEMGN* |
| 12 | 0.0222 | 0.0452 | 2024.0176 | *SYT1* |
| 19 | 0.0368 | 0.0478 | 675.6476 | *GRIK5* |
| 22 | 0.0880 | 0.0438 | 2000.9901 | *APOL1* |

The *NNT* gene, located on chromosome 5, exhibits a clear signal with a 10-window mean sweep probability score of 0.989 (Figure 10A). This gene codes for a protein named nicotinamide nucleotide transhydrogenase, which is vital for cellular energy metabolism [Yin et al., 2012, Xiao et al., 2018] and therefore may have possible connection to metabolic adaptations in the YRI population. The *HEMGN* gene on chromosome 9 is another noteworthy candidate, with a mean sweep probability score of 0.974 (Figure 10C) that is involved in hematopoiesis by which the body forms blood cells [Li et al., 2004, Jiang et al., 2010]. Possible responses to environmental variables or pathogenic pressures may have prompted changes in blood cell formation and function. Another strong candidate highlighted by our scan with mean sweep probability of 0.987 is the *SYT1* gene on chromosome 12 (Figure 10D). *SYT1* encodes Synaptotagmin 1, an essential regulator of neurotransmitter release in the nervous system [Xu et al., 2009, Park and Ryu, 2018], and the adaptive relevance of this gene may be connected to cognitive and neurological processes in the YRI population. The *GRIK5* gene on chromosome 19, which encodes a subunit of kainate receptors involved in synaptic transmission and plasticity [Shibata et al., 2006], also shows a prominent signal with a mean sweep probability of 0.991 (Figure 10E). Schrider and Kern [2017] also identified sweep signals at several glutamate receptor genes, highlighting the potential role of neurotransmitter receptors as targets of recent selection in humans. Furthermore, the gene *APOL1* on chromosome 22, which is associated with kidney diseases in individuals of African ancestry [Kruzel-Davila et al., 2017], has been detected with a relaxed qualification criteria by *TrIdent* with a mean sweep probability of 0.801 (Figure 10F).

Within the HLA region of chromosome 6, the *HLA-DRB1*, *HLA-DRB5*, and *HLA-DRB6* genes show strong signals consistent with positive selection (Figure 10B). In particular, the *HLA-DRB6* gene, also identified in our CEU scan, attains a 10-window mean sweep probability of 0.9989. Together with the discovery of noteworthy signals in two neighboring candidate genes (*HLA-DRB1* with a mean sweep probability of 0.963 and *HLA-DRB5* with a mean sweep probability of 0.988), these findings emphasize the ongoing selection pressures to maintain genetic diversity in this chromosomal region. Though *TrIdent* was primarily trained to detect selective sweeps that reduce haplotypic diversity, regions subject to recent balancing selection can sometimes exhibit signatures resembling those of soft or incomplete sweeps [Hermisson and Pennings, 2005, Fijarczyk and Babik, 2015]. This similarity is because balancing selection, particularly when recent or strong, may lead to localized increases in linkage disequilibrium and allele frequency patterns that mimic the sweep-like signals captured by the model.

The signals found by *TrIdent* at *RBMS3* on chromosome 3 and *MTOR* on chromosome 1, identified in both YRI and CEU scans, underscore their broad adaptive significance. The relatively lower mean sweep probabilities in the YRI scan (0.911 for *RBMS3* and 0.906 for *MTOR*) (Figures 10G and 10H) compared to CEU may reflect differences in sweep completion status at these genes between the populations. Additionally, two particularly interesting candidates in the YRI genome are *ROBO2* (mean sweep probability score of 0.9975) on chromosome 3 (Figure 10I) and *FOXP2* (mean sweep probability score of 0.996) on chromosome 7 (Figure 10J). Though *FOXP2* is a classical candidate for selection related to language traits, both *ROBO2* and *FOXP2* are associated with brain development and cognitive processes [Enard et al., 2002, López-Bendito et al., 2007]. Previous studies have hypothesized that these genes might reside in introgression deserts, potentially indicating selective pressure to preserve specific brain functions against introgressed variants from archaic humans that could disrupt these traits [Kuhlwilm, 2018, Buisan et al., 2022]. However, Atkinson et al. [2018] found no evidence of recent adaptation at *FOXP2*, attributing previous signals to sample composition in which an intronic region enriched for sites that are conserved in non-human primates but are polymorphic among humans may be connected to a loss of function in humans. While Atkinson et al. [2018] attributed earlier findings of selection at this locus to sample composition, their methodology may have been underpowered to detect older or subtler sweeps. Flex-sweep [Lauterbur et al., 2023], a modern and sensitive method, also identifies *FOXP2* as a high-confidence candidate for positive selection. Despite the detection of sweep signals at *FOXP2* using *TrIdent*, further investigation may be warranted to reconcile these results and to better understand the evolutionary history of this gene.

To explore whether high-scoring sweep candidates were enriched for specific biological processes, molecular functions, or cellular components, we performed Gene Ontology (GO) analysis using GOrilla [Eden et al., 2009] applied to a single ranked list, where genes were ranked in decreasing order based on their highest 10-window mean sweep probabilities. Significant GO term enrichment was selected based on false discovery rate adjusted *q*-values of less than 0.05. The GO analysis of our CEU scan (Tables S3 and S4) identified several significant biological process terms, including regulation of cell migration, regulation of cell motility, and positive and negative regulation of locomotion, consistently involving key genes *MTOR* and *LAMC2* that we highlighted earlier. Both genes are critical in pathways associated with cell movement and localization, and their dysregulation may contribute to cancer progression through mechanisms like abnormal cellular migration, localization, and growth [Kariya and Miyazaki, 2004, Stipp, 2010, Zoncu et al., 2011, Masuda et al., 2012, Saxton and Sabatini, 2017]. Additionally, the GO analysis highlighted an enrichment of genes involved in essential cellular components, especially those associated with the plasma membrane and cytoskeleton, which are central to cellular structure and movement [Alberts et al., 2002, Fletcher and Mullins, 2010, Keren, 2011, Jacobson et al., 2019]. Alterations in these components can facilitate uncontrolled cell motility and invasion seen in metastatic cancer [Condeelis and Pollard, 2006, Yilmaz and Christofori, 2009]. The identified cellular component GO terms also include laminin complex and cell projection part that are associated with *LAMC2*. *LAMC2* helps encode the laminin complex protein that supports cell growth, motility, and adhesion, essential processes that are often disrupted in cancer progression. In contrast, the YRI GO analysis did not yield significantly enriched terms, indicating that, unlike the CEU population, the sweep candidate genes in the YRI population do not appear to center on specific molecular functions, cellular components, or biological processes to the same extent.

## Discussion

Our results show that the *TrIdent* [*IRV2*] model demonstrates strong performance in detecting selective sweeps, outperforming *T-REx*, a technique that also uses images of haplotype variation as input, in both accuracy and power. *TrIdent* [*IRV2*] effectively captured the complex genetic patterns suggestive of selective sweeps through the utilization of the pre-trained *InceptionResNetV2* architecture, which is known for its sophisticated deep feature extraction prowess. Due to overall higher accuracy and power, we adopted the *Trident* [*IRV2*] model over four alternative *TrIdent* models. The complex architecture of *InceptionResNetV2* allows *TrIdent* [*IRV2*] to effectively identify signals of selection from relatively small amounts of data that may be missed by simpler models. Moreover, the use of a pre-trained deep learning model as an upstream feature extractor in *TrIdent* [*IRV2*] offers practical benefits beyond just performance, including reduction in time and resources required for model training.

A key strength of *TrIdent* is its ability to achieve strong classification performance even when trained with a relatively small dataset, with the original implementation requiring only 1,000 samples per class. Notably, increasing the training data size does not yield substantial improvements in performance, suggesting that *TrIdent* effectively captures the relevant patterns with limited data. For the CEU test case, increasing the training set to 5,000 samples per class results in only a marginal 0.4% improvement in classification accuracy, with a further increase to 10,000 samples per class yielding just an additional 0.2% gain (Figure S13). Similarly, for the YRI test case, increasing the training set to 5,000 samples per class improves accuracy by only 0.2%, and expanding it to 10,000 samples per class provides no further improvement in classification accuracy. These findings indicate that *TrIdent* reaches near-optimal performance with relatively small training datasets, making it a highly data-efficient approach for sweep detection.

The differences between the image generation strategies of *TrIdent* and *T-REx* lie in several key aspects. First, *T-REx* processes an entire 1.1 Mb region by sorting haplotypes in windows of 100 SNPs with a stride of 10 SNPs, whereas *TrIdent* focuses on a more fine-grained approach, using windows of 25 SNPs with a stride of two SNPs across a region of 499 SNPs. Second, at the SNPs where different windows overlap, *T-REx* averages the minor allele count at these SNPs after performing an *L*_1_-norm-based haplotype rearrangement. To accomplish this task, *T-REx* first stores the windows, rearranges them, and then calculates the intersections between windows to average the minor allele counts at these intersections. This process is meant to smooth transitions between windows and retain information, especially useful for detecting weak or old sweeps. In contrast, *TrIdent* follows a more streamlined approach, which sequentially fills the input image matrix by calculating minor allele counts and sorting these counts directly within windows, avoiding the need to store windows or recalculate quantities at window intersections. This procedure not only allows *TrIdent* to bypass the additional space overhead required by *T-REx*, it also allows for faster processing times and based on the presented results, is a more computationally efficient set of operations that does not compromise the detection of selection signals. Our image generation approach therefore prevents the need for extra manipulation of genomic images, which can lead to the emergence of summary statistics and poor generalization [Cecil and Sugden, 2023].

In contrast, when comparing *TrIdent* to shallow CNN-based methods that use images of summary statistic variation as input (*e.g.*, diploS/HIC), we found that *TrIdent* trained with native images underperformed relative to these competitors. However, the nature of input data explains this disparity. In particular, diploS/HIC benefits from the usage of relatively shallow CNNs specifically trained on structured matrices of summary statistics, yet *TrIdent* outperforms diploS/HIC when trained on images of the same summary statistics. This performance boost highlights the advantage *TrIdent* gains from utilizing the *InceptionResNetV2* architecture, which excels in extracting complex patterns from images. Unlike the shallower CNNs in diploS/HIC, *InceptionResNetV2* combines both deep and wide feature extraction pathways, enabling *TrIdent* to capture genomic variation patterns in a richer and more nuanced manner.

While diploS/HIC is a widely recognized strong performer in detecting sweeps, newer and more powerful summary statistic-based tools like Flex-sweep [Lauterbur et al., 2023] have demonstrated further improvement in detection accuracy. Flex-sweep not only employs a broader set of advanced summary statistics than diploS/HIC but also incorporates custom-designed statistics that have been shown to be highly effective. A key advantage of Flex-sweep lies in its unique approach to summary statistic computation as it computes nine out of its 11 summary statistics within each window at five different nested scales. This multi-scale approach allows Flex-sweep to capture genomic variation patterns more comprehensively, leading to superior performance. Despite both methods utilizing similar CNN-based architectures, the enhanced information content within the summary statistic images used by Flex-sweep likely explains its improved accuracy. Furthermore, our results with *TrIdent* [*IRV2*, *SS*] reinforce the idea that summary statistic-based images, when carefully designed for pre-trained models, can significantly boost performance. Though *TrIdent* integrates both image generation and feature extraction-prediction methodologies, users can selectively deploy either component depending on their analytical goals, offering flexibility in genomic sweep detection tasks.

In analyzing the GradCAM-based class activation maps generated from *TrIdent* [*IRV2*], we observed a consistent focus on the lower-middle regions of the images, which corresponds to the difference of high minor allele count pattern between sweep and neutral replicates. Interestingly, a slight asymmetry in the gradient patterns was noted, with the right side of the lower-center region appearing more emphasized than the left. We hypothesize that this asymmetry may arise from redundancy in the pixel patterns across the two halves of the center pattern. Specifically, the pattern on the right side could render the left side redundant due to the regularization penalty applied during the training of the logistic regression model. To investigate this hypothesis, we horizontally flipped the input images from the original training datasets and recomputed the GradCAM maps to generate the mean heatmaps. Despite this transformation, the resulting maps continued to emphasize the right side of the lower-center pattern (Figure S14). This consistent focus, irrespective of flipping, suggests that the model identifies key features in these regions that are central to distinguishing sweeps from neutrality, even if one half of the central change in gradient renders the other half marginally redundant.

To evaluate the robustness of *TrIdent* in detecting sweeps across different selection strengths, we analyzed its performance across varying selection coefficient ranges. Because the original implementation of *TrIdent* [*IRV2*] was trained on sweeps with selection coefficients within the interval [0.005, 0.1], it predictably struggled to detect sweeps in the weaker selection range of [0.001, 0.005], as these values fell outside the training distribution. Specifically, sweep classification accuracy in this range was 59.15% for the CEU test case and 63.3% for the YRI test case (Figure S15). To explore whether we can improve generalizability, we trained new *TrIdent* [*IRV2*] models on datasets where selection coefficients were drawn from the broader range of [0.001, 0.1], sampled uniformly on a logarithmic scale. This adjustment caused a slight decrease in performance on the original test dataset. Detection of sweeps in the weaker selection range of [0.001, 0.005] for the CEU test case was increased by only around 3%, yet the YRI test case improved by almost 7% (Figure S16; right panels). While this analysis demonstrates that the performance of *TrIdent* in detecting weaker sweeps is contingent upon the selection regime used during training, we hypothesize that we may need to subtrantially increase training samples to further improve generability of *TrIdent* in regards to detecting both stronger and weaker sweeps.

Low-coverage data presents significant challenges for genomic analysis due to the uncertainty in determining genotypes [Nielsen et al., 2011, Fumagalli, 2013, Korneliussen et al., 2014]. This uncertainty stems from various sources, including mapping errors, sequencing errors, and the random sampling of haploid reads from a diploid genotype, making it difficult to accurately infer the underlying genetic variation. In genomic regions of low coverage, the detection of selective sweeps is particularly problematic, as the likelihood of missing key genetic variants increases, leading to incomplete or inaccurate genomic signals that may masquerade as sweeps. *TrIdent* may be well-suited for applications to low-coverage data, as it relies on windows of minor allele counts and works with unphased multilocus genotype data. These minor allele counts could be approximated with expected allele counts from genotype likelihoods [Fumagalli et al., 2014, Fox et al., 2019], which would permit *TrIdent* to account for the uncertainties in calling genotypes from low-coverage sequencing and has been shown to be a suitable approach for similar settings [Gower et al., 2021]. Finally, using minor allele counts within *TrIdent* input images is advantageous as it does not require a reliable outgroup sequence to polarize alleles as derived or ancestral. This characteristic is important, as erroneous allele polarization can alter the distribution of allele frequencies to be skewed toward high-frequency derived alleles, much like that of a selective sweep [Hernandez et al., 2007].

To expand upon how our local sorting of haplotypes influences model performance, we explored its benefits compared to a global sorting approach, which we denote as *TrIdent* [*IRV2*, *global*]. The image generation process of *TrIdent*, as described in the *Image Generation* subsection of the *Methods*, arranges haplotypes based on minor allele counts independently within small sub-windows, preserving localized structural patterns critical for classification. In contrast, global sorting organizes the entire 499-SNP window at once based on total minor allele counts. While global sorting guarantees that every pixel value in a row of the resulting image corresponds to the same haplotype, this uniform arrangement may obscure subtle, localized patterns essential for distinguishing sweeps from neutrality. Our results demonstrate that local sorting significantly improves predictive accuracy, with global sorting resulting in accuracy losses of 9.25% in the CEU dataset and 6.1% in the YRI dataset (Figure S17). The larger effect in the CEU population, which has experienced a bottlenecked demographic history, highlights the critical role of local patterns that are more susceptible to being blurred by global sorting. By preserving the fine-grained haplotype structures within sub-windows, local sorting enables *TrIdent* to capture nuanced signals associated with selective sweeps, especially in populations with complex evolutionary histories.

In our efforts to bridge the gap between the highly complex deep CNN architecture of *InceptionResNetV2* and comparatively shallower sequential CNN architecture used in *smbCNN* (which is identical to the diploS/HIC architecture), we examined a custom designed residual [He et al., 2016] and multi-path [Szegedy et al., 2017] CNN architecture that we term *scCNN* (shallow complex CNN) for ease of reference. By incorporating advanced architectural features, *scCNN* enhances complex feature extraction and maintains training efficiency, making it well-suited for complex tasks without the need for excessively deep networks like *InceptionResNetV2*. The architecture of *scCNN* is detailed in *Constructing the scCNN architecture* in *Methods*. Despite the structural complexities introduced in *scCNN* compared to *smbCNN*, *scCNN* achieved accuracies of 84.10% on the CEU dataset and 86.75% on the YRI dataset (Figure S18). The *scCNN* model still falls short of the performance levels achieved by *TrIdent* [*IRV2*] with its pre-trained *InceptionResNetV2* backbone. The reason lies in the inherent strengths of the *InceptionResNetV2* architecture, which benefits from extensive pre-training on large-scale datasets and a more sophisticated multi-path design that can capture a broader range of features across different scales. While this *scCNN* model represents a marginal improvement over the simpler *smbCNN* architecture, it underscores the superior capability of the *InceptionResNetV2* architecture employed by *TrIdent* [*IRV2*] in detecting subtle patterns of selection in our haplotype re-arrangement based images.

Though *TrIdent* achieved good performance on demographic settings with fluctuating populations, it is also important to consider other, more complex, demographic factors that could impact method performance, such as significant population substructure and admixture. These factors, particularly the timing and scale of migration events, can strongly influence the observed spatial haplotypic diversity and may result in misleading indicators of selection [Harris et al., 2018]. The mean of the sweep images used by *TrIdent* (Figure S1) reveals a funnel-shaped dark region at the selected locus (center columns of the image), indicating low pixel intensity values that are absent in the mean of the neutral images. Class activation maps (Figure 7) also show that our *TrIdent* focuses on the lower-middle input image pixels for classification. Now, the image generation process (Figure 1) illustrates that pixels at the top of the images represent haplotypes with low minor allele counts resulting in zero or near-zero values, whereas pixels toward the bottom of the images reflect haplotypes with higher minor allele counts resulting in higher values that can reach values up to 25 (*Modeling Description*). This observation suggests that the loss of haplotype diversity near sweep loci leads to more near-zero values in the middle-to-lower parts of the image, which *TrIdent* uses for classification. Therefore, any form of migration or population structure introducing near-zero minor allele counts in this region could be misclassified as a sweep, whereas higher values could obscure sweep signals. Migration from a population with a smaller effective size can introduce haplotypes with limited diversity to the recipient population. As a consequence, high migration rate settings may replace much of the haplotypic variation in the recipient with this low haplotype diversity from the donor, leading many polymorphic sites with low minor allele counts, thereby increasing the occurrence of near-zero values in the middle-to-lower sections of image matrices and potentially causing false inferences of sweeps. On the other hand, such migration from a population with a moderate to large effective size after the beneficial allele has swept can mask sweep footprints by replacing the low haplotypic diversity with moderate to high levels from the donor population, resulting in higher pixel values in the middle-to-lower image regions. Therefore, if it is expected that the study population may have received significant gene flow from other populations, then accounting for such processes when generating training datasets is likely important to guard against both false positive and false negative results when detecting sweeps.

Furthermore, we conducted additional analyses to assess the impact of sweep shoulders on the classification performance of *TrIdent*. Specifically, we evaluated the ability of *TrIdent* to detect sweeps when the beneficial mutation was positioned at different locations within a 1.1 Mb sequence. By centering sweeps at 500, 375, 250, 125, and zero kb away from the center, we found that *TrIdent* gradually loses detection ability when the sweep moves farther off-center for both CEU and YRI test cases (Figure S19; left panel). For example, when the sweep is positioned 125 kb away from the image center, sweep detection accuracy is approximately 78%, but performance predictably drops substantially at greater distances. To further investigate how sweep probability signals diffuse across the genomic region, we conducted a scan across the simulated sweep sequences, protocol of which is detailed in the *Assessing sweep shoulders* subsection of the *Methods*. This analysis enabled us to systematically assess how predicted sweep probabilities change with distance from the beneficial mutation (Figure S19; right panel), demonstrating that while *TrIdent* can detect selective sweeps beyond a perfectly centered mutation, its accuracy declines as the focal mutation shifts further from the image center, emphasizing the importance of generating images in small genomic distances when analyzing empirical data.

The pre-trained *InceptionResNetV2* model is especially suitable for assessing the image of multilocus variation used by *TrIdent* because of its hybrid architecture, which effectively integrates the advantageous qualities of both Inception and Residual networks. Inception modules facilitate the extraction of features at several scales, enabling the detection of patterns of different sizes and complexities in the input images. The added residual connections enhance the feature extraction process by utilizing network depth, as model layers have the ability to capture diverse sets of features, which allows for accurate identification of small changes in genomic variation with multi-level inspection and concatenation of information from input images without the need for further training. The promising performance of *TrIdent* on detecting sweeps when confronted with common empirical hurdles while also having the capacity to predict evolutionary parameters underlying sweeps makes it a versatile tool for population genetic studies. This versatility is underscored by consistently good performance of *TrIdent* across two demographic histories, showing marginally higher accuracy in the YRI dataset due to greater genetic diversity, whereas on the CEU dataset, affected by a recent severe bottleneck, exhibited reduced background variation that slightly diminished sweep detection accuracy. The robust performance of *TrIdent* when faced with non-sweep patterns that can masquerade as sweeps, such as technical issues that lead to large missing genomic segments and the process of background selection further underscores its ability to handle challenging genomic scenarios effectively. Missing data, such as undiscovered polymorphism due to poor mappability, can mimic selective sweeps by reducing local haplotypic diversity. Despite this challenge, *TrIdent* maintains high and only marginally-decreased accuracy under such settings. Additionally, *TrIdent* shows resilience against misleading patterns of lost diversity due to background selection. By analyzing haplotype distributions, the model can effectively guard against false classification of background selection as a selective sweep while maintaining a low false positive rate.

Though our empirical filtration method, as detailed in the *Filtration of empirical data and empirical image generation* subsection of the *Methods*, is effective in reducing false discoveries, it can also inadvertently exclude some true signals from the analysis. A notable case study of this phenomenon is the *ACKR1* gene (Atypical Chemokine Receptor 1), also known as the *DARC* gene (Duffy Antigen Receptor for Chemokines), which has been widely recognized as a target of positive selection due to its role in conferring resistance to malaria caused by *Plasmodium vivax* infection [Horuk, 2015, Yin et al., 2018]. This highly recombining locus poses a particular challenge for detecting selection signals due to its complex genetic landscape [McManus et al., 2017, Lauterbur et al., 2023]. In our analysis, *ACKR1* was excluded during the filtration process because it exhibited a low mean CRG score. To assess whether this exclusion impacted the detection of a true signal at this locus, we reanalyzed SNPs within and around *ACKR1*. This reanalysis revealed evidence of a sweep in the YRI scan, with a mean peak probability of 0.858 (Figure S20). However, no similar signal was detected in the CEU scan. This example illustrates the trade-offs inherent in applying stringent filtration criteria and highlights the importance of carefully balancing false discovery control with the retention of biologically meaningful signals.

Finally, we scanned two human populations (CEU and YRI) for sweeps using *TrIdent*, and reported candidate genes with previous literature support, as well as some novel candidate loci. However, it is also important to highlight those candidates that *TrIdent* identified with highest confidence. We therefore considered the 10 sweep candidates (Tables S1 and S2) with highest 10-window mean sweep probabilities within each population to better compare and contrast the genomic regions strongly favored by *TrIdent*. Two genes (*LAMC2* and *MTOR*) in our CEU analysis and two genes (*HLA-DRB6* and *ROBO2*) in our YRI analysis are among the top 10 candidates in their respective scans. By focusing on these top 10 candidates, which have no overlap between the pair of sweep scans, we may gain insights into the key genetic adaptations and distinct evolutionary pressures that have shaped these populations. Whereas the CEU candidates reflect adaptations potentially linked to metabolism, neurodevelopment, and cancer-related pathways, those from the YRI focus more on immune function and pathogen resistance. This potential functional divergence highlights how different environments and historical pressures may have shaped the genetic landscape of these populations in unique ways. However, the exclusivity of candidates may also suggest a limitation of the trained *TrIdent* model, in which it may have reduced capacity for detecting more ancient sweeps that would have occurred prior to the split of CEU and YRI, thus missing shared sweep signals across these populations. Future studies could train *TrIdent* models on variation across multiple populations to detect shared sweeps and discriminate them from population-specific events.

## Methods

### Simulation protocol

To simulate replicates under both the European and sub-Saharan African human demographic histories, we drew the per-site per-generation mutation rate *µ* uniformly at random within the interval [2.21 × 10*^−^*^9^, 2.21 × 10*^−^*^8^] with a mean of 1.21 × 10*^−^*^8^ [Scally and Durbin, 2012, Schrider and Kern, 2017] and the per-site per-generation recombination rate *r* at random from an exponential distribution with mean 10*^−^*^8^ [Payseur and Nachman, 2000, Schrider and Kern, 2017] and truncated at three times the mean [Schrider and Kern, 2017]. For each replicate simulated under the inferred demographic history [Tennessen et al., 2012], we sampled 198 haplotypes for CEU and 216 haplotypes for YRI of length 1.1 megabases (Mb) to match the number of sampled haplotypes in our empirical experiments.

We generated sweep simulations with per-generation selection strength *s* acting on a beneficial allele at frequency *f* that arose at a locus at the center of a simulated genomic region, with *s* drawn uniformly at random within the interval [0.005, 0.1] and *f* drawn uniformly at random on a logarithmic scale within the interval [1*/*(2*N_e_*), 0.2] [Schrider and Kern, 2017], where *N_e_* is the present-day effective population size. Moreover, we also drew the number of generations *τ* in the past at which the beneficial allele reached fixation uniformly at random within the interval [0, 2000] [Schrider and Kern, 2017]. This simulation protocol guarantees that both training and test sets reflect an array of sweep settings, including weaker (small *s*) and stronger (large *s*) selection scenarios, harder (small *f*) and softer (large *f*) sweeps, and time *τ* in which mutation, recombination, and genetic drift can erode signals of sweeps, all of which affect the size and prominence of the genomic footprint left by a sweep after fixation. These modeled genetic, demographic, and selection parameters are expected to lead to significant overlap in the distributions of genetic variation between sweeps and neutrality.

### Image generation

For each simulated replicate, we retain only bi-allelic SNPs with a minor allele count of at least three (*i.e.*, removed singletons and doubletons). We then generate a matrix **M** of dimension *n*×499, where *n* denotes the number of haplotypes in a replicate, with each row representing one of the sampled haplotypes and each column representing one of 499 single nucleotide polymorphisms (SNPs), where these SNPs were chosen as the central position, 249 closest SNPs upstream of the central position, 249 closest SNPs downstream of the central position of the simulated genomic region, which is where the selected mutation is introduced in sweep simulations. The element **M***_ij_* takes a value of zero if the haplotype within the *i*th row has the major allele at the SNP in the *j*th column, and otherwise has the value of one for the minor allele. From this matrix, we sought to create a representation of genome variation that might make pattern recognition easier by generating a matrix **X** of dimension *n*×237 computed using windows of 25 SNPs with a stride of two between each window, with element values **X***_ij_* ∈ {0, 1*, … ,* 25}. That is, for *j* ∈ {1, 2*, … ,* 237}, we computed the minor allele counts for each of the *n* rows of a genomic window of length 25 SNPs starting at column 2*j* − 1 of **M**, arranged these minor allele counts values in increasing order, and set the values in column *j* of **X** as this sorted list of minor allele counts. This procedure summarizes the number of minor alleles within a given 25-SNP window for each haplotype, and thus rows toward the top of **X** summarize haplotypes with a greater number of major alleles, whereas rows toward the bottom have a greater number of minor alleles. We then perform the same computation of each 25-SNP window by taking a stride of two SNPs. This matrix is subsequently resized using linear interpolation, depending on the input image size requirements of the pre-trained neural network architectures. For example, pre-trained models like *InceptionResNetV2* require input image sizes of 299 × 299, whereas for some pre-trained models, such as *VGG16* or *MobileNetV2*, the required input image size is 224 × 224. An example illustrating this image generation procedure is presented in Figure 1 (top panel).

### Constructing the *scCNN* architecture

The *scCNN* architecture begins with an initial convolutional layer that applies a set of filters to the input image. Following this layer, the model incorporates a residual block that contains two convolutional layers, each followed by batch normalization [Bjorck et al., 2018] and ReLU activation [Krizhevsky, 2012]. A skip connection within the residual block bypasses these two convolution layers, directly adding the input to the next convolution layer, which maintains gradient flow during training and enables deeper feature learning [He et al., 2016]. There is an additional skip connection before the fifth convolution layer that allows the concatenation of fourth and fifth convolution layer outputs that are then fed into the dense layer. The model is optimized using the Adam optimizer and binary cross-entropy loss, ensuring efficient training and robust performance.

### Computational resources and requirements

Our analysis was conducted on a system equipped with an AMD EPYC 7702 64-core CPU with 100 GB of RAM. The *TrIdent* image generation method requires a mean of 6.83 seconds to generate an image from a simulated replicate. Loading the *InceptionResNetV2* architecture with its pretrained weights uses approximately 513 MB of RAM. The mean time to compute the GAP layer output using *InceptionResNetV2* is 0.018 seconds for a single input image. Training the penalized logistic regression model on 2,000 images with hyperparameter tuning, if run parallelly, requires approximately six minutes and 30 seconds and consumes 56 MB of memory.

Conversely, training the entire *InceptionResNetV2* architecture, as executed in the training of the *IRV2* model, necessitates additional resources beyond those previously mentioned. Specifically, training *IRV2* on 2,000 images, incorporating early stopping to prevent overfitting, attained convergence in 37 epochs for the CEU dataset and 26 epochs for the YRI dataset. Each epoch required approximately four minutes to complete and utilized around 26 GB of RAM in total for both test cases. We anticipate that as more training images are used, the RAM utilization and the training time would increase substantially for the full *IRV2* model.

For comparison, we also assessed the computational resources required by diploS/HIC. Extracting 101 summary statistics from each simulated replicate takes on average 17.17 seconds. Training the CNN model with early stopping, using a dataset consisting of summary statistics from 1,000 replicates per class, allows diploS/HIC to converge in 13 epochs, requiring approximately 42 minutes and consuming 7.5 GB of RAM.

### Application of CLUES2

To benchmark the accuracy and precision of the *TrIdent* [*IRV2*, *ANN*] nonlinear regression model in estimating selection coefficients, we compared its results against CLUES2 [Vaughn and Nielsen, 2024]. First, we used the ms2vcf tool of the coatli [Klassmann, 2013] package to convert discoal output ms formatted files to VCF format, as SINGER [Deng et al., 2024], one of the two software supported by CLUES2 to compute ancestral recombination graphs (ARGs), accepts VCF files as input. Because the exact mutation rate applied in each simulated replicate is unavailable, we used a mean mutation rate of 1.21 × 10*^−^*^8^, consistent with our simulations. Additionally, as we could not extract the recombination to mutation rate ratio for each replicate, we set this parameter to the default value of one in SINGER. We also retained the default number of posterior samples of 100. After generating ARGs, we used the SingerToCLUES pipeline of CLUES2 to convert the output from SINGER to the required CLUES2 input format. For the genomic position of the beneficial mutation, we selected the SNP closest to the simulation center. Finally, to run the CLUES2 inference script, we used the sample derived allele frequency of this central SNP. For the maximum time (in generations) to be considered in the analysis, we chose 2,000 generations in the past. We also tested 1,000 and 4,000 generations, but the results remained largely unchanged.

### Assessing sweep shoulders

To analyze classification performance of *TrIdent* on sweep shoulders, for both CEU and YRI datasets, we computed the mean sweep probability across 1,000 test images within overlapping 20 kb bins spanning the 1.1 Mb simulated region. Using a stride of 10 kb, we created 48 bins with the first bin spanning 300–320 kb to last bin spanning 780–800 kb. We exclude the first and last 300 kb of the region because the image generation process requires at least 499 SNPs with the center SNP falling within a bin. When a bin is positioned too far from the simulation center (550 kb), the number of qualifying images meeting these criteria becomes insufficient for reliable analysis. To generate images for this experiment, we scanned the entire 1.1 Mb sweep replicates in an identical manner to how images were generated from chromosomes in our empirical analysis, as described in the *Filtration of empirical data and empirical image generation* subsection. Once generated, images were assigned to bins based on their center SNP positions. Any image with a center SNP falling outside these predefined bins was excluded from further analysis.

### Filtration of empirical data and empirical image generation

To apply *TrIdent* to the phased genotype data from the 99 CEU and 108 YRI individuals of the 1000 Genomes Project dataset [1000 Genomes Project Consortium, 2015], we retained only bi-allelic SNPs with a minor allele count of at least three (*i.e.*, removed singletons and doubletons). Following the protocol of Mughal et al. [2020], we further filtered this dataset by excluding genomic segments of length 100 kb with mean CRG (Consensus Reference Genomes) mappability and alignability scores [Talkowski et al., 2011] below 0.9 to reduce the possibility of misleading signals due to technical concerns. CRG mappability scores reflect the probability of accurately mapping short sequencing reads to a specific genomic region. We then created images for input to *TrIdent* by considering the first 499 contiguous SNPs on an autosome, converting the haplotype variation across these SNPs into an image according to the procedure outlined in the *Image generation* subsection, processing subsequent images by moving the SNP window by a stride of two SNPs along the autosome, and repeating this procedure for all autosomes. The chromosomal location of each observation was set as mean position of the 249th and 250th SNP.

## Acknowledgments

This work was supported by National Institutes of Health grant R35GM128590, by National Science Foundation grants DBI-2130666, DEB-1949268, and BCS-2001063, and by the UKRI Natural Environment Research Council grant NE/Y003519/1. Computations for this research were performed using the services provided by Research Computing at the Florida Atlantic University.

## Data Availability

We release the source code for *TrIdent* under the MIT open source license, and this repository can be accessed on GitHub (https://github.com/sandipanpaul06/TrIdent/). The CEU and YRI data from the 1,000 Genomes Project can be accessed from the project website (https://www. internationalgenome.org/category/phase-3/).

## Notes

### Competing Interest Statement

The authors have declared no competing interest.

