## Supplemental Figures and Tables for "Efficient detection and characterization of targets of natural selection using transfer learning"

### Supplementary material

Table S1: Top 10 sweep candidates in the CEU scan.

| Rank | Gene | Chromosome | Sweep Probability |
|---|---|---|---|
| 1 | *LAMC2* | 1 | 0.999929 |
| 2 | *MTOR* | 1 | 0.996903 |
| 3 | *SLC22A23* | 17 | 0.999170 |
| 4 | *MOXD1* | 6 | 0.996720 |
| 5 | *RBFOX2* | 22 | 0.996598 |
| 6 | *CRIP1* | 14 | 0.996533 |
| 7 | *RPSAP52* | 12 | 0.996452 |
| 8 | *SMURF2* | 17 | 0.996008 |
| 9 | *SDK1* | 7 | 0.995703 |
| 10 | *FHIT* | 3 | 0.995284 |

Table S2: Top 10 sweep candidates in the YRI scan.

| Rank | Gene | Chromosome | Sweep Probability |
|---|---|---|---|
| 1 | *HLA-DRB6* | 6 | 0.998939 |
| 2 | *NANS* | 9 | 0.998311 |
| 3 | *CATSPERG* | 19 | 0.998290 |
| 4 | *LRRFIP1* | 2 | 0.998174 |
| 5 | *HIF1AN* | 10 | 0.997970 |
| 6 | *SEC31B* | 10 | 0.997898 |
| 7 | *ROBO2* | 3 | 0.997526 |
| 8 | *ITGAE* | 17 | 0.997499 |
| 9 | *TUBE1* | 6 | 0.997198 |
| 10 | *NDUFB8* | 10 | 0.996982 |

Table S3: Gene Ontology (GO) Biological Process for CEU genes ranked based on *TrIdent* prediction output.

| GO Code | Description | $P$-value | Genes |
|---|---|---|---|
| GO:0030334 | Regulation of cell migration | $4.12 \times 10^{-7}$ | *SEMA3C*, *LIMCH1*, *PTPRK*, *FGR*, *CD300A*, *FRMD5*, *GCNT2*, *CLASP1*, *IL34*, *LRIG2*, *C5AR1*, *SMURF2*, *PODN*, *ATP8A1*, *RHOJ*, *ITGB1BP1*, *STARD13*, *RIN2*, *PPARD*, *ZMYND8*, *FGF1*, *SLC8A1*, *DACH1*, *LAMC2*, *ROCK1*, *OSGIN1*, *NISCH*, *BCAS3*, *PTPRT*, *MMP28*, *MTOR*, *DNM1L*, *SEMA3G* |
| GO:2000145 | Regulation of cell motility | $4.18 \times 10^{-7}$ | *SEMA3C*, *LIMCH1*, *PTPRK*, *FGR*, *CD300A*, *FRMD5*, *GCNT2*, *CLASP1*, *IL34*, *LRIG2*, *C8orf44-SGK3*, *C5AR1*, *SMURF2*, *PODN*, *ATP8A1*, *MCU*, *PIK3CD*, *SPOCK2*, *KIT*, *RHOJ*, *ITGB1BP1*, *STARD13*, *RIN2*, *PPARD*, *GAS2L2*, *ZMYND8*, *FGF1*, *SLC8A1*, *DACH1*, *LAMC2*, *ROCK1*, *OSGIN1*, *NISCH*, *BCAS3*, *PTPRT*, *MMP28*, *MTOR*, *DNM1L*, *SEMA3G* |
| GO:0051270 | Regulation of cellular component movement | $1.14 \times 10^{-6}$ | *SEMA3C*, *LIMCH1*, *PTPRK*, *FGR*, *CD300A*, *FRMD5*, *GCNT2*, *CLASP1*, *IL34*, *ATP1A1*, *LRIG2*, *C5AR1*, *SMURF2*, *PODN*, *ATP8A1*, *SPOCK2*, *RHOJ*, *ITGB1BP1*, *STARD13*, *RIN2*, *PPARD*, *GAS2L2*, *ZMYND8*, *FGF1*, *SLC8A1*, *DACH1*, *LAMC2*, *ROCK1*, *OSGIN1*, *NISCH*, *BCAS3*, *PTPRT*, *MMP28*, *MTOR*, *DNM1L*, *SEMA3G* |
| GO:0040012 | Regulation of locomotion | $2.13 \times 10^{-6}$ | *SEMA3C*, *LIMCH1*, *PTPRK*, *FGR*, *CD300A*, *FRMD5*, *GCNT2*, *CLASP1*, *IL34*, *LRIG2*, *C5AR1*, *SMURF2*, *PODN*, *ATP8A1*, *SPOCK2*, *RHOJ*, *ITGB1BP1*, *STARD13*, *RIN2*, *PPARD*, *GAS2L2*, *ZMYND8*, *FGF1*, *SLC8A1*, *DACH1*, *LAMC2*, *ROCK1*, *OSGIN1*, *NISCH*, *BCAS3*, *PTPRT*, *MMP28*, *MTOR*, *DNM1L*, *SEMA3G* |

| | | | |
|---|---|---|---|
| GO:0032879 | Regulation of localization | $6.18 \times 10^{-6}$ | *TACR1*, *P2RY6*, *ITPR1*, *ABCA1*, *TRAPPC12*, *ABCA3*, *ABL1*, *RAB3GAP1*, *PARK2*, *ASIC2*, *RHOJ*, *KCNC3*, *KCND3*, *GAS2L2*, *ZMYND8*, *BRAF*, *SPAG9*, *SCN4A*, *KCNJ6*, *PLA2G6*, *MEGF8*, *LIMCH1*, *CCL5*, *KCNQ1*, *MSR1*, *PLXNA4*, *C5AR1*, *ADCY8*, *KIT*, *DNAJC1*, *SEMA4A*, *MTUS1*, *RIN2*, *CNPY2*, *DHRS7C*, *PTPN6*, *HECW2*, *ENG*, *RAB17*, *CACNA1A*, *PTPRK*, *PTPRM*, *SFRP1*, *ADRBK1*, *PTPRO*, *CACNB4*, *CRY1*, *CACNG2*, *GCNT2*, *DNAJC27*, *CORO2B*, *PODN*, *ATP8A1*, *ATG3*, *SPOCK2*, *MX2*, *CAMK2G*, *LAMA3*, *LAMB1*, *LILRA2*, *RAB31*, *NECAB2*, *LAMC2*, *LCK*, *LYPLA1*, *CASQ2*, *GAB2*, *C2CD5*, *MYLK*, *ALOX12*, *FRMD5*, *REEP6*, *DPP10*, *RAPGEF4*, *SLC2A2*, *LRIG2*, *EEF2K*, *MAGI2*, *PARD3*, *CTNNA2*, *PPP1R12A*, *SMURF2*, *PIK3CD*, *PIK3R1*, *ITGB1BP1*, *TRPV2*, *GNA12*, *SLC8A1*, *SLC9A1*, *ANK2*, *OSGIN1*, *SYNE2*, *TP63*, *PTPRT*, *TNFRSF1B*, *ICA1*, *RCVRN*, *CD9*, *ARHGAP44*, *SEMA3C*, *CD36*, *SCARB1*, *C8orf44-SGK3*, *CTDSPL2*, *RFX3*, *LYN*, *KLHL24*, *MARK4*, *GTSE1*, *POFUT2*, *AHI1*, *FER*, *FGF1*, *DACH1*, *LAMA1*, *NR1H4*, *NISCH*, *BCAS3*, *DAPK1*, *FGF14*, *GPC5*, *SEMA3G*, *FGR*, *IGF1R*, *NEDD4L*, *CLASP1*, *BBC3*, *BMP2K*, *RIPK1*, *GRB7*, *SHISA6*, *KCNH5*, *MAPT*, *GRIK5*, *PACSIN2*, *PDPN*, *PPARA*, *PPARD*, *PPP1R13B*, *NOS1*, *ROBO1*, *FLT1*, *ROCK1*<br>*GRM7*, *RGS9*, *DNM1L*, *KHDRBS1*, *RAB11FIP5*, *CD300A*, *CACNG8*, *AKAP6*, *CACNG6*, *PTPRU*, *ONECUT2*, *ATP1A1*, *FGF18*, *NPY2R*, *GNB5*, *DLG2*, *SHANK1*, *MAP3K3*, *LGR6*, *PPP3CB*, *PPP3CC*, *PPP3R1*, *KCNH7*, *CASS4*, *MADCAM1*, *NRG3*, *NKAIN2*, *NTRK3*, *LDLRAP1*, *CCDC88A*, *ROR2*, *CHRM3*, *VPS4B*, *TMEM59*, *NUMA1*, *BAG4*, *MGAT5*, *DOCK1*, *ULK4*, *NRG2*, *RAB3D*, *PRKAR1B*, *DPP4*, *PRKCB*, *PRKCG*, *DPYSL3*, *STK10*, *IL34*, *FTO*, *NTNG1*, *TMEM30A*, *DOCK5*, *DSCAM*, *MCU*, *UTRN*, *NLGN1*, *OPRK1*, *STXBP5*, *NTNG2*, *DYSF*, *MEMO1*, *BCL2*, *STARD13*, *AKAP12*, *PCLO*, *MCTP2*, *VEGFC*, *HCN2*, *RYR1*, *DVL1*, *SYT1*, *MMP28*, *MTOR*, *RUFY3*, *AP3D1*, *GLIS2*, *P2RX1* |

| | | | |
|---|---|---|---|
| GO:0010975 | Regulation of neuron projection development | $8.60 \times 10^{-6}$ | *SEMA3C*, *CDKL3*, *SFRP1*, *PTPRO*, *ABL1*, *FSTL4*, *MYLIP*, *GRIP1*, *NRCAM*, *LYN*, *PARK2*, *CAMK2G*, *ITM2C*, *NTRK3*, *ZMYND8*, *UBE2V2*, *ROR1*, *SPAG9*, *ROR2*, *CCDC88A*, *SEMA3G*, *ACAP3*, *CDH4*, *CUX2*, *ULK4*, *ALK*, *NEDD4L*, *PLXNA4*, *DPYSL3*, *LRIG2*, *SDK1*, *EEF2K*, *MAGI2*, *NTNG1*, *CTNNA2*, *TMEM30A*, *MAPT*, *DSCAM*, *NLGN1*, *PLK5*, *NTNG2*, *SEMA4A*, *TNIK*, *CREB3L2*, *TRPV2*, *ROBO1*, *HECW2*, *SPOCK1*, *SETX*, *RAB17*, *SS18L1*, *DVL1*, *SYT1*, *MTOR*, *DNM1L*, *RUFY3*, *DISC1*, *ARHGAP44*, *FBXO38* |
| GO:0051271 | Negative regulation of cellular component movement | $1.64 \times 10^{-5}$ | *SEMA3C*, *LIMCH1*, *PTPRK*, *ITGB1BP1*, *CD300A*, *FRMD5*, *STARD13*, *PPARD*, *ZMYND8*, *DACH1*, *CLASP1*, *NISCH*, *PTPRT*, *MMP28*, *PODN*, *SEMA3G* |
| GO:0040013 | Negative regulation of locomotion | $2.36 \times 10^{-5}$ | *SEMA3C*, *LIMCH1*, *PTPRK*, *ITGB1BP1*, *CD300A*, *FRMD5*, *STARD13*, *PPARD*, *ZMYND8*, *DACH1*, *CLASP1*, *NISCH*, *PTPRT*, *MMP28*, *PODN*, *SEMA3G* |
| GO:0040017 | Positive regulation of locomotion | $2.59 \times 10^{-5}$ | *SEMA3C*, *TACR1*, *GCNT2*, *ONECUT2*, *SCARB1*, *ABL1*, *FGF18*, *MAP3K3*, *ATP8A1*, *LGR6*, *LYN*, *SPOCK2*, *GTSE1*, *RHOJ*, *CASS4*, *MADCAM1*, *FER*, *NTRK3*, *LAMB1*, *FGF1*, *SPAG9*, *ROR2*, *LAMC2*, *VPS4B*, *BAG4*, *MGAT5*, *BCAS3*, *MEGF8*, *DOCK1*, *SEMA3G*, *FGR*, *CCL5*, *MYLK*, *IGF1R*, *ALOX12*, *CLASP1*, *IL34*, *GRB7*, *C5AR1*, *DOCK5*, *SMURF2*, *DSCAM*, *MCU*, *PIK3CD*, *PIK3R1*, *KIT*, *ITGB1BP1*, *SEMA4A*, *PDPN*, *BCL2*, *AKAP12*, *RIN2*, *SLC8A1*, *FLT1*, *VEGFC*, *SYNE2*, *MTOR*, *DNM1L*, *RUFY3* |

| | | | |
|---|---|---|---|
| GO:0022603 | Regulation of anatomical structure morphogenesis | $3.11 \times 10^{-5}$ | *SEMA3C*, *MMP20*, *JAK1*, *CREB3L1*, *CD36*, *ABL1*, *PARK2*, *RHOJ*, *AHI1*, *PSMA7*, *PSMB2*, *WNT9B*, *TBX2*, *FGF1*, *SPAG9*, *MYH14*, *MEGF8*, *PSMD8*, *SEMA3G*, *CDH4*, *CUX2*, *FGR*, *NEDD4L*, *AGO1*, *PLXNA4*, *CLASP1*, *WTIP*, *HIPK2*, *C5AR1*, *SH3D19*, *MAPT*, *ADAM12*, *KIT*, *SEMA4A*, *TNIK*, *PDPN*, *RIN2*, *ARHGEF18*, *FGD4*, *FLT1*, *ROBO1*, *HECW2*, *ROCK1*, *GATA6*, *ENG*, *SS18L1*, *HMGA2*, *DNM1L*, *DISC1*, *PPP1R16B*, *CDKL3*, *SFRP1*, *PTPRM*, *PTPRO*, *FSTL4*, *FGF18*, *GRIP1*, *NRCAM*, *MAP3K3*, *CASS4*, *LARP4*, *CAPN2*, *EPB41L3*, *ROR1*, *ROR2*, *ZMYM4*, *ZMYM6*, *STAT2*, *DOCK1*, *ALOX12*, *PRKCB*, *EEF2K*, *MAGI2*, *NTNG1*, *PRKDC*, *DOCK5*, *SMURF2*, *DSCAM*, *MCU*, *PIK3CD*, *NLGN1*, *LIMS1*, *ITGB1BP1*, *NTNG2*, *PLEKHA4*, *TRPV2*, *STARD13*, *BCL2*, *ITGA7*, *PKM*, *VEGFC*, *DVL1*, *TNFRSF1B*, *SYT1*, *MTOR*, *CELSR1*, *RUFY3*, *ARHGAP44* |
| GO:0051272 | Positive regulation of cellular component movement | $3.47 \times 10^{-5}$ | *SEMA3C*, *TACR1*, *GCNT2*, *ONECUT2*, *SCARB1*, *ABL1*, *FGF18*, *MAP3K3*, *ATP8A1*, *LGR6*, *LYN*, *SPOCK2*, *GTSE1*, *RHOJ*, *CASS4*, *MADCAM1*, *FER*, *NTRK3*, *LAMB1*, *FGF1*, *SPAG9*, *ROR2*, *LAMC2*, *BAG4*, *MGAT5*, *BCAS3*, *MEGF8*, *DOCK1*, *SEMA3G*, *FGR*, *CCL5*, *MYLK*, *IGF1R*, *ALOX12*, *CLASP1*, *IL34*, *GRB7*, *C5AR1*, *DOCK5*, *SMURF2*, *DSCAM*, *MCU*, *PIK3CD*, *PIK3R1*, *KIT*, *ITGB1BP1*, *SEMA4A*, *PDPN*, *BCL2*, *AKAP12*, *RIN2*, *SLC8A1*, *FLT1*, *VEGFC*, *SYNE2*, *MTOR*, *DNM1L*, *RUFY3* |

Table S4: Gene Ontology (GO) Cellular Component for CEU genes ranked based on *TrIdent* prediction output.

| GO Code | Description | $P$-value | Genes |
|---|---|---|---|
| GO:0005886 | Plasma membrane | $1.16 \times 10^{-6}$ | *TACR1, P2RY6, ITPR1, BAIAP2L2, DSG4, ABCA1, SLC4A10, ABCC1, ABCA3, PCSK6, APCDD1, GNAT3, GPHN, HLA-DQA1, ASIC2, RHOJ, SGCZ, PAWR, KCNC3, PKD2L1, COL6A3, KCND3, HGSNAT, CNKSR1, GAS2L2, BRAF, XKR5, EFNA2, ANO10, BSG, SLC24A3, RAMP1, KCNJ6, PLEKHG5, RAB44, SCNN1B, TENM2, PLA2G6, SPG11, SRL, GPRC5C, CHIC2, COL23A1, GABRG3, KCNQ1, CLDN6, MSR1, PLXNA4, ERBB2IP, NOX5, PREX2, ERAP2, C5AR1, ADCY8, RASGRP4, KIT, CPE, DNAJC1, ADD3, SEMA4A, CUL1, MTUS1, PDE6B, GPR158, CEP112, CR1, ENG, RAB17, CACNA1A, SLC39A11, PTPRK, EPB41L1, PTPRM, SFRP1, PTPRN, ADRBK1, PTPRN2, DMTN, HIF3A, PTPRO, OR6J1, CACNB4, CACNG2, DGKI, MYLIP, MUC2, EPOR, MUC5AC, NFASC, ITGA8, TTYH3, ATP8A1, CLASP2, GAS7, MX1, PEMT, MX2, CSF2RB, FCAMR, CAPN2, EPB41L3, LILRB3, HRH2, TGM2, RAB31, NECAB2, RAB5C, LRRFIP1, PMEL, MELK, LCK, PHKB, FAIM3, LYPLA1, LCT, MAGI1, GAB2, C2CD5, MYLK, ALK, DLGAP1, ALOX12, GCA, ARHGAP10, DPP10, SLC1A7, RAPGEF4, SLC2A2, LRIG2, MAGI2, PIGR, PARD3, CTNNA2, PIK3C2B, PPP1R12A, DNAJB4, SMURF2, SLC2A10, CTNND2, LIM2, PIK3CD, PIK3R1, LIMS1, SLC6A6, SPRED3, RGS20, ITGB1BP1, PLEKHA4, AIG1, TRPV2, PLA2G4C, PIP4K2A, KIAA1549, SLC7A2, GNA12, HTR4, SLC8A1, PKD1, SLC9A1, GPR55, NCAM2, SLC9A3, GNAL, ANK2, SNX9, PTPRT, TNFRSF1B, ABCC11, PI4K2A, TPTE2, ICAM2, CD9, GNG4, GNG7, PLCD1, KIAA0922, SCHIP1, SLC20A2, SLA2, APBA2, GPR160, CD36, SCARB1, KAZN, SLC13A3, SLITRK6, ENTPD3, GLP2R, APLP2, SLC4A4, FAM155A, LYN, KCMF1, RGS3, OR52R1* |

| | | | |
|---|---|---|---|
| | | | *SLC30A8, UTS2R, MGRN1, NRXN3, LRBA, DGKB, NISCH, TRHR, DAPK1, GPC5, SLC4A5, CATSPERD, CDH4, AMICA1, FGR, CDH7, FHIT, IGF1R, CDH12, AKAP10, NEDD4L, PAQR5, PMEPA1, NOX3, CDH18, SOS1, ARHGDIG, CLMP, RIPK1, LRTOMT, NID1, GRB7, SH3D19, KCNH5, MAPT, DEF6, FNBP1L, ADAM12, GRIK1, GRIK4, GRIK5, PACSIN2, PDPN, SIGLEC10, STAP2, PPP1R13B, ARHGEF18, TENC1, GRM1, NOS1, ROBO1, FLT1, ITM2B, ROCK1, GRM7, RGS9, ITFG1, FLVCR2, PPP1R16B, TULP3, CD300A, CACNG8, CACNG6, PTPRU, RHOT2, ATP1A1, ATP1A3, NPY2R, DLG2, GRIP1, NRCAM, AHNAK2, SHANK1, UBE2B, LGR6, PPP3CB, WWOX, PPP3R1, KCNH7, CASS4, ENPP6, MADCAM1, ITM2C, NKAIN2, SLC44A1, KIAA0319L, ITGA11, TMEM106A, NTRK3, LDLRAP1, ROR1, GNG13, CCDC88A, ROR2, TBC1D10C, ABCA13, CHRM3, NDC1, ASAP2, TMEM59, BAG4, STAT2, CHRNB3, MGST1, NRG2, RAB3D, PRKAR1B, INPP4A, INPP5A, DPP4, DLGAP4, PRKCB, PRKCG, STK10, FTO, SDK1, NTNG1, COL25A1, TMEM30A, DOCK5, DSCAM, CLRN1, UTRN, NLGN1, OPRK1, STXBP5, NTNG2, VAV1, DYSF, DTNA, EXOC2, NRSN2, AKAP12, ITGA7, PIEZO2, SLCO3A1, SVIL, LHFPL4, UNC80, HCN2, RYR1, SCUBE3, SYT1, CELSR1, P2RX1* |

| | | | |
|---|---|---|---|
| GO:0097458 | Neuron part | $8.2 \times 10^{-6}$ | *EYS*, *ITPR1*, *APBA2*, *SLC4A10*, *GNAT3*, *ABCA4*, *ABL1*, *SMARCA2*, *GPHN*, *LYN*, *KLHL24*, *MARK4*, *PARK2*, *ASIC2*, *KCNC3*, *DST*, *KCND3*, *ZMYND8*, *MYH14*, *SCN4A*, *EFNA2*, *BSG*, *PLEKHG5*, *MAP7*, *TENM2*, *PSD2*, *SPG11*, *BRINP1*, *IGF1R*, *GABRG3*, *URI1*, *MAK*, *SOS1*, *WDR7*, *ESPN*, *SHISA6*, *STX12*, *MAPT*, *ADCY8*, *PCDH15*, *GRIK4*, *GRIK5*, *ADD3*, *TNIK*, *SPAST*, *PDE6B*, *SH2D3C*, *GRM1*, *NMNAT2*, *NOS1*, *ROBO1*, *SPOCK1*, *RPTOR*, *GRM7*, *SETX*, *RAB17*, *SS18L1*, *RGS9*, *SYBU*, *CNTNAP2*, *TPGS1*, *CACNA1A*, *DNM1L*, *DISC1*, *SFPQ*, *PTPRN*, *ERC1*, *PTPRN2*, *PTPRO*, *CACNG8*, *DICER1*, *PEX6*, *CACNG2*, *PVALB*, *ATP1A1*, *ATP1A3*, *DGKI*, *GNB5*, *DLG2*, *MYO3B*, *GRIP1*, *NRCAM*, *ITGA8*, *NFASC*, *SHANK1*, *PPP3CC*, *MX1*, *SYNPO*, *MX2*, *AHCY*, *CAMK2G*, *CASC3*, *TMEM163*, *LSM1*, *RSPH9*, *CAPN2*, *EPB41L3*, *NTRK3*, *LDLRAP1*, *GNG13*, *ROR1*, *ROR2*, *CHRM3*, *NECAB2*, *NUMA1*, *CHRNB3*, *ACAP3*, *RAB3D*, *INPP4A*, *INPP5A*, *DLGAP1*, *PRKCB*, *AURKA*, *PRKCG*, *REEP6*, *DPYSL3*, *ELFN1*, *SLC1A7*, *LRIG2*, *NTNG1*, *EEF2K*, *MAGI2*, *PARD3*, *PKHD1L1*, *CTNNA2*, *CTNND2*, *DSCAM*, *CLRN1*, *UTRN*, *NLGN1*, *TXN2*, *OPRK1*, *SLC6A6*, *STXBP5*, *NTNG2*, *TRPV2*, *NRSN2*, *AKAP12*, *PCLO*, *KIAA1549*, *HTR4*, *CLSTN1*, *SLC8A1*, *LHFPL4*, *NCAM2*, *ANK2*, *UNC80*, *MCTP2*, *SNX9*, *TP63*, *PTPRT*, *DVL1*, *TNFRSF1B*, *SYT1*, *PI4K2A*, *ICA1*, *MTOR*, *RUFY3*, *AP3D1*, *RCVRN*, *LMTK3*, *P2RX1*, *IFT52*, *ARHGAP44*, *TOPORS* |
| GO:0043256 | laminin complex | $1.35 \times 10^{-5}$ | *LAMC2*, *NTNG1*, *LAMA1*, *NTNG2*, *LAMB4*, *LAMA3*, *LAMB1* |

| | | | |
|---|---|---|---|
| GO:0005856 | Cytoskeleton | $4.3 \times 10^{-5}$ | *JAK1*, *KAZN*, *ABL1*, *SMARCA2*, *CCDC110*, *GPHN*, *GTSE1*, *MARK4*, *SGCZ*, *ARPC1A*, *PAWR*, *KCNC3*, *DST*, *FER*, *IQCA1*, *NISCH*, *DAPK1*, *BCAS3*, *MAP7*, *FHOD3*, *TUBA1C*, *FGR*, *CLASP1*, *APC2*, *HAUS3*, *CCDC6*, *MAPT*, *DEF6*, *EFCAB2*, *FNBP1L*, *ADCY8*, *DENND2A*, *ADD3*, *TNIK*, *PACSIN2*, *SPAST*, *MTUS1*, *ARHGEF18*, *NOS1*, *CR1*, *FGD4*, *CENPV*, *FLT1*, *ROCK1*, *TCTN2*, *RSPH3*, *TUBA4A*, *DISC1*, *SPTAN1*, *ODF3L2*, *EPB41L1*, *ERC1*, *DMTN*, *SLAIN1*, *ARHGAP26*, *ONECUT2*, *MYLIP*, *CORO2B*, *ARPC1B*, *GTF2F2*, *MSRB1*, *EVC2*, *GAS7*, *MX1*, *MX2*, *SYNPO*, *CASS4*, *CAPN2*, *EPB41L3*, *SSH3*, *NDC1*, *DCDC2C*, *LRRFIP1*, *FRMD4B*, *JAKMIP2*, *MYLK*, *FRMD5*, *AURKA*, *KANK4*, *PPP1R42*, *SLC30A9*, *PARD3*, *CTNNA2*, *WASF2*, *PPP1R12A*, *SPAG6*, *CLRN1*, *UTRN*, *ITGB1BP1*, *DTNA*, *AKAP12*, *PCLO*, *NUAK1*, *SVIL*, *ANK2*, *NOL9*, *SYNE2* |

| | | | |
|---|---|---|---|
| GO:0120038 | Plasma membrane bounded cell projection part | $4.41 \times 10^{-5}$ | *TACR1*, *EYS*, *APBA2*, *SLC4A10*, *CD36*, *SCARB1*, *GNAT3*, *ABCA4*, *ABL1*, *GPHN*, *MARK4*, *ASIC2*, *AHI1*, *KCNC3*, *DST*, *PKD2L1*, *KCND3*, *GAS2L2*, *ZMYND8*, *MYH14*, *BSG*, *TENM2*, *PSD2*, *SPG11*, *ARMC9*, *BRINP1*, *CATSPERD*, *FGR*, *GABRG3*, *URI1*, *MAK*, *APC2*, *ESPN*, *SHISA6*, *MAPT*, *C2CD3*, *ADCY8*, *PCDH15*, *GRIK5*, *PACSIN2*, *PSD3*, *PDPN*, *SPAST*, *ARHGEF4*, *PDE6B*, *SH2D3C*, *GRM1*, *NOS1*, *SPOCK1*, *RPTOR*, *GRM7*, *SETX*, *RAB17*, *ARL8B*, *TCTN2*, *SYBU*, *CNTNAP2*, *TPGS1*, *DISC1*, *SFPQ*, *TULP3*, *PTPRN*, *ERC1*, *PTPRN2*, *DMTN*, *PTPRO*, *DICER1*, *PEX6*, *DGKI*, *DLG2*, *MYO3B*, *GRIP1*, *NRCAM*, *ITGA8*, *NFASC*, *SHANK1*, *EVC2*, *CLASP2*, *MX1*, *PEMT*, *SYNPO*, *MX2*, *CASC3*, *LSM1*, *RSPH9*, *CAPN2*, *EPB41L3*, *HRH2*, *GNG13*, *ROR1*, *CCDC88A*, *ROR2*, *TBC1D10C*, *CHRM3*, *NECAB2*, *NUMA1*, *DNAH7*, *ACAP3*, *PRKAR1B*, *C2CD5*, *CCDC178*, *INPP5A*, *DPP4*, *PRKCB*, *AURKA*, *PRKCG*, *DPYSL3*, *ELFN1*, *MXRA8*, *TTC26*, *TTLL1*, *LRIG2*, *EEF2K*, *MAGI2*, *PARD3*, *PKHD1L1*, *SPAG6*, *CTNND2*, *DSCAM*, *UTRN*, *NLGN1*, *TXN2*, *SLC6A6*, *TRPV2*, *DNAH10*, *KIAA1549*, *GNA12*, *HTR4*, *CLSTN1*, *SLC8A1*, *PKD1*, *LHFPL4*, *SLC9A3*, *TP63*, *SYNE2*, *DVL1*, *TNFRSF1B*, *SYT1*, *PI4K2A*, *DNAH17*, *MTOR*, *AP3D1*, *RUFY3*, *RCVRN*, *LMTK3*, *IFT52*, *ARHGAP44*, *TOPORS* |

| | | | |
|---|---|---|---|
| GO:0044463 | Cell projection part | $4.41 \times 10^{-5}$ | *TACR1, EYS, APBA2, SLC4A10, CD36, SCARB1, GNAT3, ABCA4, ABL1, GPHN, MARK4, ASIC2, AHI1, KCNC3, DST, PKD2L1, KCND3, GAS2L2, ZMYND8, MYH14, BSG, TENM2, PSD2, SPG11, ARMC9, BRINP1, CATSPERD, FGR, GABRG3, URI1, MAK, APC2, ESPN, SHISA6, MAPT, C2CD3, ADCY8, PCDH15, GRIK5, PACSIN2, PSD3, PDPN, SPAST, ARHGEF4, PDE6B, SH2D3C, GRM1, NOS1, SPOCK1, RPTOR, GRM7, SETX, RAB17, ARL8B, TCTN2, SYBU, CNTNAP2, TPGS1, DISC1, SFPQ, TULP3, PTPRN, ERC1, PTPRN2, DMTN, PTPRO, DICER1, PEX6, DGKI, DLG2, MYO3B, GRIP1, NRCAM, ITGA8, NFASC, SHANK1, EVC2, CLASP2, MX1, PEMT, SYNPO, MX2, CASC3, LSM1, RSPH9, CAPN2, EPB41L3, HRH2, GNG13, ROR1, CCDC88A, ROR2, TBC1D10C, CHRM3, NECAB2, NUMA1, DNAH7, ACAP3, PRKAR1B, C2CD5, CCDC178, INPP5A, DPP4, PRKCB, AURKA, PRKCG, DPYSL3, ELFN1, MXRA8, TTC26, TTLL1, LRIG2, EEF2K, MAGI2, PARD3, PKHD1L1, SPAG6, CTNND2, DSCAM, UTRN, NLGN1, TXN2, SLC6A6, TRPV2, DNAH10, KIAA1549, GNA12, HTR4, CLSTN1, SLC8A1, PKD1, LHFPL4, SLC9A3, TP63, SYNE2, DVL1, TNFRSF1B, SYT1, PI4K2A, DNAH17, MTOR, AP3D1, RUFY3, RCVRN, LMTK3, IFT52, ARHGAP44, TOPORS* |
| GO:0098984 | Neuron to neuron synapse | $1.11 \times 10^{-4}$ | *RAPGEF4, NLGN1, ATP1A3, GRIK4, PRKAR1B, GRIK5, STXBP5, GRM7, SHISA6, SYT1, CACNG2, ADCY8* |
| GO:0099572 | postsynaptic specialization | $1.23 \times 10^{-4}$ | *DMTN, INPP4A, ITPR1, CACNG8, DLGAP1, DLGAP4, PRKCG, SOS1, ATP1A1, DGKI, MAGI2, EEF2K, SHISA6, DLG2, GRIP1, ITGA8, SHANK1, CTNND2, LYN, GPHN, ADCY8, NLGN1, MX1, MX2, SYNPO, ADD3, PSD3, KCND3, PCLO, SLC8A1, NOS1, SPOCK1, DVL1, DISC1, ARHGAP44* |
| GO:0015630 | Microtubule cytoskeleton | $1.59 \times 10^{-4}$ | *AURKA, KANK4, APC2, PPP1R42, HAUS3, GTF2F2, SPAG6, MAPT, CLRN1, GTSE1, MARK4, MX1, MX2, DST, FER, SPAST, MTUS1, NUAK1, SS18, CENPV, NISCH, MAP7, JAKMIP2, TUBA4A, TUBA1C, SPTAN1* |

| | | | |
|---|---|---|---|
| GO:0043005 | Neuron projection | $1.69 \times 10^{-4}$ | *APBA2*, *SLC4A10*, *ABL1*, *GPHN*, *KLHL24*, *MARK4*, *ASIC2*, *PARK2*, *KCNC3*, *KCND3*, *ZMYND8*, *SCN4A*, *BSG*, *PLEKHG5*, *MAP7*, *TENM2*, *PSD2*, *SPG11*, *BRINP1*, *IGF1R*, *GABRG3*, *URI1*, *ESPN*, *MAPT*, *ADCY8*, *PCDH15*, *GRIK5*, *SH2D3C*, *NMNAT2*, *GRM1*, *NOS1*, *ROBO1*, *RPTOR*, *SETX*, *GRM7*, *RAB17*, *CNTNAP2*, *TPGS1*, *PTPRO*, *DICER1*, *PEX6*, *CACNG2*, *PVALB*, *DGKI*, *ATP1A3*, *DLG2*, *GRIP1*, *NRCAM*, *NFASC*, *SHANK1*, *MX1*, *MX2*, *SYNPO*, *AHCY*, *CAMK2G*, *CASC3*, *LSM1*, *RSPH9*, *CAPN2*, *NTRK3*, *LDLRAP1*, *GNG13*, *ROR2*, *CHRM3*, *NECAB2*, *NUMA1*, *CHRNB3*, *INPP5A*, *PRKCG*, *ELFN1*, *EEF2K*, *MAGI2*, *CTNNA2*, *CTNND2*, *DSCAM*, *CLRN1*, *TXN2*, *NLGN1*, *OPRK1*, *SLC6A6*, *NTNG2*, *NRSN2*, *PCLO*, *HTR4*, *SLC8A1*, *LHFPL4*, *NCAM2*, *ANK2*, *UNC80*, *TP63*, *DVL1*, *SYT1*, *PI4K2A*, *MTOR*, *RUFY3*, *RCVRN*, *LMTK3*, *P2RX1*, *ARHGAP44* |
| GO:0005955 | Calcineurin complex | $1.82 \times 10^{-4}$ | *PPP3CC*, *PPP3R1*, *ITPR1*, *PPP3CB* |

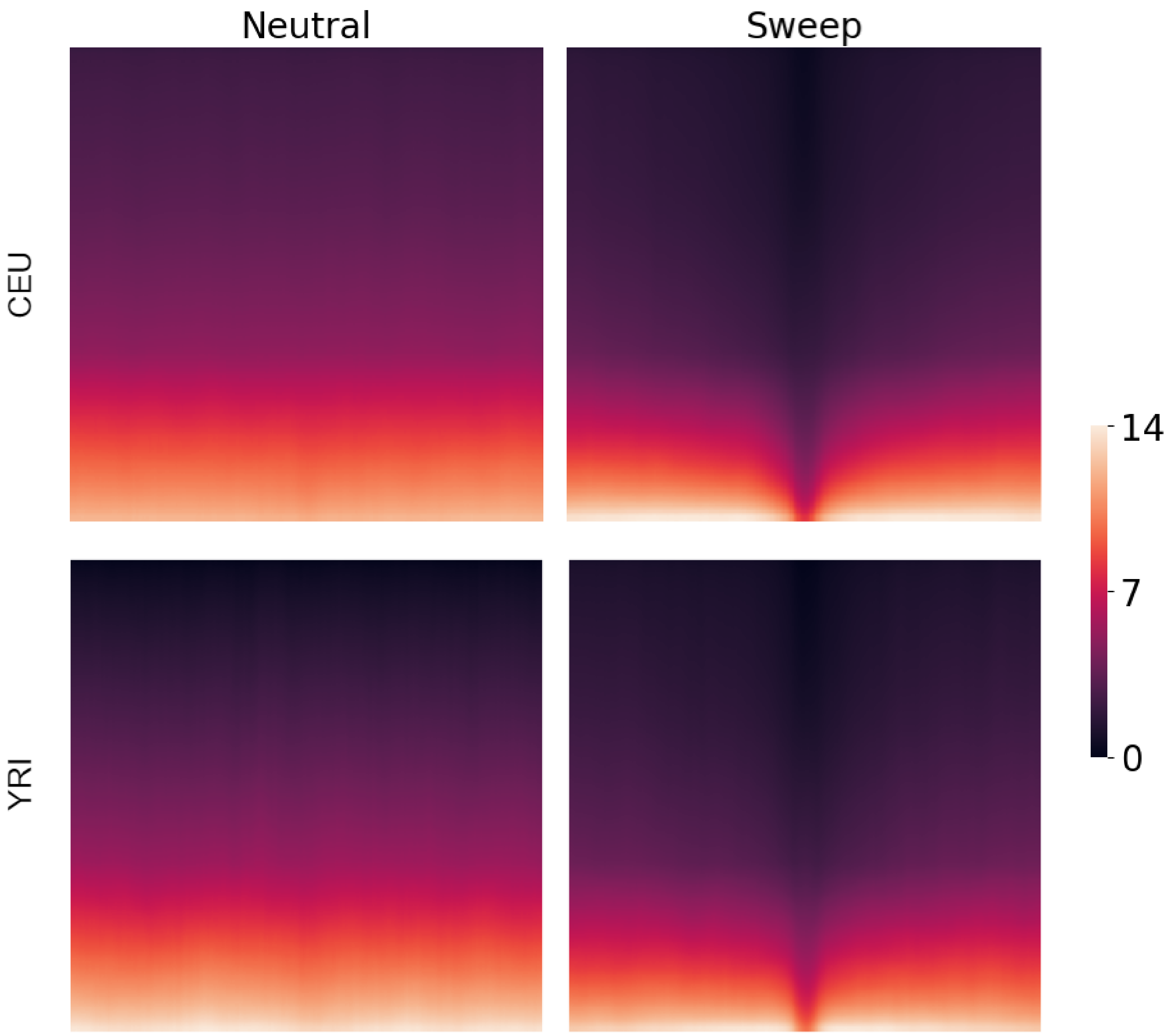


Figure S1: Heatmaps depicting input images of size 224 × 224, averaged across the 1,000 training replicates for either the neutral or sweep class simulated under either the European (CEU) or Sub-Saharan African (YRI) human demographic history [Tennessen et al., 2012]. Input images are processed as in the *Image generation* subsection of the *Methods*. Rows of the images represent haplotypes, whereas columns represent genomic window of 25 contiguous SNPs within a haplotype, with an equal number of windows flanking the center of a simulated genomic region. The colorbar indicates the number of minor alleles within the haplotype window, with darker colors representing a high number of major alleles and brighter colors a high number of minor alleles. Figures S23 and S24 provide visual representations of the haplotype diversity surrounding select sweep and neutral replicates from the CEU and YRI datasets, respectively.

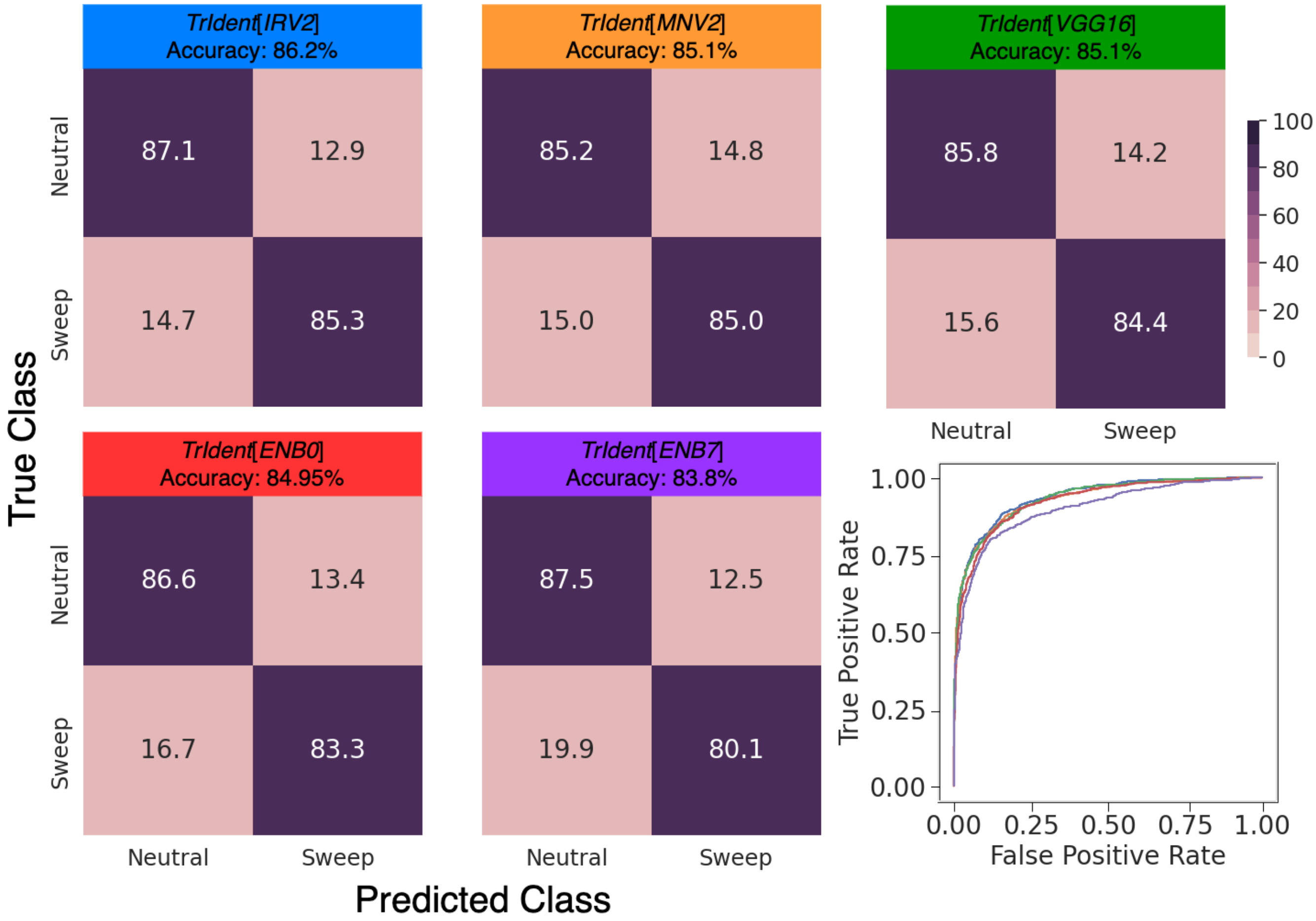


Figure S2: Classification rates and accuracies as depicted by confusion matrices and powers (true positive rates) to detect sweeps as depicted by receiver operating characteristic curves to differentiate sweeps from neutrality on the CEU dataset for *TrIdent*[*IRV2*], *TrIdent*[*MNV2*], *TrIdent*[*VGG16*], *TrIdent*[*ENB0*], and *TrIdent*[*ENB7*]. The CEU dataset is based on the recent strong bottleneck demographic history of central European humans (CEU population of the 1000 Genomes Project). This history includes a selective sweep that was completed within the last 2000 generations before sampling.

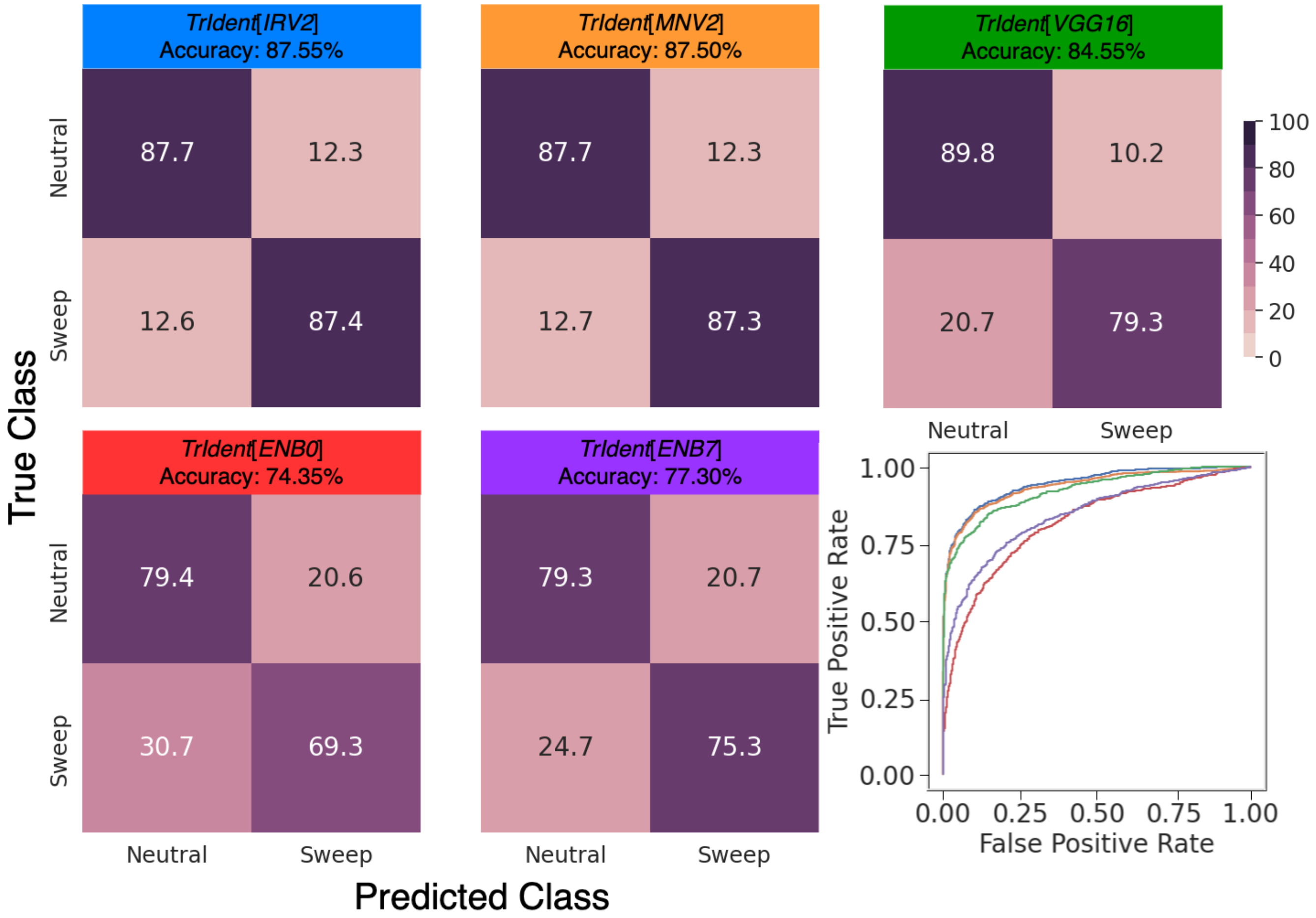


Figure S3: Classification rates and accuracies as depicted by confusion matrices and powers (true positive rates) to detect sweeps as depicted by receiver operating characteristic curves to differentiate sweeps from neutrality on the YRI dataset for *TrIdent*[*IRV2*], *TrIdent*[*MNV2*], *TrIdent*[*VGG16*], *TrIdent*[*ENB0*] and *TrIdent*[*ENB7*]. The YRI dataset is based on the demographic history of Sub-Saharan African humans experiencing a recent population expansion (YRI population of the 1000 Genomes Project). This history includes a selective sweep that was completed within the last 2000 generations before sampling.

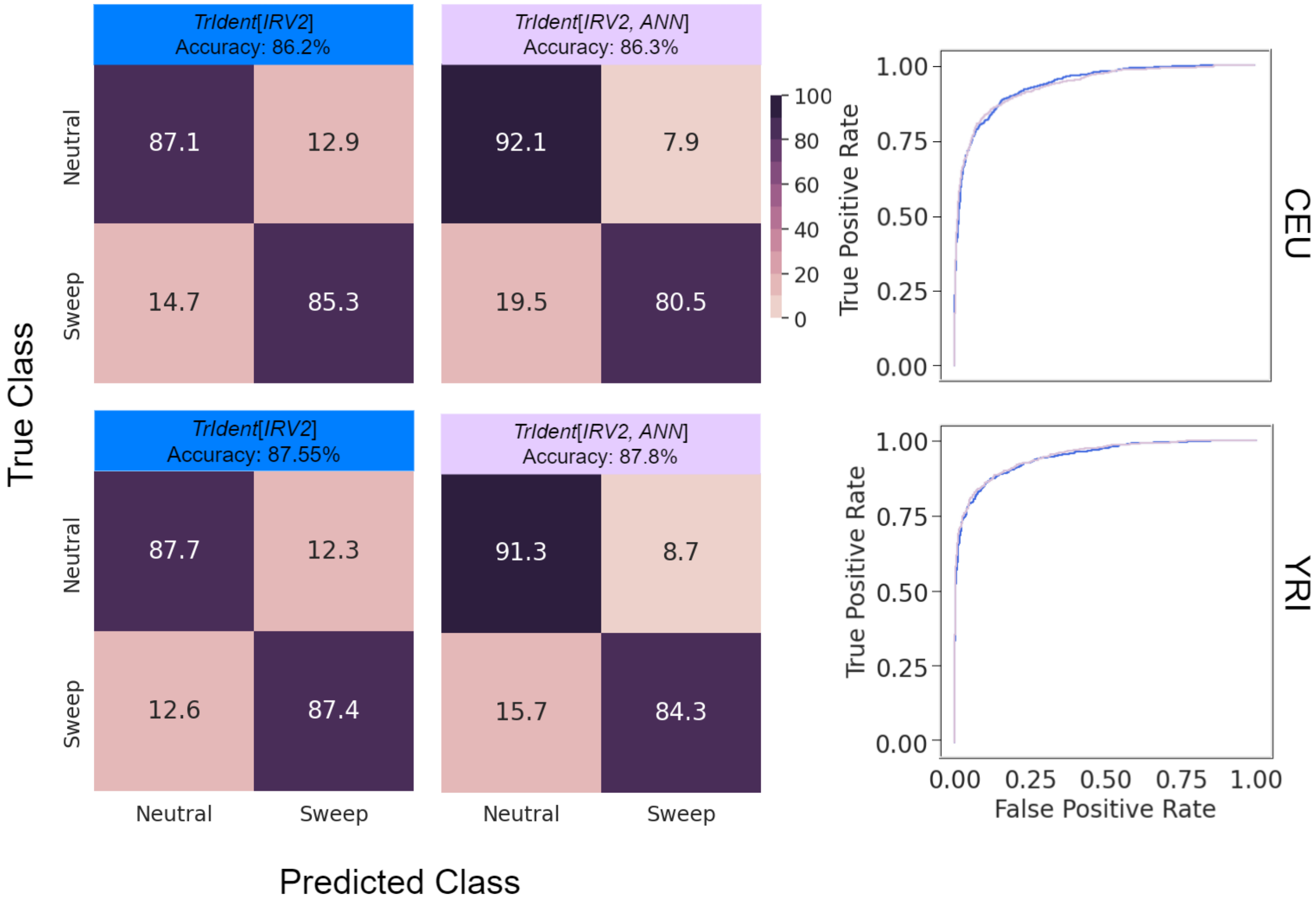


Figure S4: Classification rates and accuracies as depicted by confusion matrices, powers (true positive rates) to detect sweeps as depicted by receiver operating characteristic curves to differentiate sweeps from neutrality on the CEU (top panel) or YRI (bottom panel) test datasets for *TrIdent*[*IRV2*] and *TrIdent*[*IRV2*, *ANN*]. The *TrIdent*[*IRV2*, *ANN*] architecture is similar to that of *TrIdent*[*IRV2*] with the difference being that the GAP layer outputs are fed into a nonlinear artificial neural network (ANN) classifier instead of a linear (logistic regression) classifier.

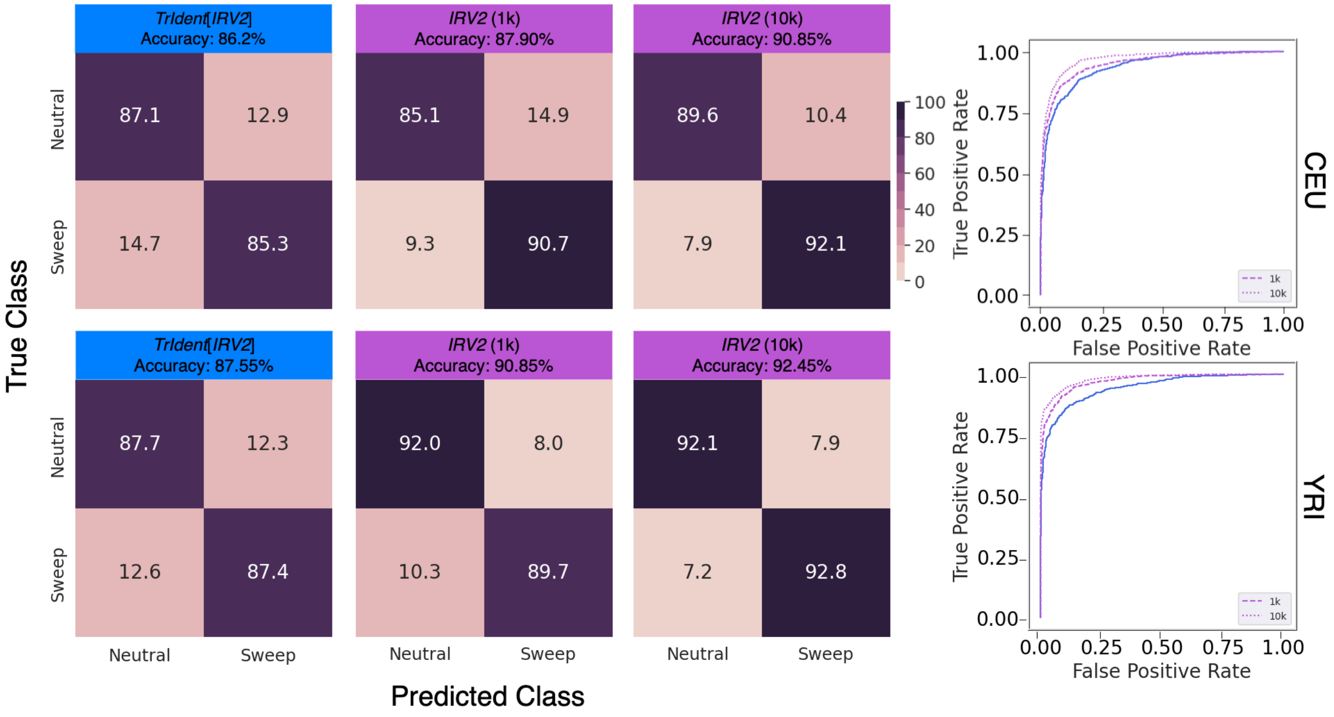


Figure S5: Classification rates and accuracies as depicted by confusion matrices and powers (true positive rates) to detect sweeps as depicted by receiver operating characteristic curves to differentiate sweeps from neutrality on the CEU (top panel) or YRI (bottom panel) test datasets for *TrIdent*[*IRV2*] and *IRV2*. The *IRV2* model incorporates the *InceptionResNetV2* architecture in addition to a GAP layer and a single node output layer. Contrary to *TrIdent*[*IRV2*] where we use *InceptionResNetV2* as a feature extraction block, we train all parameters within the complete *IRV2* model. For both CEU and YRI datasets, we present results for two *IRV2* models: one trained with 1,000 samples per class (*IRV2* (1k)) and another with 10,000 samples per class (*IRV2* (10k))

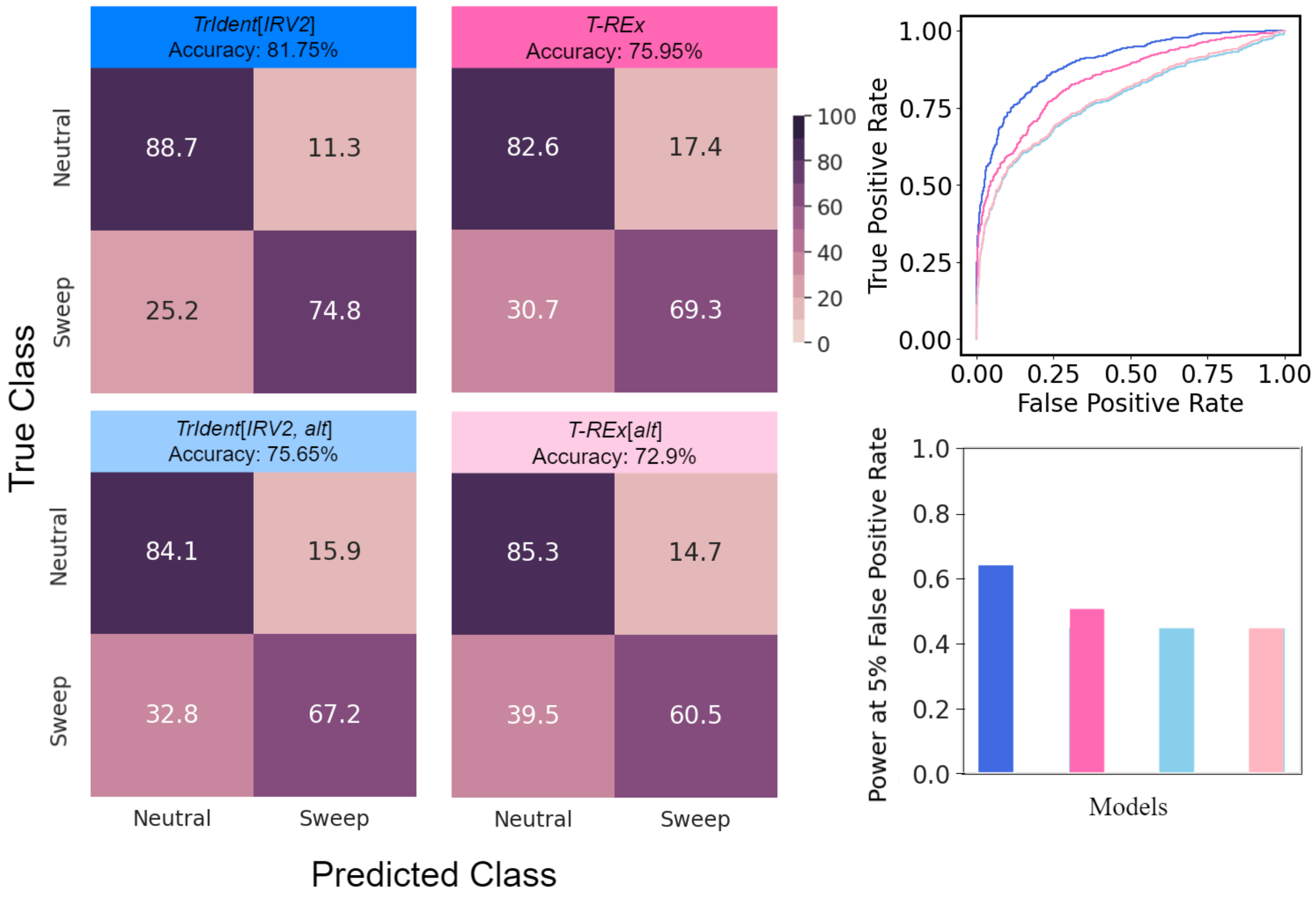


Figure S6: Classification rates and accuracies as depicted by confusion matrices, powers to detect sweeps as depicted by receiver operating characteristic curves to differentiate sweeps from neutrality, and powers at a 5% false positive rate to detect sweeps on the CEU test dataset containing missing genomic segments for *TrIdent*[*IRV2*], *TrIdent*[*IRV2*, *alt*], *T-REx*, and *T-REx*[*alt*]. The trained models are identical to those of Figure 3 and are trained to fit the training data without any missing information. However, the test data consists of sequences that have missing genomic blocks, which imitate the distribution of human genomic segments with low mean CRG score.

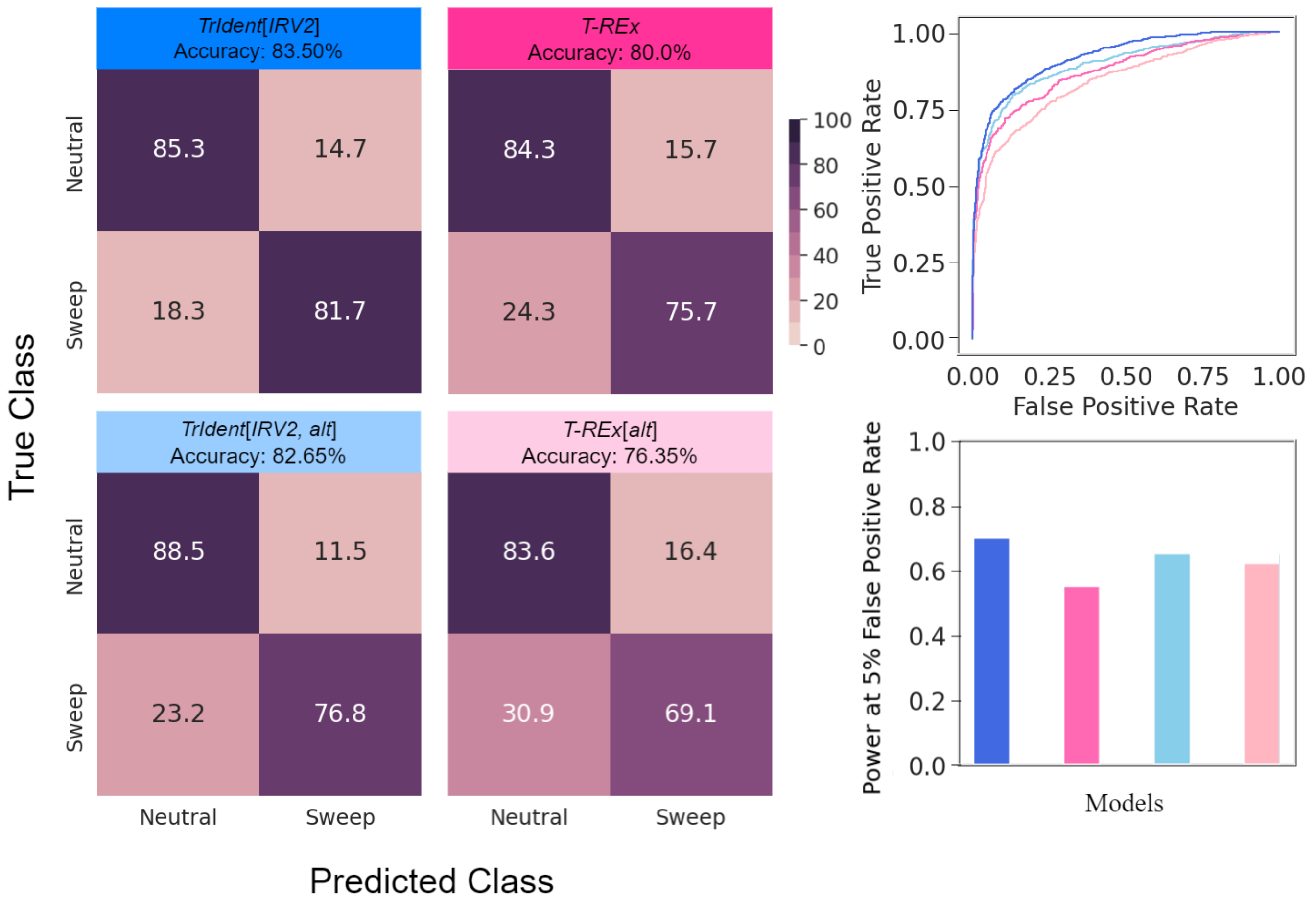


Figure S7: Classification rates and accuracies as depicted by confusion matrices, powers (true positive rates) to detect sweeps as depicted by receiver operating characteristic curves to differentiate sweeps from neutrality, and powers at a 5% false positive rate to detect sweeps on the YRI test dataset containing missing genomic segments for *TrIdent*[*IRV2*], *TrIdent*[*IRV2, alt*], *T-REx*, and *T-REx*[*alt*]. The trained models are identical to those of Figure 4 and are trained to fit the training data without any missing information. However, the test data consists of sequences that have missing genomic blocks, which imitate the distribution of human genomic segments with low mean CRG score.

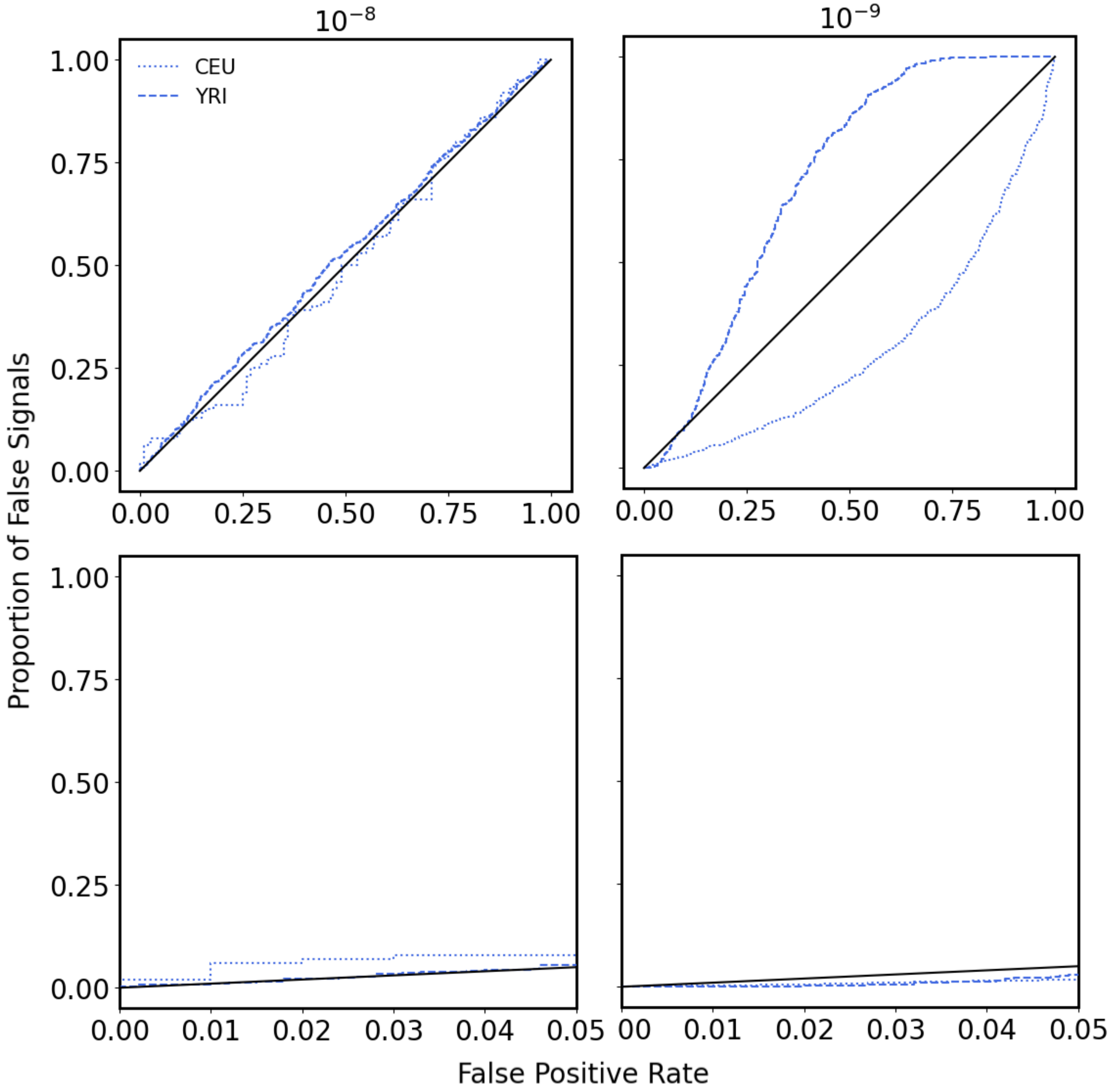


Figure S8: The probability of falsely detecting background selection as a sweep as a function of the false positive rate for the *TrIdent*[*IRV2*] model applied to the CEU and YRI datasets. Background selection test replicates had recombination rates drawn from an exponential distribution with mean of $10^{-8}$ (left) or $10^{-9}$ (right) per site per generation and truncated at three times the mean. Bottom panels are the same as the top panels, but with false positive rates restricted to less than 5%. Trained models are identical to those of Figure 3 for CEU and Figure 4 for YRI, with training sets that did not include background selection replicates. Test observations consisted either of neutral replicates to obtain the false positive rate distribution or of background selection replications to obtain the distribution of the proportion of false signals. The proportion of false signals is computed as the fraction of background selection replicates with a sweep probability greater than the sweep probability that generated a given false positive rate for neutral replicates.

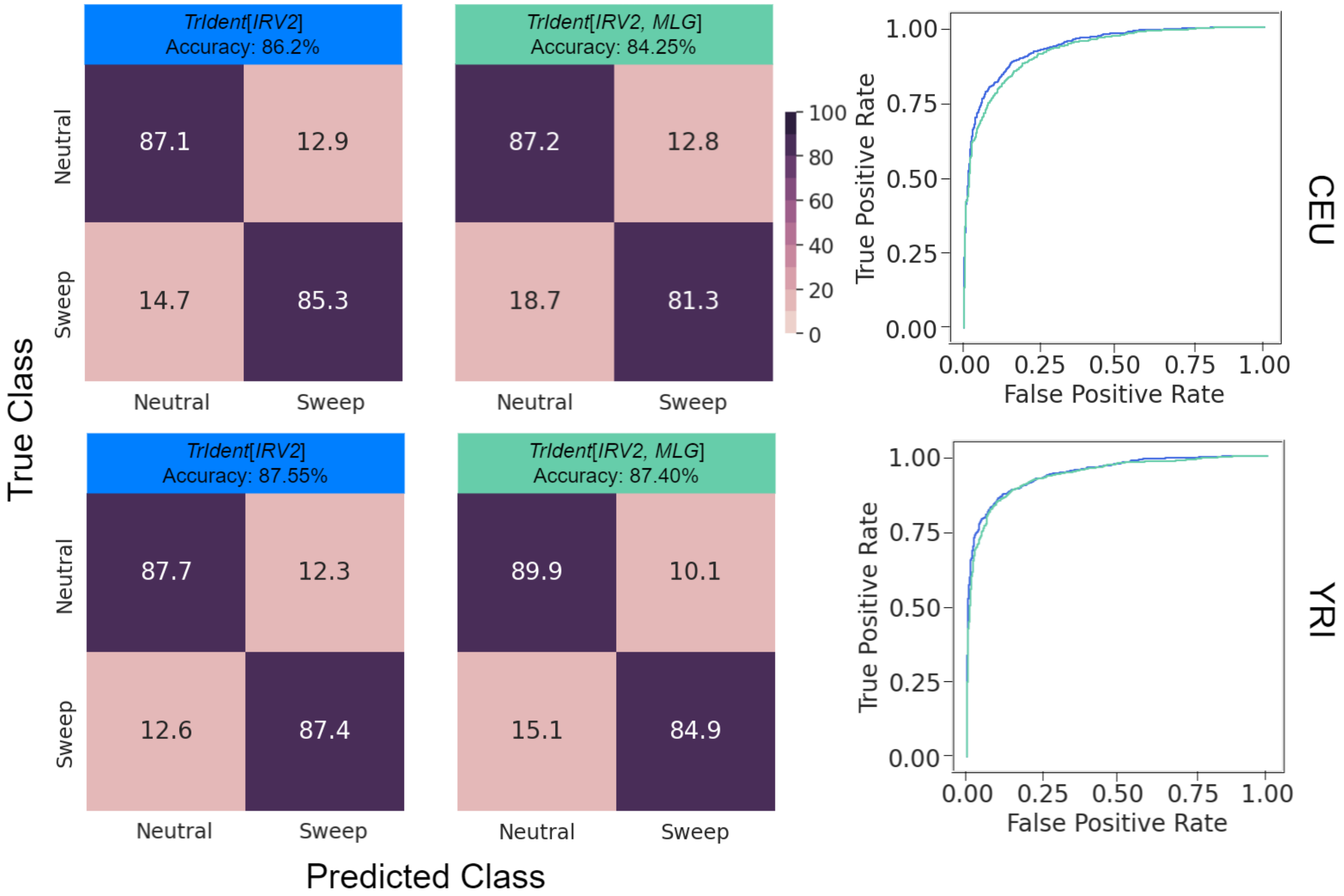


Figure S9: Classification rates and accuracies as depicted by confusion matrices, powers (true positive rates) to detect sweeps as depicted by receiver operating characteristic curves to differentiate sweeps from neutrality on the CEU (top panel) or YRI (bottom panel) test datasets for *TrIdent*[*IRV2*] and *TrIdent*[*IRV2*, *MLG*]. The *TrIdent*[*IRV2*, *MLG*] architecture is identical to that of *TrIdent*[*IRV2*] with the difference being that the input is based on unphased multilocus genotype data instead of phased haplotype data.

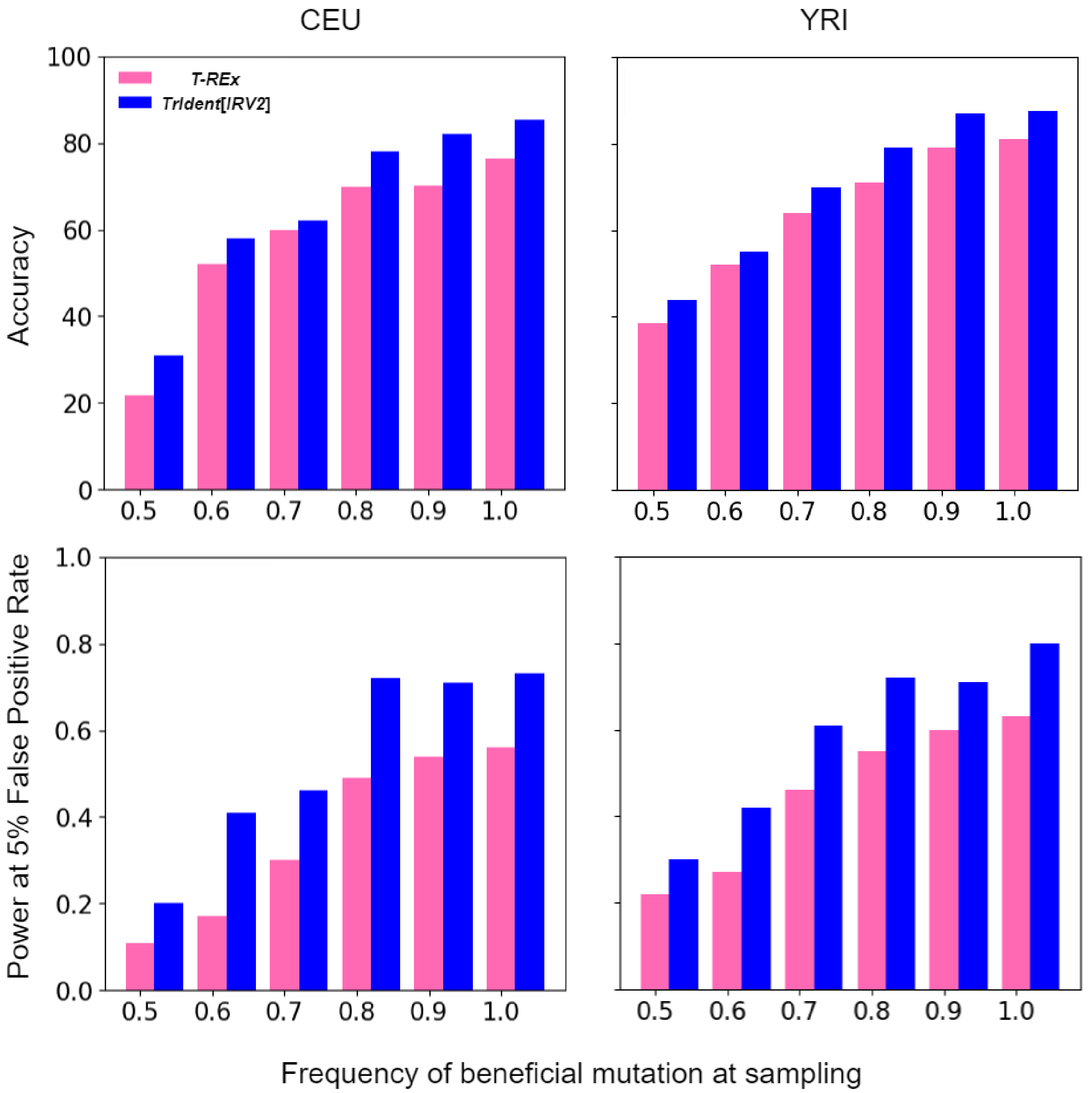


Figure S10: Accuracies, and powers at a 5% false positive rate to detect incomplete sweeps for the *TrIdent*[*IRV2*] and *T-REx* classifiers trained on replicates simulated with complete sweeps and applied to test replicates simulated under incomplete sweeps for which the beneficial mutation at the time of sampling is 0.5, 0.6, 0.7, 0.8, or 0.9, and compared against complete sweeps for the CEU (left panel) or YRI (right panel) datasets.

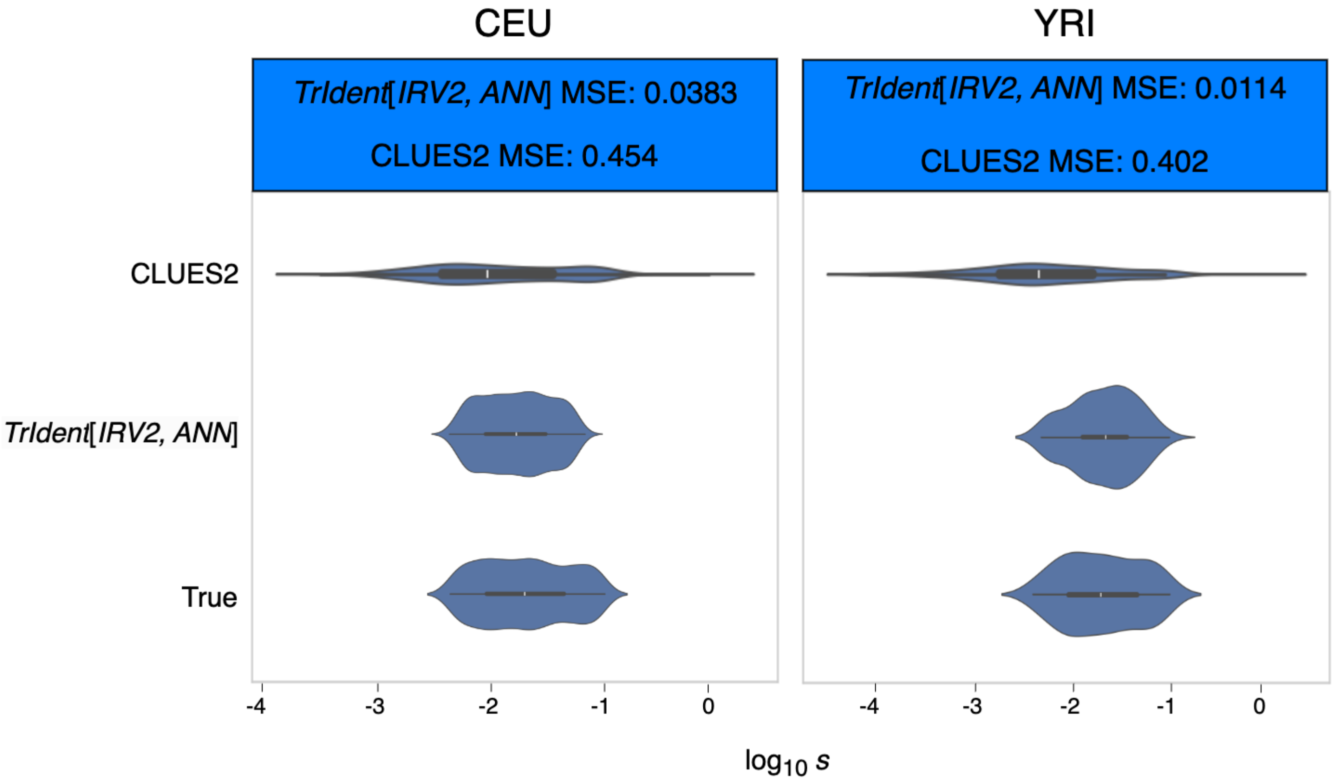


Figure S11: Summaries of distributions for true and predicted values of selection coefficient ($s$) computed using CLUES2 and the nonlinear *TrIdent*[*IRV2*, *ANN*] regression model. Distributions are summarized using violin plots with embedded box plots for the CEU (left) and YRI (right) datasets.

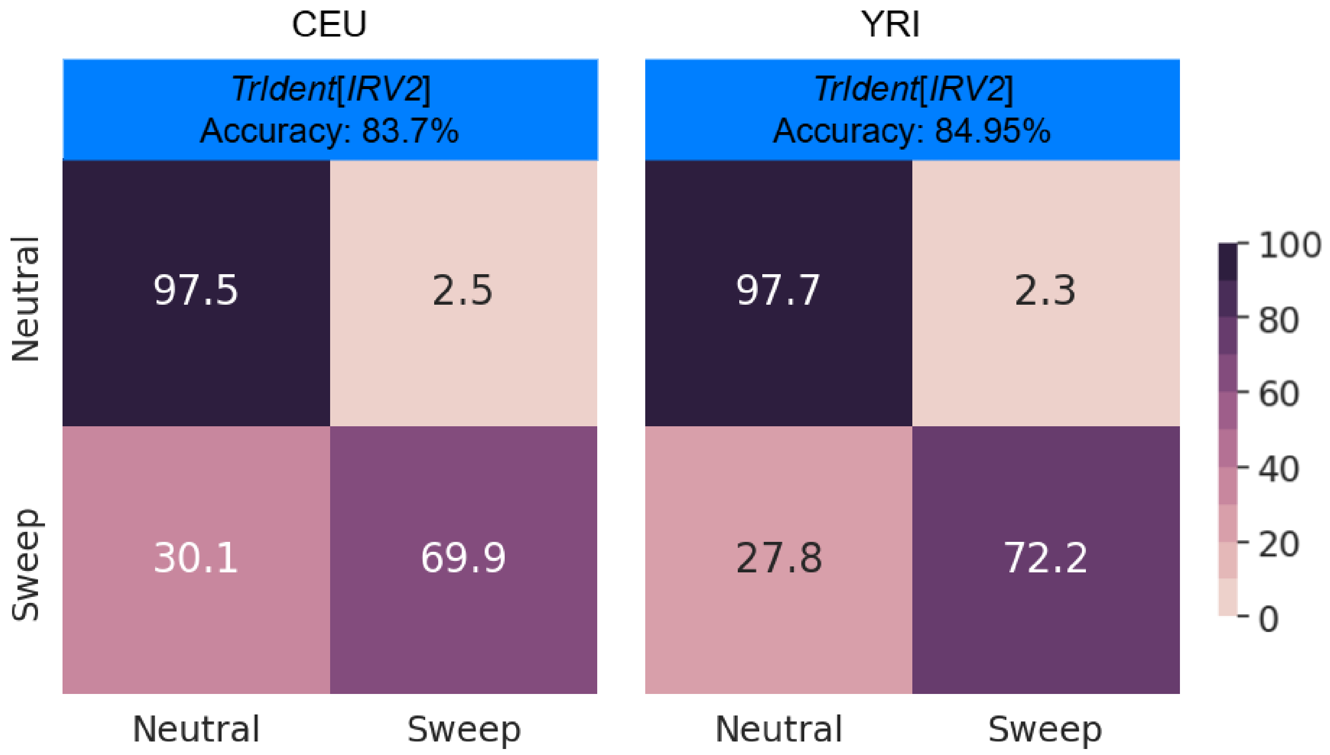


Figure S12: Classification rates and accuracies as depicted by confusion matrices to differentiate sweeps from neutrality on the CEU (left panel) and YRI (right panel) datasets for the *TrIdent*[*IRV2*] model when the sweep probability threshold is set at 0.9.

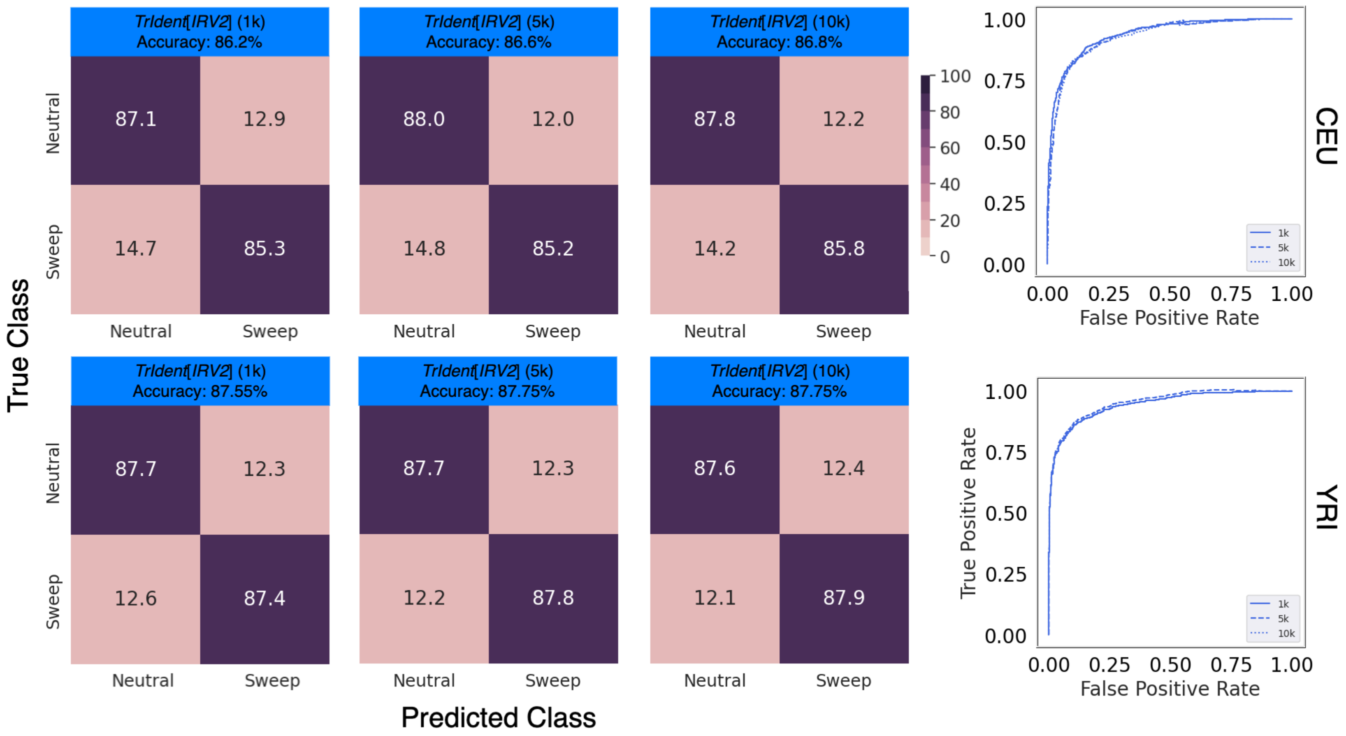


Figure S13: Classification rates and accuracies as depicted by confusion matrices and powers (true positive rates) to detect sweeps as depicted by receiver operating characteristic curves to differentiate sweeps from neutrality on the CEU (top panel) or YRI (bottom panel) test datasets for *TrIdent*[*IRV2*]. We compare test performance of *TrIdent*[*IRV2*] (1k), which is the original implementation of *TrIdent*[*IRV2*] trained with 1,000 samples per class, against *TrIdent*[*IRV2*] (5k) and *TrIdent*[*IRV2*] (10k) trained with 5,000 and 10,000 samples per class, respectively.

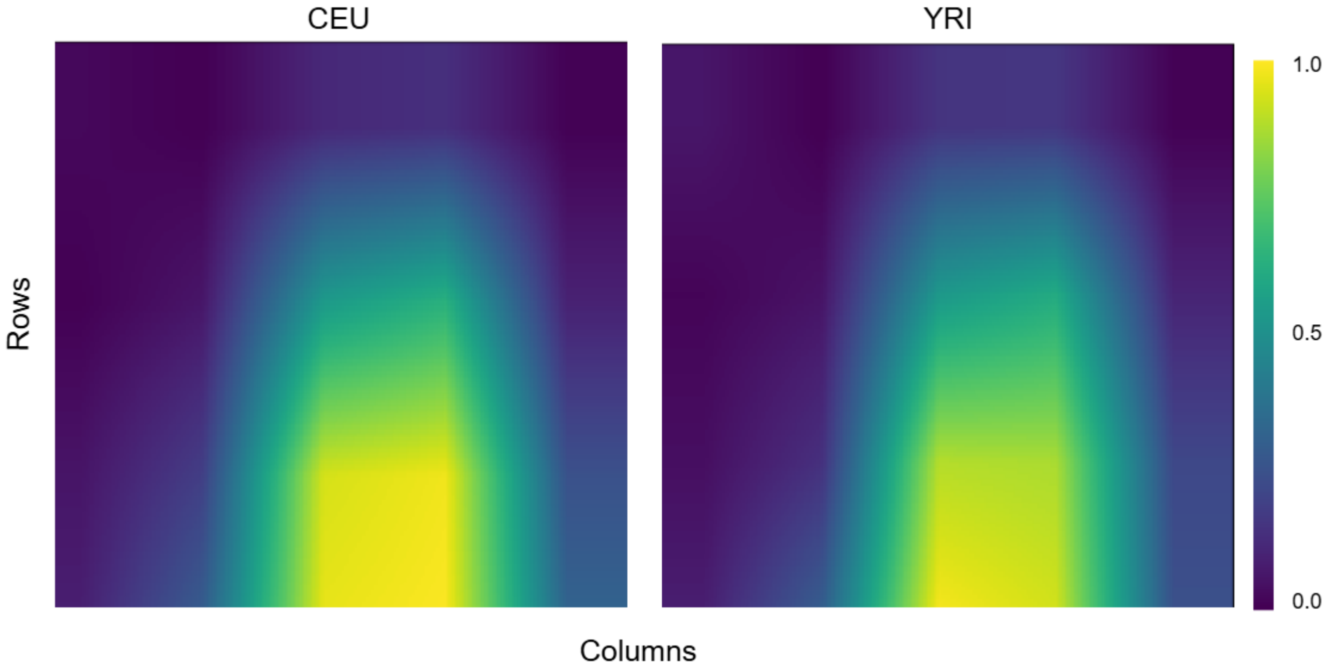


Figure S14: Heatmaps of mean gradient-weighted class activation maps (GradCAM) from *InceptionResNetV2* applied to CEU (left panel) and YRI (right panel) training data when training images are flipped horizontally, with the mean taken across 2,000 training observations with 1,000 observations from each of the neutral and sweep classes. Results show that the model still has a skew in emphasis toward the right of the central column, likely due to the redundancy of pixels on the left with those on the right accounted for through regularization during model fitting.

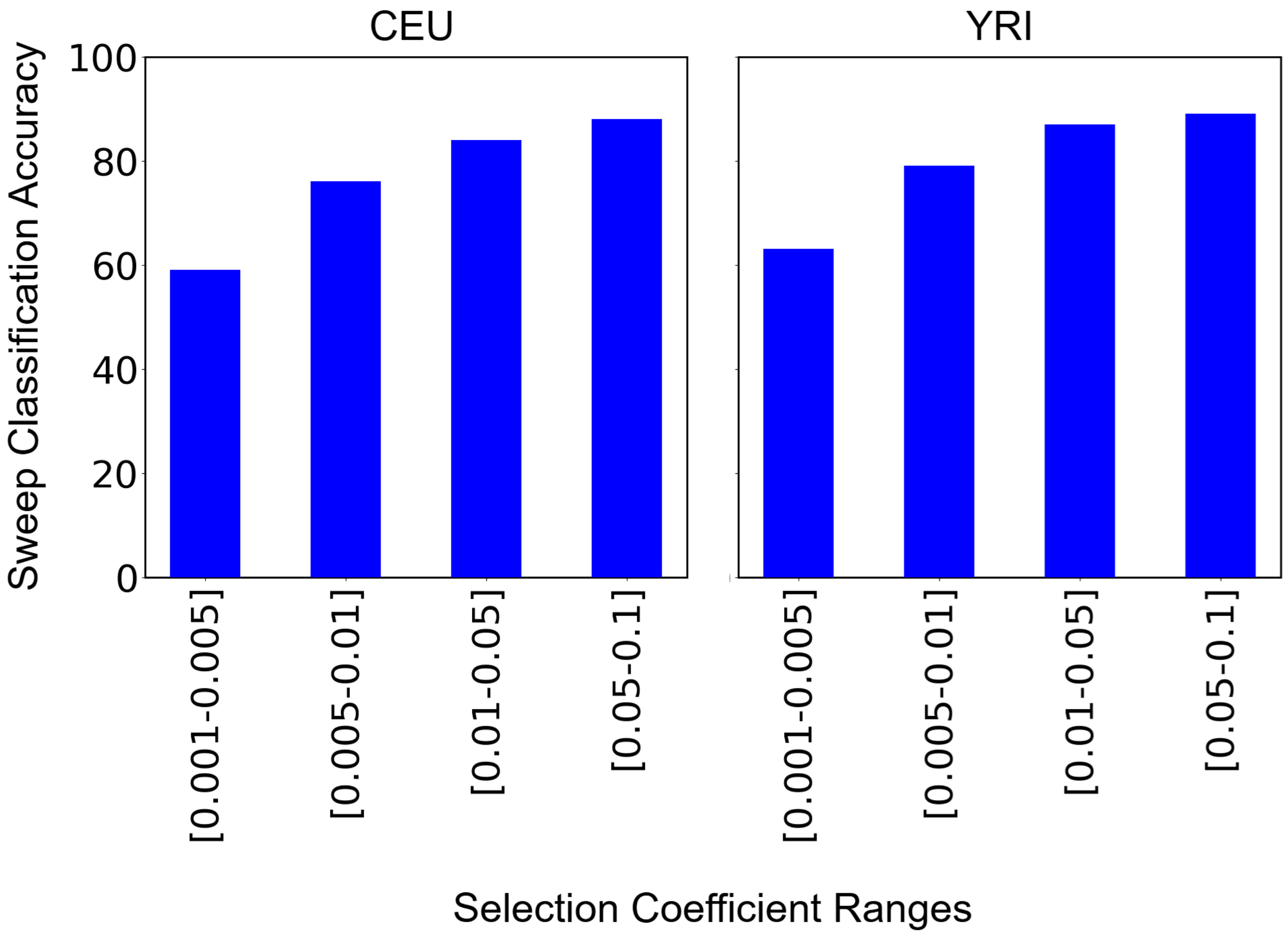


Figure S15: Sweep detection performance of *TrIdent*[*IRV2*] across different selection coefficient ranges. Sweep detection accuracy is shown for sweeps simulated with selection coefficients in the ranges $[0.001, 0.005]$, $[0.005, 0.01]$, $[0.01, 0.05]$, and $[0.05, 0.1]$ for the CEU test case (left panel) and the YRI test case (right panel).

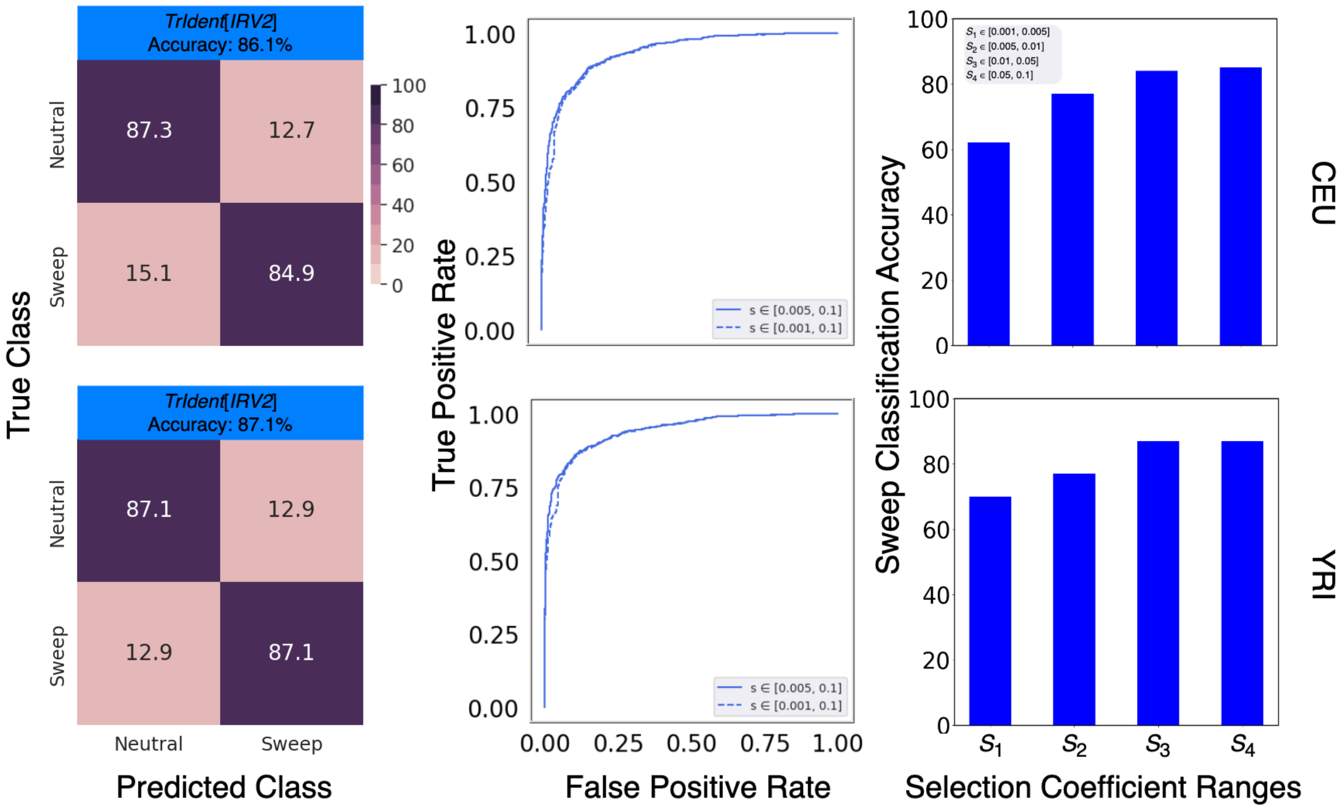


Figure S16: Classification rates and accuracies as depicted by confusion matrices (left panel) and powers (true positive rates; middle panel) to detect sweeps as depicted by receiver operating characteristic curves to differentiate sweeps from neutrality on the CEU (top panel) or YRI (bottom panel) test datasets for *TrIdent*[*IRV2*] when selection coefficients for training and test simulations were drawn from a distribution along the interval $[0.001, 0.1]$. Sweep detection accuracy (right panel) is shown for sweeps simulated with selection coefficients in the ranges $S_1 = [0.001, 0.005]$, $S_2 = [0.005, 0.01]$, $S_3 = [0.01, 0.05]$, and $S_4 = [0.05, 0.1]$ for the CEU test case and the YRI test case. A comparison against *TrIdent*[*IRV2*] trained on simulations with selection coefficients drawn from the interval $[0.005, 0.1]$ is provided in the middle panel.

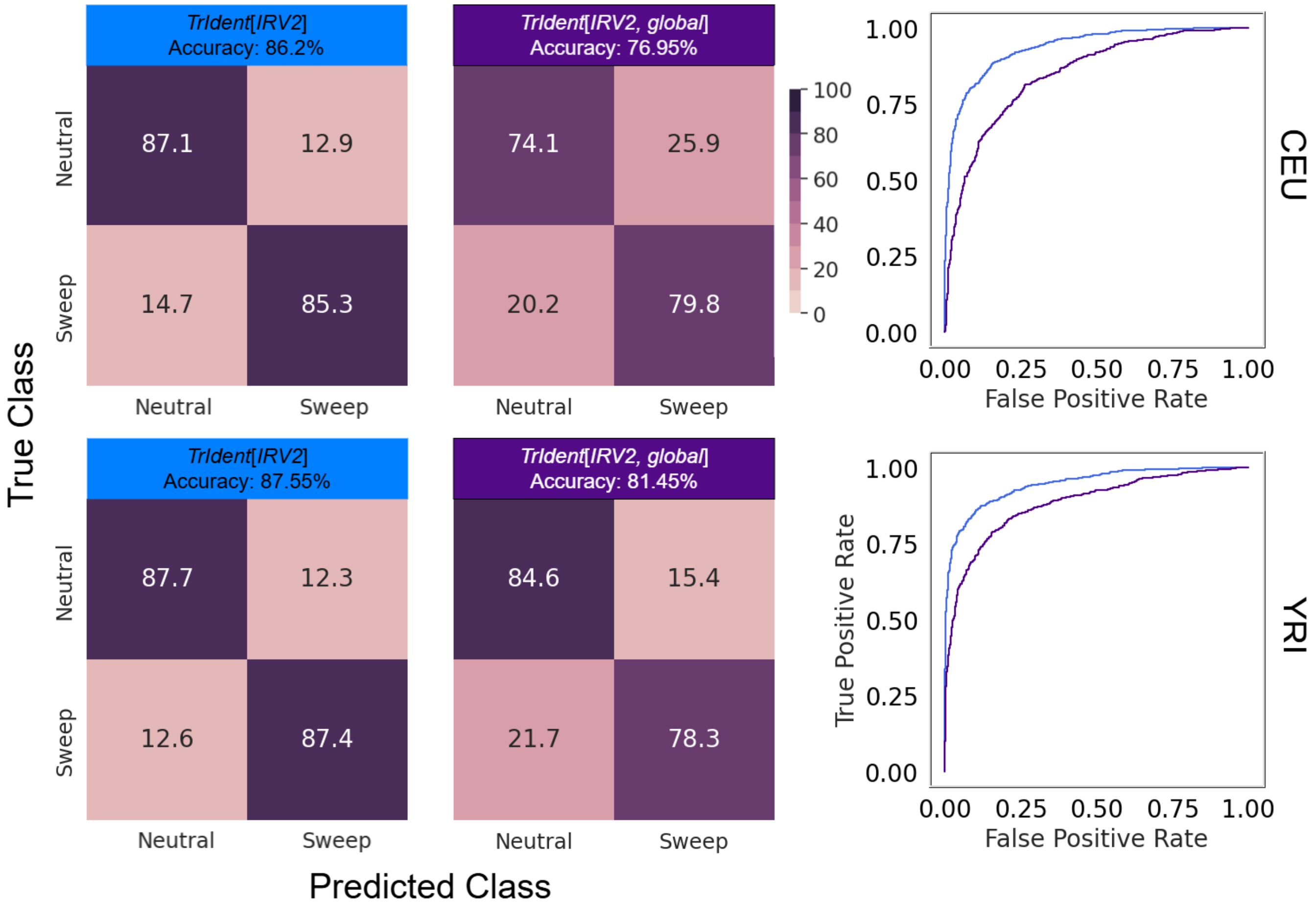


Figure S17: Classification rates and accuracies as depicted by confusion matrices and powers (true positive rates) to detect sweeps as depicted by receiver operating characteristic curves to differentiate sweeps from neutrality on the CEU (top panel) or YRI (bottom panel) test datasets for *TrIdent*[*IRV2*] and *TrIdent*[*IRV2*, *global*]. The images used as input to *TrIdent*[*IRV2*, *global*] are nearly identical to the process described in the *Image Generation* subsection of the *Methods*, with the exception that the 499 SNP window was sorted at once based on total minor allele counts.

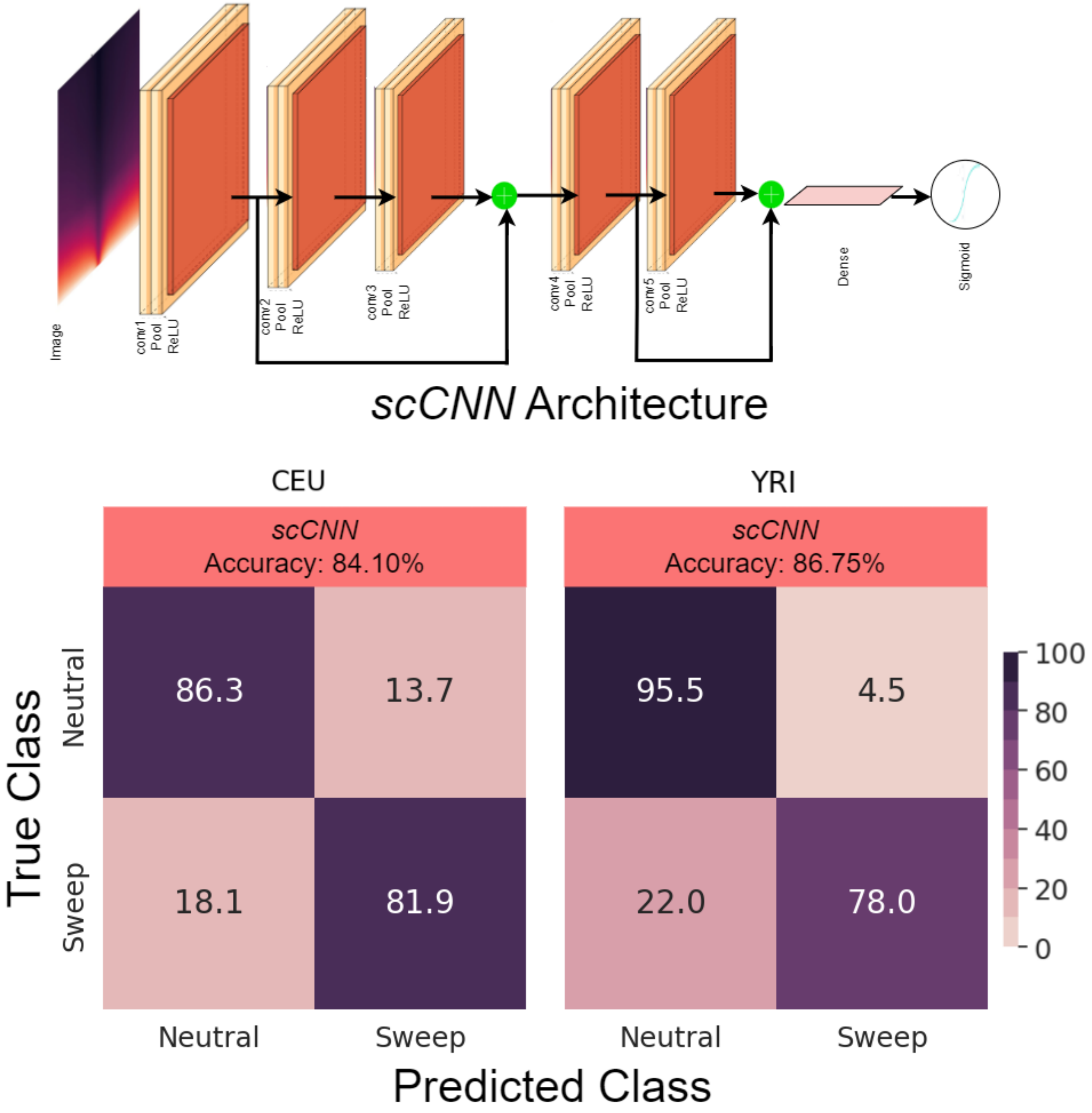


Figure S18: Classification results for a shallow yet complex architecture, termed *scCNN*, that is trained with *TrIdent* images (top panel). Classification rates and accuracies as depicted by confusion matrices to differentiate sweeps from neutrality on the CEU and YRI datasets for the *scCNN* model (bottom panel). Details of the *scCNN* model can be found in the *Discussion*.

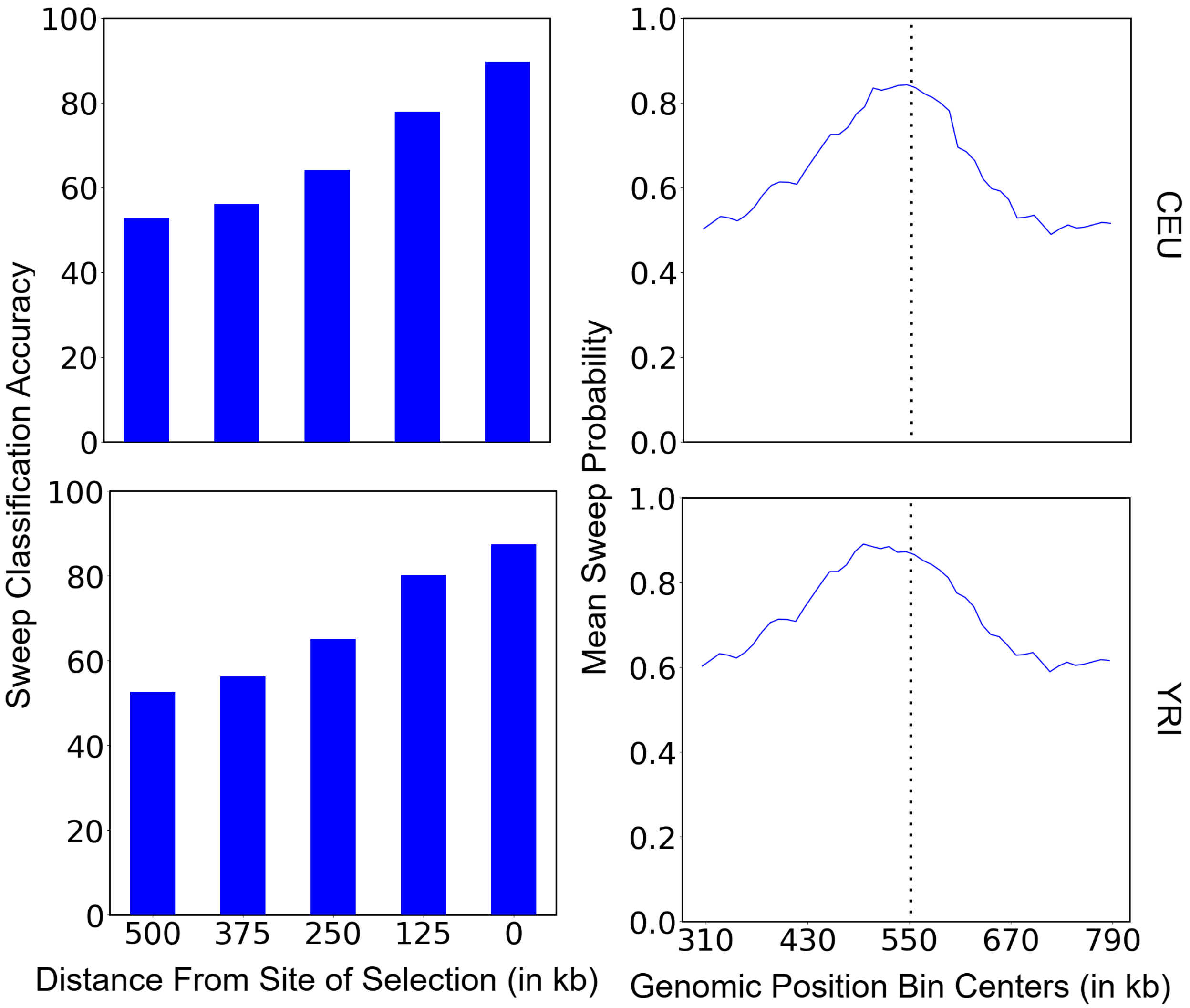


Figure S19: Sweep detection performance of *TrIdent*[*IRV2*] when the beneficial mutation is located at different distances from the image center for both CEU (top row) and YRI (bottom row) test cases. The left column presents sweep detection accuracy for sweeps positioned 500, 375, 250, 125, and zero kb away from the center of a 1.1 Mb simulated region, with *TrIdent* input images generated using the center SNPs at 550 kb. The right column displays the mean predicted sweep probability across 1,000 test images within overlapping 20 kb bins across the 1.1 Mb region, using a stride of 10 kb, illustrating how sweep probability signals diffuse with increasing distance from the beneficial mutation.

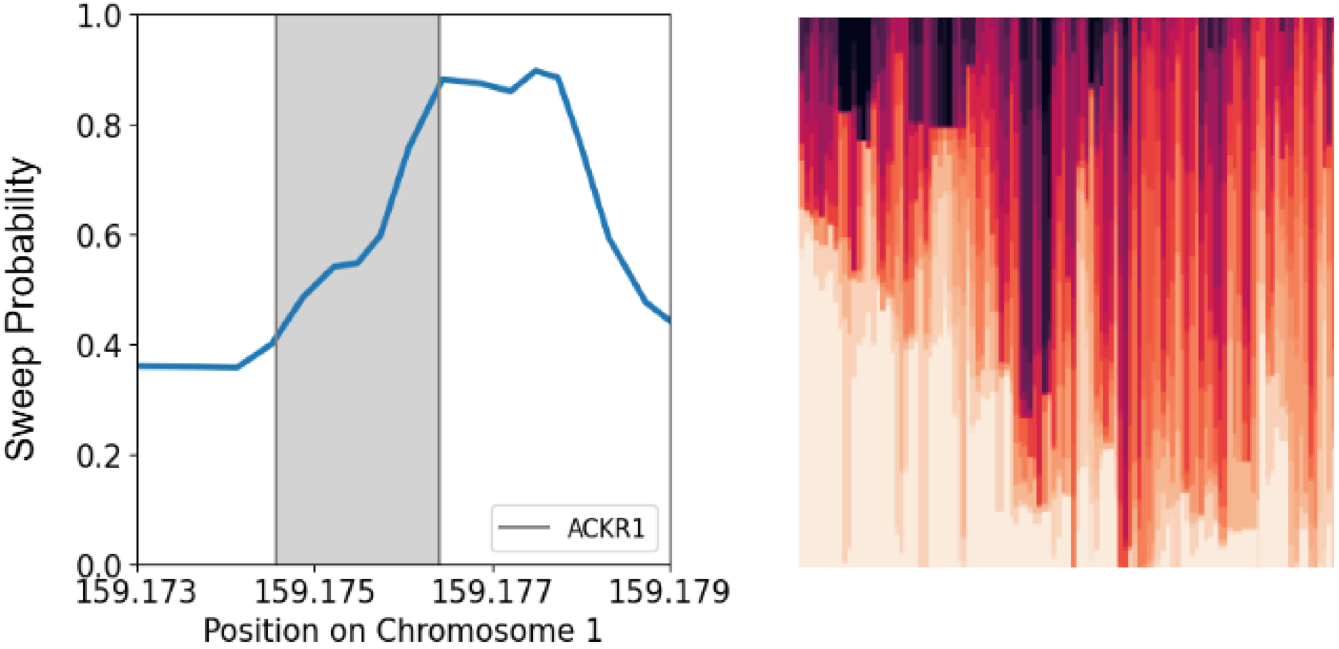


Figure S20: Identified candidate sweep region near the *ACKR1* (DARC) gene from the genome-wide scan produced using the trained *TrIdent*[*IRV2*] model on the sub-Saharan African (YRI) population in the 1000 Genomes Project dataset. Sweep probability as a function of chromosomal position (left) and visual representation of the haplotype variation surrounding the candidate region (right) are provided.

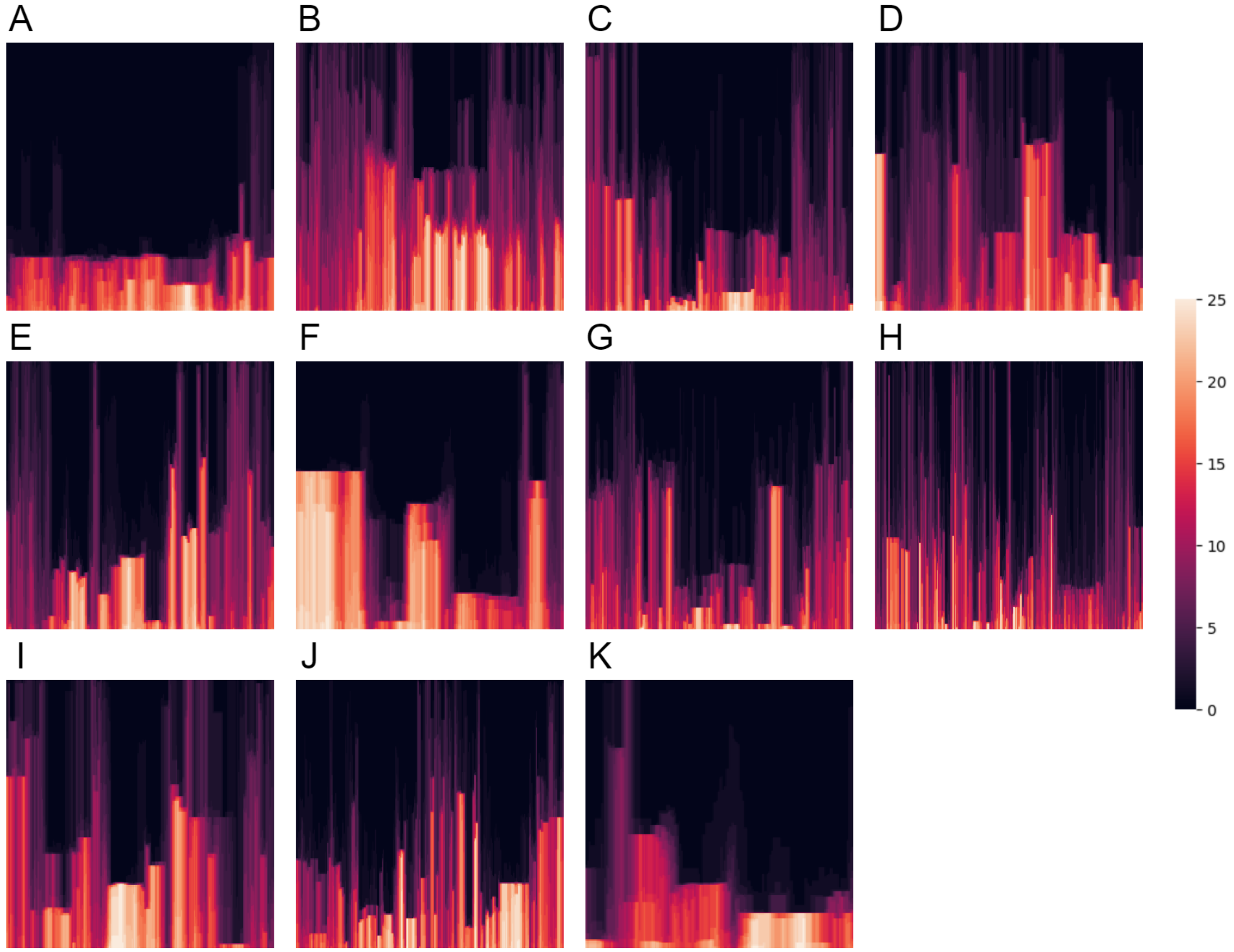


Figure S21: Images generated using the protocol outlined in the *Image Generation* subsection of the *Methods* for regions containing notable sweep candidates in the CEU population, as highlighted in Figure 9. However, unlike the standard procedure, all SNPs in the reported region of the corresponding panel of Figure 9 were utilized to create these images, which were then resized to $299 \times 299$.

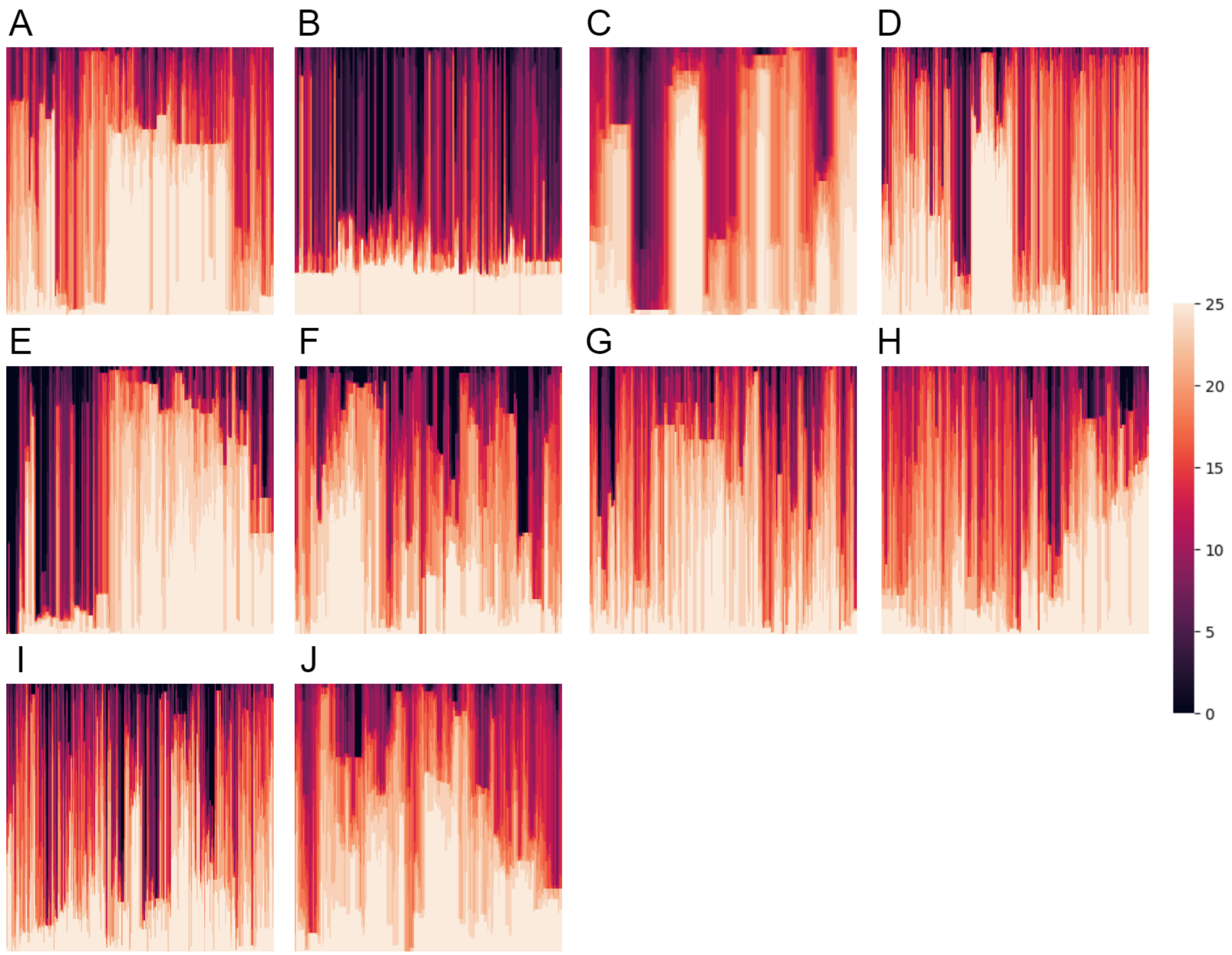


Figure S22: Images generated using the protocol outlined in the *Image Generation* subsection of the *Methods* for regions containing notable sweep candidates in the YRI population, as highlighted in Figure 10. However, unlike the standard procedure, all SNPs in the reported region of the corresponding panel of Figure 10 were utilized to create these images, which were then resized to $299 \times 299$.

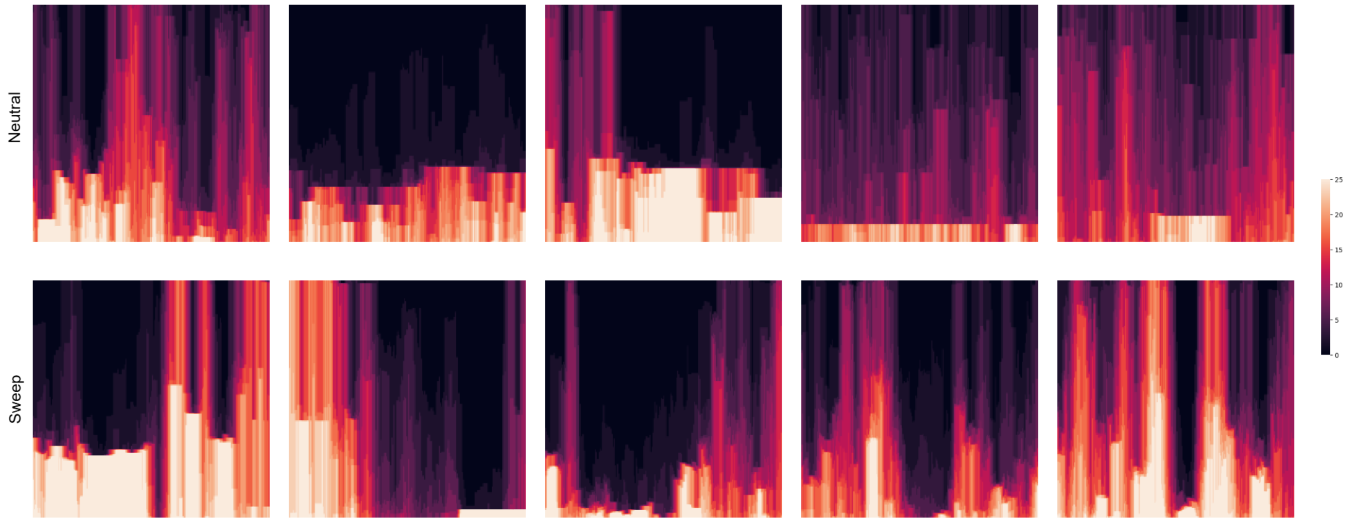


Figure S23: Images generated using the protocol outlined in the *Image Generation* subsection of the *Methods* for five test replicates with lowest sweep probability from the neutral test set and five test replicates with highest sweep probability from the sweep test set predicted by *TrIdent*[*IRV2*] under the CEU demographic history.

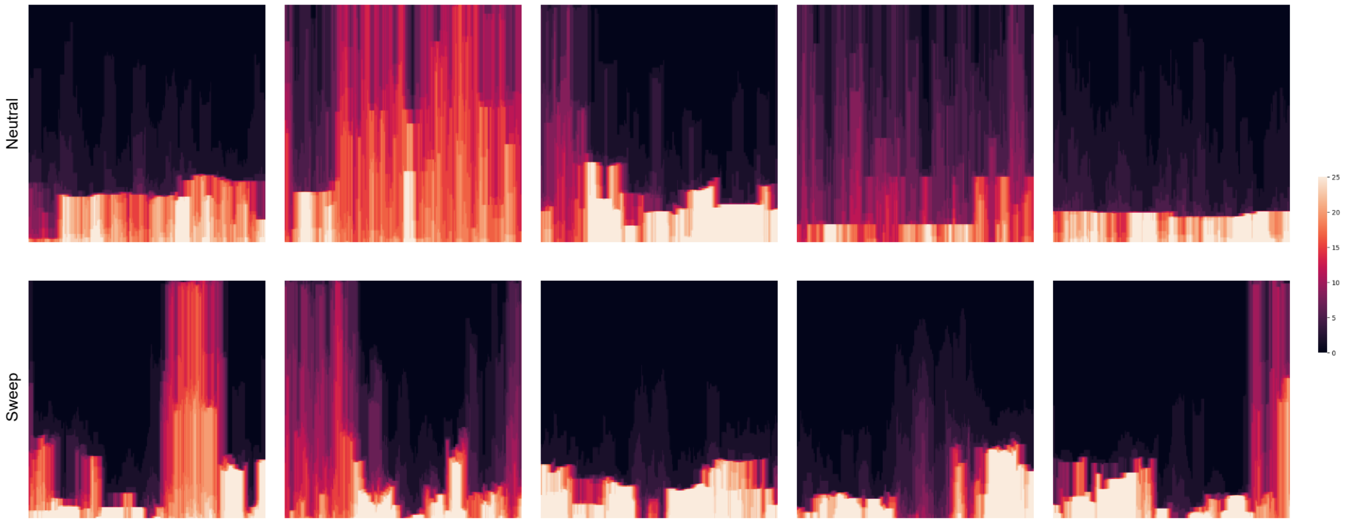


Figure S24: Images generated using the protocol outlined in the *Image Generation* subsection of the *Methods* for five test replicates with lowest sweep probability from the neutral test set and five test replicates with highest sweep probability from the sweep test set predicted by *TrIdent*[*IRV2*] under the YRI demographic history.
